# Leonardo: a toolset to correct sample-induced artifacts in light sheet microscopy images

**DOI:** 10.1101/2025.10.26.684661

**Authors:** Yu Liu, Gesine F. Müller, Lennart Kowitz, Tomáš Chobola, Kurt Weiss, Paul Maier, Jie Luo, Malte Roeßing, Martin Stenzel, Anika Grüneboom, Johannes Paetzold, Ali Ertürk, Nassir Navab, Carsten Marr, Jianxun Chen, Jan Huisken, Tingying Peng

## Abstract

Selective plane illumination microscopy (SPIM, also known as light sheet fluorescence microscopy) is the method of choice for studying morphogenesis and function in biological specimens over extended periods, as it permits gentle and rapid volumetric imaging. In inhomogeneous samples, however, sample-induced artifacts, including light absorption, scattering, and refraction, can impact the image quality, particularly as the focal plane gets deeper into the sample. Here, we present *Leonardo*, the first toolbox designed to address the major sample-induced artifacts by using two modules: (1) *DeStripe* removes stripe artifacts in SPIM caused by light absorption while preserving fine sample structures; (2) *Fuse* reconstructs a single high-quality image from dual-sided illumination and/or dual-sided detection, while eliminating blur and optical distortions caused by light scattering and refraction. The efficacy of Leonardo is validated on a wide range of biological samples, from minimally invasive experiments on sensitive specimens (translucent embryonic and optically opaque larval zebrafish) to cleared mouse samples up to two centimeters in size. We provide model code and a Napari-based graphical user interface, enabling the SPIM community to easily apply Leonardo to advance light sheet imaging of inhomogeneous and complex specimens.

## Introduction

More than ever, biologists rely on advanced imaging techniques to study large, ideally living, specimens in three dimensions (3D)^1,2^. In response to this demand, selective plane illumination microscopy (SPIM), also known as light sheet fluorescence microscopy (LSFM), has emerged as a powerful technique^3,4^. It uses two orthogonal optical paths (Fig. 1A): one to illuminate the specimen with a thin light sheet and the other to detect the emitted fluorescence. This sidewise illumination of a thin volume is particularly beneficial when imaging living specimens, as it minimizes photodamage, photobleaching, and out-of-focus fluorescence background^5^. Moreover, the detection of fluorescent signals in SPIM can be remarkably rapid, as it captures the entire field of view in a single exposure using fast and sensitive cameras^5,6^.

**Fig. 1.**
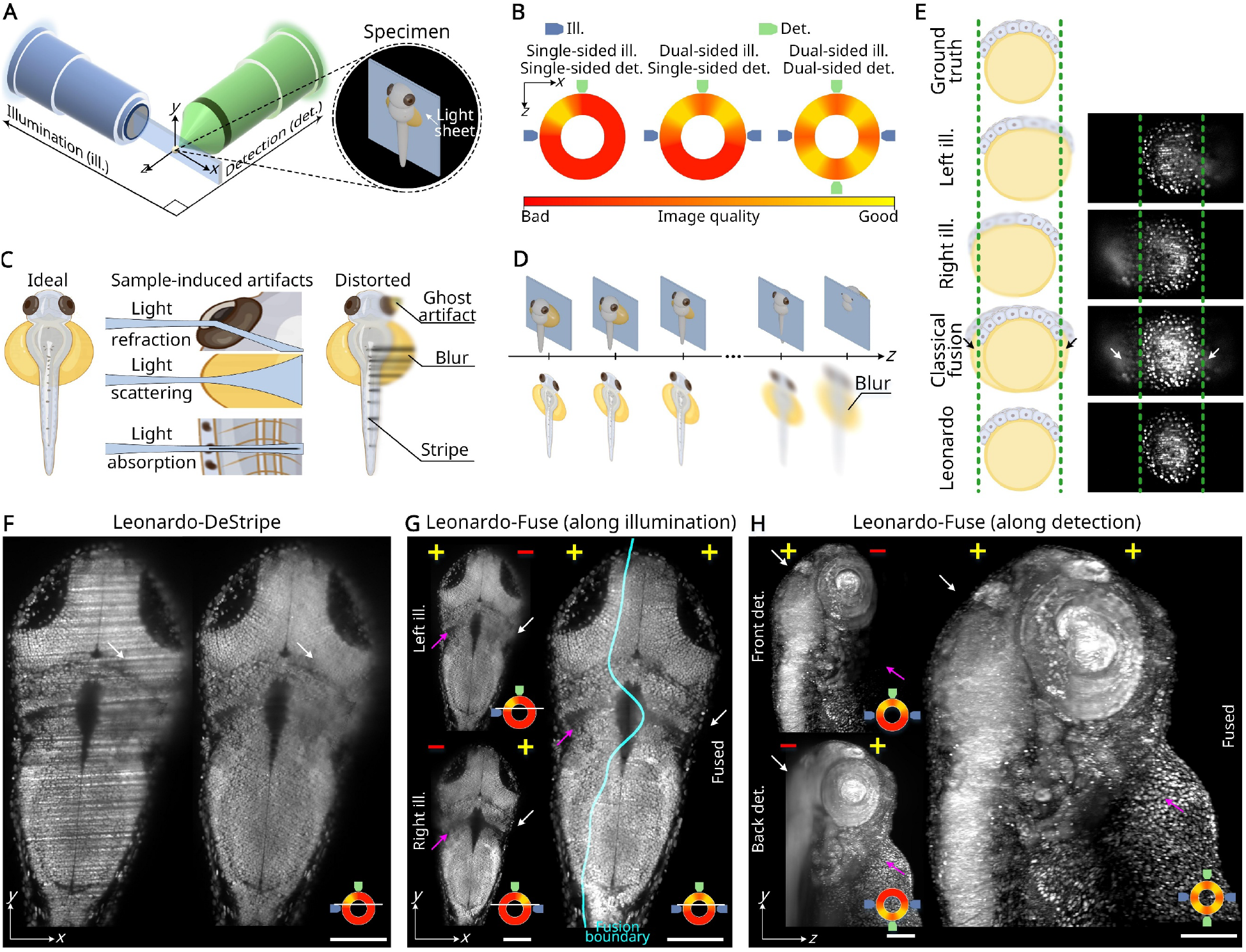
Leonardo is an AI-powered toolkit for reconstructing light sheet datasets with reduced sample-induced artifacts. (A) SPIM features illumination from the side and may be vulnerable to artifacts caused by the optical inhomogeneity of the specimen. (B) Multi-view imaging, i.e., dual-sided illumination and/or dual-sided detection, is widely used in SPIM to increase the sample coverage. (C) Common sample-induced artifacts include light absorption, scattering, and refraction, which lead to stripes, blur, and ghost artifacts, respectively. (D) Additionally, due to limited penetration depth along the detection axis, images from deeper layers of the specimen become increasingly blurry and lose contrast. (E) Schematic illustration showing that ghost artifacts render the far side of the image—away from the illumination source—visually larger than the ground truth (beyond the dotted lines). These ghost artifacts are often retained in conventional fusion (white and black arrows) but are effectively suppressed by Leonardo. Leonardo operates in three key steps: (F) Leonardo-DeStripe, a deep learning-empowered method that can resolve stripes (white arrows); (G) Leonardo-Fuse (along illumination), which fuses datasets illuminated from opposing orientations. It estimates a fusion boundary (cyan curve) that selects high-quality sub-stacks which are sharp (magenta arrows) and free of ghost artifacts (white arrows); (H) Leonardo-Fuse (along detection), which fuses datasets detected from opposing views, excluding regions that are blurry (magenta arrows) and contain ghost artifacts (white arrows). Scale bars: 100 μm.

As with any light microscope, interactions between the light and the specimen in SPIM lead to image quality degradation, particularly when imaging large, optically inhomogeneous samples^7^. Here, we distinguish three types of sample-induced artifacts that often occur in parallel: light absorption, scattering, and refraction. Specifically, along the excitation path, denser or absorbing structures can attenuate the light sheet, resulting in non-uniform illumination and stripy shadows^8,9^ (Fig. 1C). Optical approaches such as using multiple static light sheets or pivoting a single one during exposure— known as multidirectional SPIM (mSPIM)^10^—can help reduce such stripes. However, the range of illumination angles in these methods is typically limited to ±10°and therefore not able to completely eliminate stripes when imaging complex samples^8,9^. To address these limitations, several software-based methods, such as the multidirectional stripe remover (MDSR)^11^, variational stationary noise-removing algorithms (VSNR)^12^, and a more recent variational destriping method proposed by Rottmayer *et al*.^13^, have been developed to computationally suppress stripe artifacts based on their directionality. Yet, these approaches often lack robustness to sample-specific variations in stripe appearance and may inadvertently remove genuine sample features, limiting their general applicability^14^.

In addition to light absorption, light scattering and refraction also degrade image quality^15^. Light scattering causes the light sheet to widen, leading to a progressive decrease in sectioning quality and an increase in blur as the light sheet propagates deeper from its entry site (Fig. 1C). Meanwhile, light refraction, resulting from refractive index mismatches between the sample and the medium or variations within the specimen, can misposition the light sheet and cause optical distortions at the distal side, leading to ghost artifacts (Fig. 1C). Unlike blur caused by light scattering, ghost artifacts occur in regions where no signal should be present—rendering the sample visually bigger—and often resemble real structures in shape and intensity (Fig. 1E). Similarly, light scattering and refraction also degrade emitted fluorescence on the detection side, leading to a loss of contrast in the deeper layers of the specimen (Fig. 1D). As a result, only the portion of the specimen facing both the illumination and the detection lenses can be imaged with optimal clarity.

To mitigate some of these problems, dual-sided illumination has proven effective by sequentially illuminating the sample from opposite orientations with a second illumination lens^10^ (Fig. 1B). Here, sequential illumination—rather than simultaneous illumination—with two counter-propagating beams is preferred to avoid beam overlap, which would otherwise result in thicker optical sections and reduced image quality.

Additionally, sample rotation by 180°or dual-sided detection with a second detection arm opposite the first one can further improve sample coverage^10,16^ (Fig. 1B). The goal of subsequent image fusion is then to reconstruct a single volume that inherits all high-resolution information from individual stacks, thereby expanding sample coverage^15,17^. Unfortunately, existing local-based fusion algorithms, such as BigStitcher^18^ and Huygens^19^, aim to preserve all structured information from individual stacks and thereby struggle to distinguish and remove structure-like ghosts.

We introduce *Leonardo*, a pioneering set of AI-based image processing tools specifically designed to enhance selective plan<u>e</u> illumination micr<u>o</u>scopy by reco<u>n</u>structing d<u>a</u>tasets with reduced sample-in<u>d</u>uced artifacts. First, we introduce *Leonardo-DeStripe*, which leverages a graph neural network (GNN) to parameterize a two-dimensional (2D) band-pass filter in the Fourier space, effectively removing shadow defects caused by light absorption within tissues while retaining structural integrity (Fig. 1F, Extended Data Fig. 1). Second, we propose *Leonardo-Fuse*, which efficiently merges high-quality information from dual-sided illumination (Fig. 1G) and/or dual-sided detection (Fig. 1H) using two submodules, respectively. A highlight of Leonardo-Fuse is its joint consideration of local image qualities, geometrical shape of the specimen, and prior knowledge of the sample’s orientations with respect to the illumination, making it effective in eliminating ghost artifacts caused by light refraction (Fig. 1E). In addition to its methodological innovation, Leonardo offers a user-friendly, Napari-based graphical interface for intuitive, interactive operation, while its open-source codebase is designed to encourage customization by advanced users (Supplementary Fig. 1). We demonstrate the effectiveness and adaptability of Leonardo across a wide range of samples acquired with different SPIM variants, including home-built microscopes and the commercial Ultramicroscopy Blaze.

## Results

### Leonardo-DeStripe removes stripe artifacts in unidirectional SPIM

Stripy shadows, caused by partial light absorption of the illumination light by dense tissue, occur in all light microscopes but are particularly pronounced in the case of SPIM due to its sidewise, mostly parallel illumination (Fig. 1A, C). Existing stripe removal methods generally fall into two categories (Supplementary Note 1): (1) Fourier-domain filtering, such as band-pass filters applied in a transformed domain like contourlet^11^ or wavelet^20^, which exploit the fact that elongated stripe artifacts mainly occupy only a narrow frequency band perpendicular to the stripe direction; and (2) spatial-domain optimization approaches, typically based on priors like anisotropic total variation^13,21^, assuming high intensity variation orthogonal but low variation parallel to the stripe direction. While the Fourier-based approaches unavoidably remove genuine structures that share similar frequencies with the stripe artifacts (Supplementary Fig. 2), optimization-based approaches also struggle to suppress stripes without compromising structural fidelity due to the ill-posed nature of the inverse problem (Supplementary Figs. 3-4, Extended Data Fig. 1).

Leonardo-DeStripe overcomes these limitations by combining the strengths of both paradigms: it uses a graph neural network (GNN) to inpaint each corrupted Fourier coefficient within a wedge-shaped mask oriented perpendicular to the stripe direction by aggregating neighboring coefficients in a polar coordinate system in Fourier space (Fig. 2A, Extended Data Fig. 2D). These weights are learned in a self-supervised fashion, with the loss computed on the reconstructed image after inverse Fourier transform (Methods). Operating in the Fourier domain enables Leonardo-DeStripe to work with a sparse, compact representation of stripe-corrupted data, improving inpainting efficiency and stability. To further improve computational efficiency, the GNN is trained in a downsampled space, with a guided upsampling strategy to restore full resolution (Methods, Supplementary Fig. 5). Particularly, to ensure that true sample structures remain intact after stripe removal, we introduce a dedicated post-processing module by incorporating the directional illumination prior in SPIM (denoted as ill. prior, Methods).

**Fig. 2.**
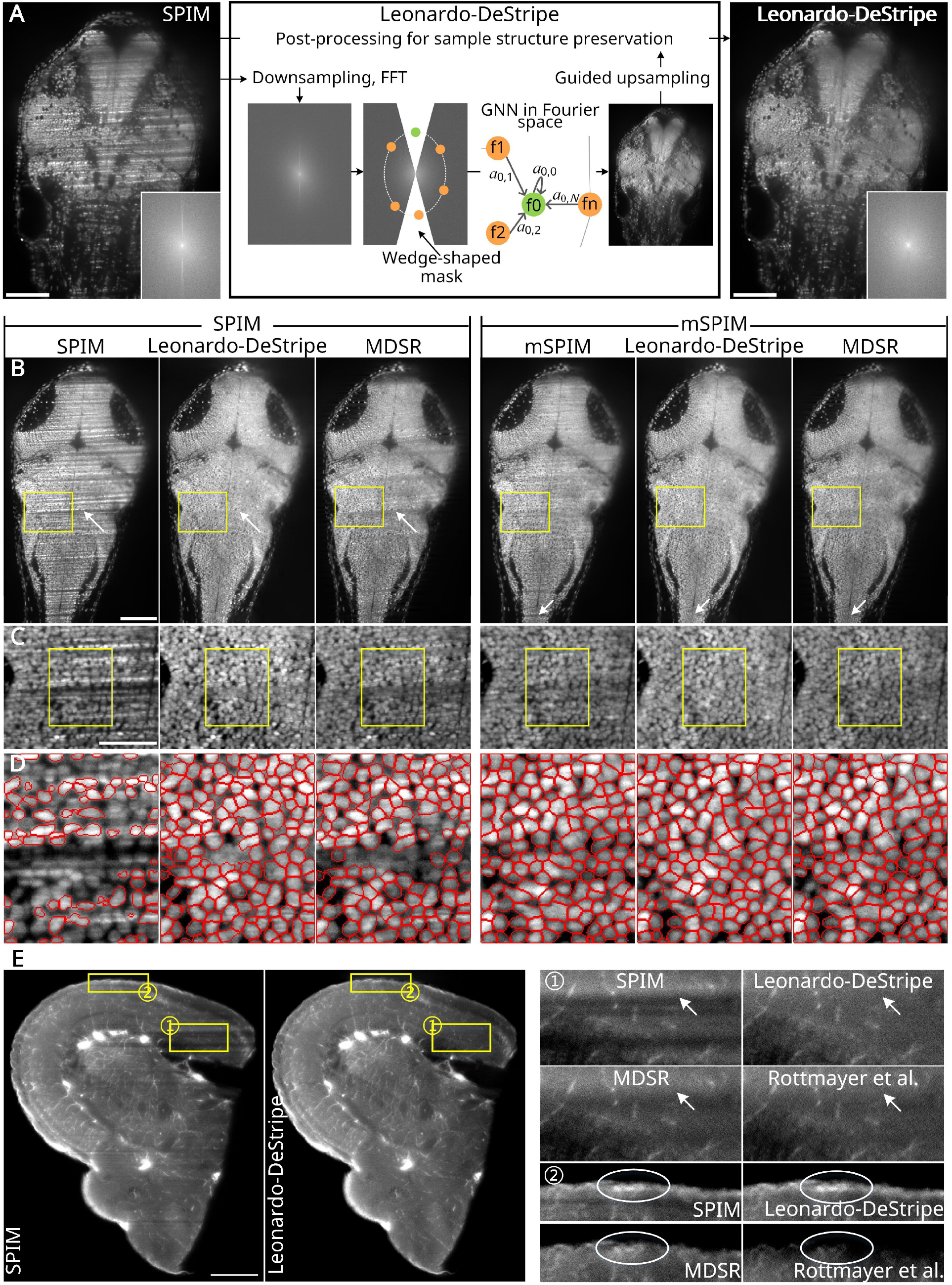
Leonardo-DeStripe features parameterizing a Fourier-based image restoration model with deep learning. (A) Leonardo-DeStripe performs stripe removal in Fourier space by inpainting stripe-corrupted Fourier coefficients as a combination of their neighbors on a polar coordinate. Weights in terms of neighbor summation are learned using a graph neural network (GNN). (B) A detailed comparison on a H2B-GFP zebrafish, between Leonardo-DeStripe and MDSR, on both SPIM and mSPIM datasets. Leonardo eliminated stripes better (white arrows), especially on SPIM data that was heavily affected by stripes in (C). (D) Stripe removal helped downstream processing, e.g., cell segmentation using Cellpose. Cellpose segmentation after Leonardo-DeStripe produced more continuous cell boundaries compared to MDSR-enhanced data, whereas segmentation largely failed on the unprocessed SPIM image. (E) Leonardo-DeStripe successfully removed occasionally very thick stripes on a zebrafish brain dataset, where both comparison methods, MDSR and Rottemayer et al., left residual stripes (arrows in region ➀). Notably, comparison methods removed genuine sample structures in zoom-in regions ➁ (circles), where Leonardo-DeStripe preserved them. Scale bars: 100 μm (A, B), 50 μm (C), 25 μm (D), 200 μm (E).

This allows Leonardo-DeStripe to differentiate between stripe artifacts and structurally similar sample structures, even when sample structures are oriented along the stripe direction (Extended Data Fig. 1, Supplementary Table 1).

We first evaluated Leonardo-DeStripe on synthetic stripes (Supplementary Fig. 6, Supplementary Note 4) that were simulated on adipose tissue^2^. Leonardo-DeStripe was benchmarked against the multidirectional stripe remover (MDSR)^11^ and Rottmayer et al.^13^, representing state-of-the-art methods from the Fourier-domain filtering or spatial-domain optimization categories, respectively. Leonardo-DeStripe left fewer residual stripes when compared with MDSR and Rottmayer et al. (Extended Data Fig. 4A-B). The superior performance of Leonardo-DeStripe over the two baseline methods was also confirmed by quantitative results, where Leonardo-DeStripe achieved a higher peak signal-to-noise ratio (PSNR) (35.4 vs. 29.3/26.9) and structural similarity index (SSIM) (0.97 vs. 0.93/0.92) (Extended Data Fig. 4C, Supplementary Note 6).

Next, we evaluated Leonardo-DeStripe on real stripe-corrupted data, e.g., a live zebrafish embryo at 3 days post-fertilization (Methods). Here, we compared Leonardo-DeStripe and both baselines against mSPIM, the optical imaging solution, which mitigates stripes by pivoting the light sheet in the detection focal plane^10^ (Fig. 2B, Supplementary Fig. 7). Although mSPIM, MDSR and Rottmayer et al. all efficiently reduced thin stripes, residual stripes, particularly thick ones, were still visible in the images (white arrows in Fig. 2B, white arrows in Supplementary Fig. 7). In contrast, Leonardo-DeStripe successfully resolved all stripes with varying thickness (Fig. 2C, Supplementary Fig. 7). Moreover, Leonardo-DeStripe can be applied in conjunction with mSPIM, effectively removing residual stripes even after optical correction.

We assessed the impact of stripe removal on downstream image analysis, specifically cell segmentation, using Cellpose^22^. On SPIM images, Leonardo-DeStripe significantly enhanced segmentation results by effectively removing stripes, with segmentation results from mSPIM serving as a pseudo-ground truth (Fig. 2D). Importantly, Leonardo-DeStripe did not alter segmentation results on mSPIM images, indicating that it preserved original sample information while removing artifacts (Supplementary Video 6). Furthermore, Leonardo-DeStripe generalized well across different specimens with varying stripe characteristics, including periodic thin stripes and aperiodic thick stripes, in large and diverse samples such as fixed murine heart tissue (Supplementary Fig. 8, Supplementary Video 1) and zebrafish brain tissue (Fig. 2E, Supplementary Video 2). Notably, Leonardo-DeStripe significantly outperformed MDSR in removing challenging thick stripes (patch ➀, Fig. 2E), which MDSR failed to eliminate. Although the method by Rottmayer et al. achieved better stripe removal than MDSR, it erroneously eliminated genuine sample structures (patches ➁, Fig. 2E), compromising biological fidelity. By contrast, Leonardo-DeStripe effectively resolved both thin and thick stripes without sacrificing authentic sample features, owing to its dedicated post-processing module designed for structure preservation.

### Leonardo-DeStripe generalizes well on multi-directional SPIM

The UltraMicroscope Blaze, a commercial SPIM-based system, uses two sets of three light sheets from both sides, positioned with a 10°offset, in addition to the normal light sheet. This setup results in complex diamond-shaped stripe patterns along multiple directions (Fig. 3A). We extended Leonardo-DeStripe to efficiently remove stripes also in such multi-directional SPIM systems.

**Fig. 3.**
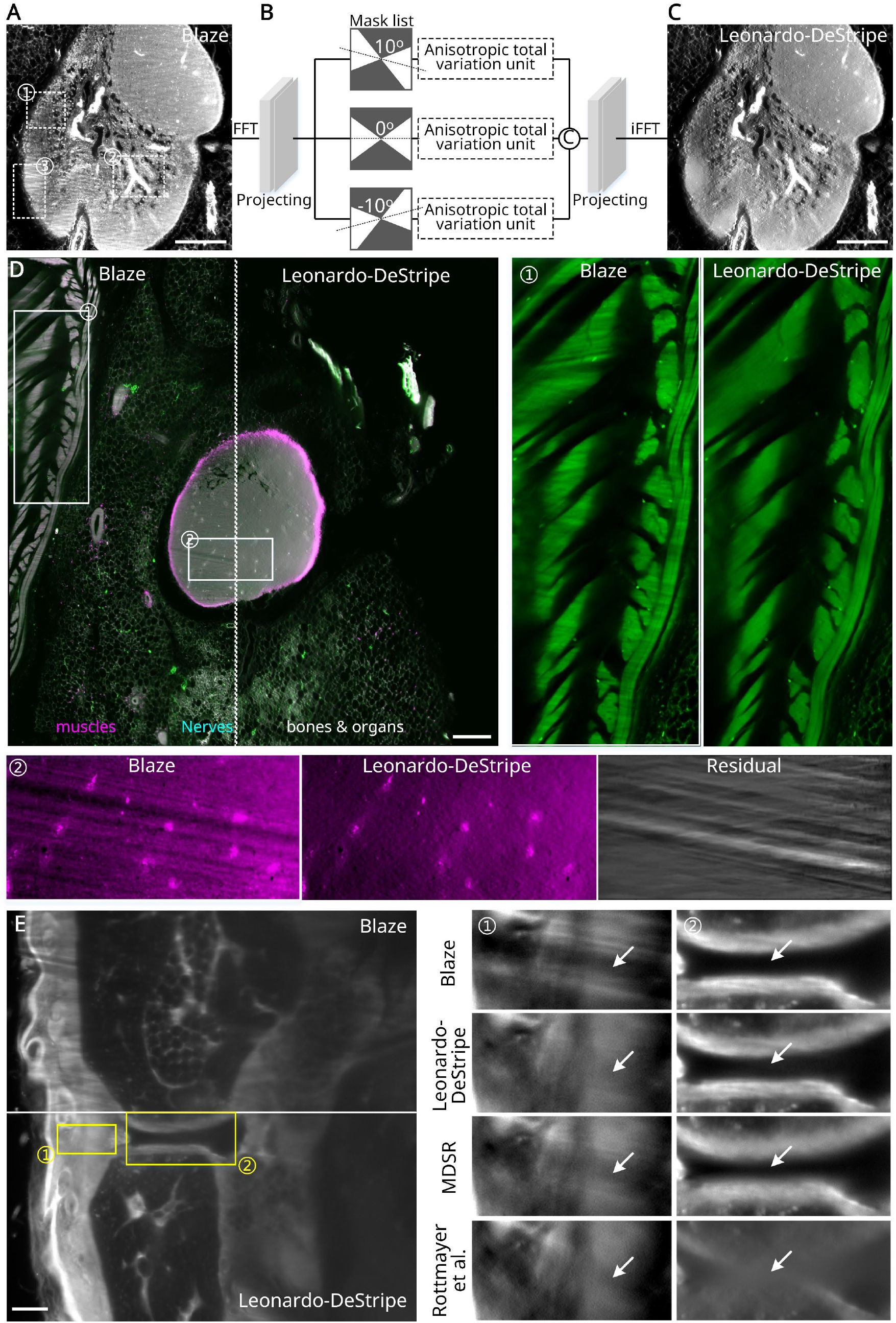
Leonardo-DeStripe excels at handling multi-directional SPIM stripe artifacts. (A-C) By defining different masks oriented in diverse directions in Fourier space and reusing anisotropic total variation units, Leonardo-DeStripe is extensible to multi-directional SPIM. For example, in UltraMicroscope Blaze imaging, the sample is illuminated along three directions (0°, 10°, and -10°). (D) A whole view of a mouse specimen with antibodies against the TH antibody (green) and the immune cell marker CD45 (magenta). Despite varying stripe characteristics within the tissue—such as periodic stripes in one region (patch ➀) and thicker stripes in another (patch ➁)—Leonardo-DeStripe successfully removed all stripes without erasing structural information, as verified by the residual image. (E) A murine hind leg dataset, where only Leonardo-DeStripe removed stripes (white arrows in patches ➀) without distorting sample structures (white arrows in patches ➁). Scale bars: 200 μm (A, C, D), 500 μm (E).

To address stripes in arbitrary unidirectional orientation, one could rotate the input image to align the stripes horizontally and apply Leonardo-DeStripe multiple times, once for each orientation. However, this will triple the processing time, and is computationally inefficient. Instead, we implemented three rotated wedge-shaped masks in the Fourier space, coupled with three anisotropic total variation regularization units for structure preservation of the graph neural network (Methods, Fig. 3B, Supplementary Fig. 9). The learned feature maps could then be concatenated, merged channel-wise in a deep learning fashion, and finally fed into the subsequent operators in the standard Leonardo-DeStripe pipeline (Supplementary Note 2). As a result, Leonardo-DeStripe was able to remove stripes in all orientations in a single step (Fig. 3C).

We benchmarked the performance of Leonardo-DeStripe on multi-directional stripes against MDSR and Rottmayer et al. In a mouse stained with antibodies against the TH antibody (green) and the immune cell marker CD45 (magenta)^2^, all methods removed stripes while preserving sample structures well in regions with fewer stripe artifacts (patch ➀, Extended Data Fig. 3C). However, in areas with more pronounced stripes, Leonardo-DeStripe consistently removed stripes while preserving sample structures, whereas MDSR either left residual stripes (patch ➂, Extended Data Fig. 3C) or introduced artifacts that severely compromised sample information (patch ➁, Extended Data Fig. 3C). A broader and qualitative comparison of the multi-tile data slice before and after applying Leonardo-DeStripe is provided in Fig. 3D. Despite the varying stripe characteristics within the tissue—such as periodic stripes in one region (patch ➀, Fig. 3D) and thicker stripes in another (patch ➁, Fig. 3D)—Leonardo-DeStripe effectively resolved all of them. Additionally, we benchmarked Leonardo-DeStripe against MDSR and the method by Rottmayer et al. using a murine hind leg dataset, where critical sample structures were aligned with the stripe direction (e.g., horizontal). In this sample, the two baseline methods either failed to completely remove stripes (white arrows in patch ➀, Fig. 3E) or heavily suppressed genuine structures (white arrows in patch ➁, Fig. 3E). By contrast, Leonardo-DeStripe successfully preserved aligned structures while effectively removing the stripe artifacts.

### Leonardo-Fuse produces highly contrasted fusion results with minimal optical distortions along the illumination

Beyond light absorption, sample-induced light scattering and refraction further deteriorate image contrast at the distal side of the light sheet. Dual-sided illumination (illuminating the specimen from opposite sides sequentially, T-SPIM) helps mitigate light falloff^10^, prompting the need for image fusion techniques to combine the two datasets into one with superior image quality. An intuitive way to fuse the two datasets is to divide the two image stacks into halves and stitch the two sections that are closer to their respective illumination lenses. Hence, blur and ghost artifacts present in the halves far from the illumination lenses should be eliminated. While straightforward, a split along the midline may not align with local image quality, as it overlooks the complex geometrical shape and optical properties of the specimen, present even in simple, spherical objects.

Recent image fusion methods, therefore, follow a different strategy: they quantify the image quality—termed saliency level—of each pixel in the respective datasets and perform a weighted average of all the available inputs on an element-wise basis, such as two state-of-the-art methods, BigStitcher^18^ and MST-SR^23^. The fusion module in BigStitcher (“Fusion type” setting “Avg, Blending & Content Based”)^18^, which will be referred to as BigStitcher-Fuse in the following, measures the saliency level for each pixel in individual stacks based on local entropy and creates the fused image stack by giving greater importance to pixels that have a lower entropy (higher amount of information). MST-SR^23^, on the other hand, quantifies the pixel-level saliency level in a transformed contourlet domain to measure the amount of visual detail, e.g., edges and corners. However, both methods often include unwanted artifacts and blur into the fusion result, as they rely on merging all available inputs and thus fail to exclude low-quality regions and ghost artifacts.

Leonardo-Fuse employs a more advanced solution by integrating *a priori* knowledge of the illumination direction and the resulting image quality gradient. For each image pair illuminated from opposite sides, Leonardo-Fuse considers both local saliency level and the global impact of light propagation within the specimen to guide the fusion process (Fig. 4A-E, Extended Data Fig. 5A). Specifically, it first derives pixel-level saliency levels using non-subsampled contourlet transform (NSCT) and measures global light travel paths by identifying sample-related pixels through a simple foreground tissue segmentation. The obtained saliency levels are then re-weighted by prior knowledge of light travel paths, modeling the degree of image degradation due to light scattering as a linear function of the distance traveled by the light within the tissue (Extended Data Fig. 6). Using Bayes’ theorem and the expectation-maximization algorithm, Leonardo-Fuse estimates a fusion boundary that maps transitions from “good” to “bad” image quality (Fig. 4D). Additionally, spatial consistency across different depths is enforced. This fusion boundary is further refined using guided filtering to ensure smooth transitions near the fusion boundary, while maintaining distinct selections from each stack in areas farther away (Methods, Extended Data Fig. 5B, Extended Data Fig. 6). By excluding regions far from the illumination lens entirely, Leonardo-Fuse produces a final fused image free from ghost artifacts and blur.

**Fig. 4.**
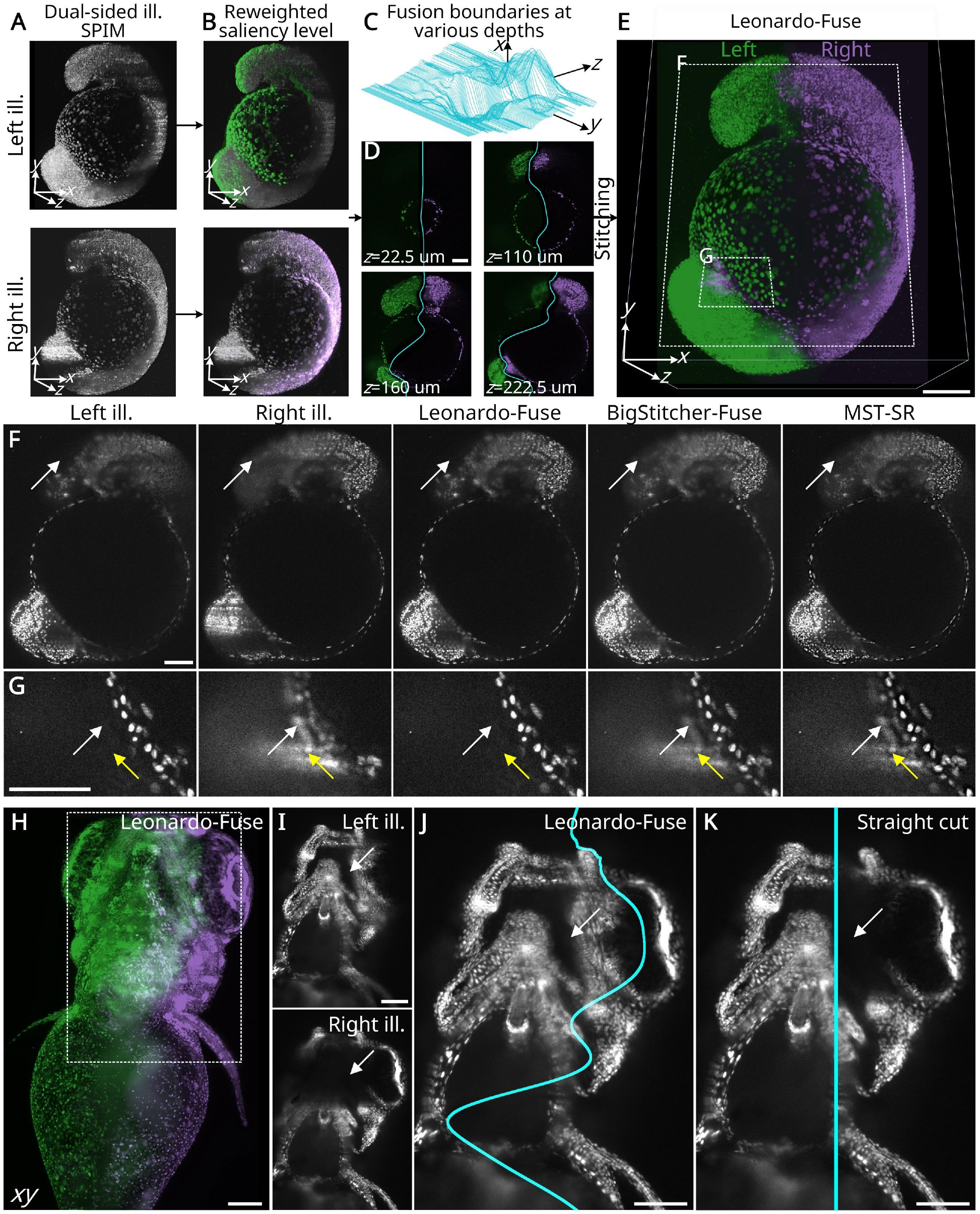
Leonardo-Fuse restores the most contrasted image for scattering tissues by fusing SPIM datasets with opposite illumination lenses. (A-B) In Leonardo-Fuse, local qualities of two SPIM datasets are quantified as saliency levels using non-subsampled contourlet transform. (C) A fusion boundary is then estimated by combining the saliency levels with the prior knowledge of the physical distance to the respective illumination lens. During optimization, the smoothness of the fusion boundary is considered both intra-slice along the y-axis and inter-slice along the z-axis. (D) Examples of fusion boundaries at different z-depths. (E) The fused image is obtained via stitching regions of better image qualities together from two stacks. Two patches with higher magnification of dotted rectangles are shown in (F) and (G). Leonardo-Fuse uniquely excludes both hazy blur (white arrows in (F)) and structured ghost artifacts (yellow arrows in (G)). By contrast, BigStitcher-Fuse and MST-SR introduce double signals (white arrows in (G)). (H-K) In a zebrafish embryo, a naive half-half fusion approach by straight cut neglects information in regions such as the shadowed area near the eye, whereas Leonardo-Fuse estimates a fusion boundary that well adapts to the complex geometry of the specimen (white arrows in (I), (J) and (K)). Scale bars: 100 μm.

We evaluated Leonardo-Fuse against state-of-the-art fusion methods on a zebrafish embryo labeled with H2B-GFP (Fig. 4E, Supplementary Table 2). While all methods succeeded in generating fused images of better overall quality than the original inputs, only Leonardo-Fuse effectively distinguished and excluded the fuzzy blur present on the distal side of the light sheet (white arrows in Fig. 4F). It also successfully eliminated structured ghost artifacts (yellow arrows in Fig. 4G) and provided a clear image of the sample (white arrows in Fig. 4G). This superior performance stems from its selective stitching approach, which uses estimated fusion boundaries (cyan curves in Fig. 4D) to exclude blurry regions on the distal side. The advantage of Leonardo-Fuse becomes particularly evident in samples with complex geometry. For example, in the instance of a zebrafish larvae, illumination from the right side is heavily absorbed by the eye, resulting in a large shadow with missing information (Fig. 4H, I, Supplementary Video 3). In this challenging scenario, Leonardo-Fuse accurately estimated a complex fusion boundary (cyan curve in Fig. 4J) that adapted to variations in local image quality, filling in the missing information with data from the opposite dataset on the left side of the fusion boundary (Fig. 4I). In contrast, a simple half-half cut would propagate missing information from the right dataset into the final fused image (Fig. 4K).

Overall, Leonardo-Fuse achieved a comprehensive, clear, and highly reliable reconstruction of sample information, confirmed by an intact intensity profile (Extended Data Fig. 6E-F) and cellular segmentation using Cellpose^22^ (Extended Data Fig. 7A-D). Notably, the unique ability of Leonardo-Fuse to eliminate ghost artifacts is further demonstrated when representing the outline of a 1-dpf (day post-fertilization) zebrafish labeled with Bodipy that results in heavy ghost artifacts. Due to ghost artifacts, neither Segment Anything (SAM)^24^ nor Segment Anything for Microscopy^25^ models accurately segment the zebrafish outline on BigStitcher or MST-SR fused images. By contrast, after eliminating ghosts, Leonardo-Fuse reconstruction enables an accurate outline segmentation (Extended Data Fig. 7E). Furthermore, we manually simulated light falloff in SPIM on a previously published PEGASOS-cleared mouse brain labeled with THY1-eGFP^26^ (Supplementary Note 5, Extended Data Fig. 8, Supplementary Fig. 10).

Quantitative evaluations using PSNR (44.9± 4.3) and SSIM (0.9847 ± 0.0015) on the simulated blur demonstrated Leonardo-Fuse’s effectiveness in handling SPIM with dual-sided illumination. Moreover, runtime benchmarking on the same dataset further highlights Leonardo-Fuse’s computational efficiency: it completed the fusion in just 18 seconds, whereas BigStitcher-Fuse required 1626 seconds, representing over a 90× speed-up.

### Leonardo-Fuse computationally extends the effective imaging depth along detection

Just as dual-sided illumination reduces the issue of light scattering and refraction during illumination, the emitted light is also scattered by tissue as it travels to the detection lenses. This leads to blur and image quality deterioration in regions farther from the detection lens. To expand the effective imaging depth along detection, i.e., *z*-depth that can be nicely imaged, dual-sided detection can be used, where the sample is rotated by 180°or two lenses are positioned on opposite sides of the sample to capture complementary datasets simultaneously (X-SPIM). In such a dual-sided illumination and dual-sided detection scenario, Leonardo-Fuse provides a solution to fuse all four datasets—each acquired using a unique combination of two illumination sources and two detection lenses—into one single high-quality stack. Specifically, Leonardo-Fuse detects the “good” to “bad” transitions along the detection axis in a manner similar to how the fusion boundary is estimated along the illumination axis. Therefore, given four complementary stacks and two fusion boundaries along the illumination axis, Leonardo-Fuse estimates two additional fusion boundaries along the detection axis (Extended Data Fig. 9A). To ensure no regions are overlooked or redundantly considered, Leonardo-Fuse (along detection) employs a decision-merging strategy (Extended Data Fig. 9B).

To evaluate its performance, we applied Leonardo-Fuse to a live 2-dpf transgenic zebrafish labeled with H2B-GFP imaged using an X-SPIM configuration (Fig. 5A, Supplementary Video 4). As expected, Leonardo-Fuse relied on patches captured by the front-side camera in the initial slices, gradually shifting to the back-side camera as it progressed to deeper layers of the specimen (Fig. 5C-E). A comparison with BigStitcher-Fuse and MST-SR showed that only Leonardo-Fuse restored the image without being affected by the ghost artifacts captured by the back camera (Fig. 5F). Additionally, we quantified image quality using an entropy-based information content metric, as previously described^27^ (Supplementary Note 7). The results revealed that the fused image stack generated by Leonardo-Fuse contained up to 1.5 times more information at any depth than the best individual stack from the four input datasets (Fig. 5B).

**Fig. 5.**
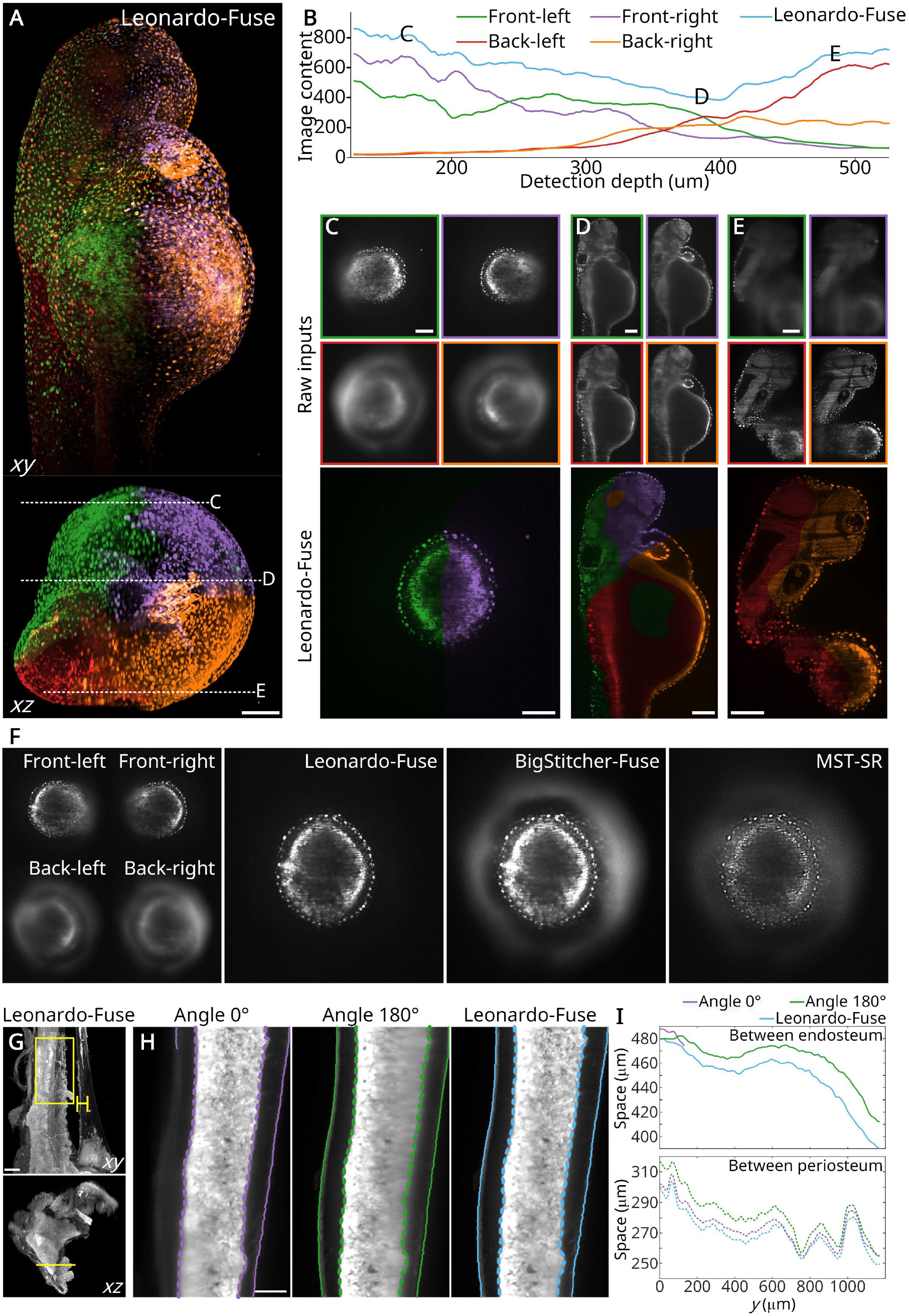
Leonardo-Fuse fuses information acquired via two opposite detection lenses into one. (A) Leonardo-Fuse efficiently combines four volumes, each illuminated from either left or right and detected from either front or back, into one fused volume. (B) The fused volume captures more information along all detection depths compared to the individual volumes. (C-E) Zoomed-in views of three regions show how Leonardo-Fuse seamlessly transfers the most trustful information from individual volumes into the fused one when imaging deeper into the specimen. Even in regions close to one detection lens (regions (C) and (E)), the fused volume still has better overall quality than the individual ones, demonstrating the algorithm’s effectiveness. (F) Compared to other algorithms like BigStitcher-Fuse and MST-SR, Leonardo-Fuse uniquely eliminates ghost artifacts in the final fused volume. (G) Leonardo-Fuse fused a murine tibia with endothelial staining (CD31) acquired using an Ultramicroscope Blaze. (H) Cropped regions from acquisitions at angle 0°, angle 180°, and the fusion result. Leonardo-Fuse successfully preserves the higher-quality information from each input (right-hand side from angle 0° and left-hand side from angle 180°), producing a seamless fused reconstruction. Solid and dotted lines mark periosteum and endosteum boundaries, respectively. (I) Thickness quantification along *y*, measured between endosteum (top) and periosteum (bottom). Accurate measurement is only possible from the fused result, as each individual acquisition alone lacks complete information. Scale bars: 100 μm (A-F, I, J), 500 μm (G), 300 μm (H).

### Leonardo-Fuse can be integrated with the registration workflow for efficient image fusion for other SPIM variants and very large specimens

With the four-lens X-SPIM systems providing well-aligned four volumes, Leonardo-Fuse is ready to reconstruct an optimal reconstruction of the specimen, minimizing the effects of light scattering and refraction. However, in many other light sheet-based systems (e.g. Blaze), dual-sided detection is achieved by rotating the specimen 180°around the *y*-axis (Fig 1A). While this rotation-based workflow is more economical, it requires the registration of the volumes imaged before and after rotation. To address this, we optimized an image-based registration workflow. The registration process begins with a 2D translational registration, followed by progressively more complex 3D registrations, from rigid to affine transformations, improving both speed and accuracy (Supplementary Fig. 11, Supplementary Notes 8-9, Supplementary Table 3). On a modestly sized zebrafish dataset (two volumes with 237 × 2048 × 2048 voxels, two volumes with 249× 2048 × 2048 voxels, Fig. 1F-H), our coarse-to-fine registration successfully aligns volumes collected before and after rotation, with the total registration time comparable to the fusion processing time (Supplementary Figs. 12-13).

With this registration process integrated, Leonardo-Fuse becomes more versatile and applicable to a wider range of SPIM variants (Supplementary Fig. 14, Supplementary Video 5). After demonstrating the effectiveness of Leonardo-Fuse on a range of medium-sized biological specimens, we turned our attention to centimeter-sized, chemically cleared tissues to assess whether Leonardo-Fuse can handle these large datasets efficiently without overwhelming computational resources. We studied murine tibia with endothelial staining (CD31) using the UltraMicroscope Blaze system equipped with a rotatable sample holder (Methods, Fig 5G). As the UltraMicroscope Blaze system uses simultaneous opposing illuminations, only two datasets (captured before and after rotation, i.e., angle 0° and angle 180°) needed to be fused. Yet, the large data size posed a significant computational challenge. Each dataset was typically more than 1000×2048×2048 voxels (approximately 5.71×5.55×5.55 mm^3^). We optimized Leonardo-Fuse for efficient performance on a moderate workstation (Extended Data Fig. 10A, Supplementary Note 10). Specifically, we differentiate between operators that can be performed in a downsampled space without losing accuracy, such as estimating the fusion boundary, or those that require the full resolution, as in the case of registration. Processing both registration and fusion took approximately 30 minutes on a workstation equipped with 51 GB of memory, an Intel(R) Xeon(R) CPU @ 2.20GHz, and a single NVIDIA T4 Tensor Core GPU with 15 GB of memory—a manageable computational requirement. Importantly, performance is not compromised, as the system selectively integrates sample information along the depth while effectively rejecting blur (Fig. 5G-I). Accurate quantitative measurement of structures is important for studying the closed circulatory system in long bones^28^. For example, one can measure the distance between the periosteum (dashed color lines in Fig. 5H) and between the endosteum (solid color lines in Fig. 5H) to quantify potential inflammation. However, due to image blur, segmentation of periosteum and endosteum on the original single-view images is either impossible (e.g. partially vanishing purple solid line in angle 0° in Fig. 5H) or less precise (dashed green line in angle 180° in Fig. 5H). Only Leonardo-Fuse’s reconstruction allows precise segmentation of both periosteum and endosteum and hence accurate quantification of their distances (Fig. 5I, Supplementary Note 11). Moreover, we also segmented Trans-cortical vessels (TCVs) (Extended Data Fig. 10C-E) using a vesselness filter^29^. As shown in the example images and the associated Intersection over Union (IoU) values, Leonardo-Fuse produces segmentation results nearly identical to those obtained from the higher-quality acquisition (angle 0° in Extended Data Fig. 10D). Moreover, when both acquisitions are partially degraded, Leonardo-Fuse effectively merges complementary information to generate the most faithful segmentation relative to the manually traced ground truth (IoU = 0.9 vs. 0.69 or 0.23, Extended Data Fig. 10E).

### Optimal integration of Leonardo-DeStripe and -Fuse modules for large samples

Our original workflow (denoted as Leonardo-(DeStripe→Fuse), Fig. 1F-H) recommended applying Leonardo-DeStripe before Leonardo-Fuse, which works well for most samples. However, DeStripe can struggle with deep shadows caused by severe light blockage, where structures are barely visible (white arrows, Supplementary Fig. 15). These regions are better corrected by the Fuse module, which uses data from the opposite illumination direction.

To address this, we recommend an optimized workflow, Leonardo-(Fuse→DeStripe), which first applies image fusion along the illumination axis, followed by stripe removal, and then fuses along the detection axis (Supplementary Figs. 15–17, Supplementary Notes 12–13). This fusion-first strategy contrasts with the Leonardo-(DeStripe→Fuse) workflow, where stripe removal is applied separately to each orientation. On a 1-dpf zebrafish labeled with Bodipy, Leonardo-(Fuse→DeStripe) improved image quality by reducing blurriness, ghost artifacts, and large shadows while effectively eliminating residual thin stripes (Fig. 6A). Importantly, fine sample structures were preserved, as confirmed by membrane segmentation using Cellpose (Fig. 6B), underscoring the biological fidelity of the reconstruction. Beyond image quality, this workflow also improved computational efficiency by requiring only a single stripe removal operation, reducing processing time from 160 minutes to 102 minutes on a single NVIDIA T4 GPU (Fig. 6C). Together, these results demonstrate that Leonardo-(Fuse→DeStripe) not only synergistically integrates the -Fuse and -DeStripe modules but also achieves superior balance between accuracy and scalability for large-scale biological imaging.

**Fig. 6.**
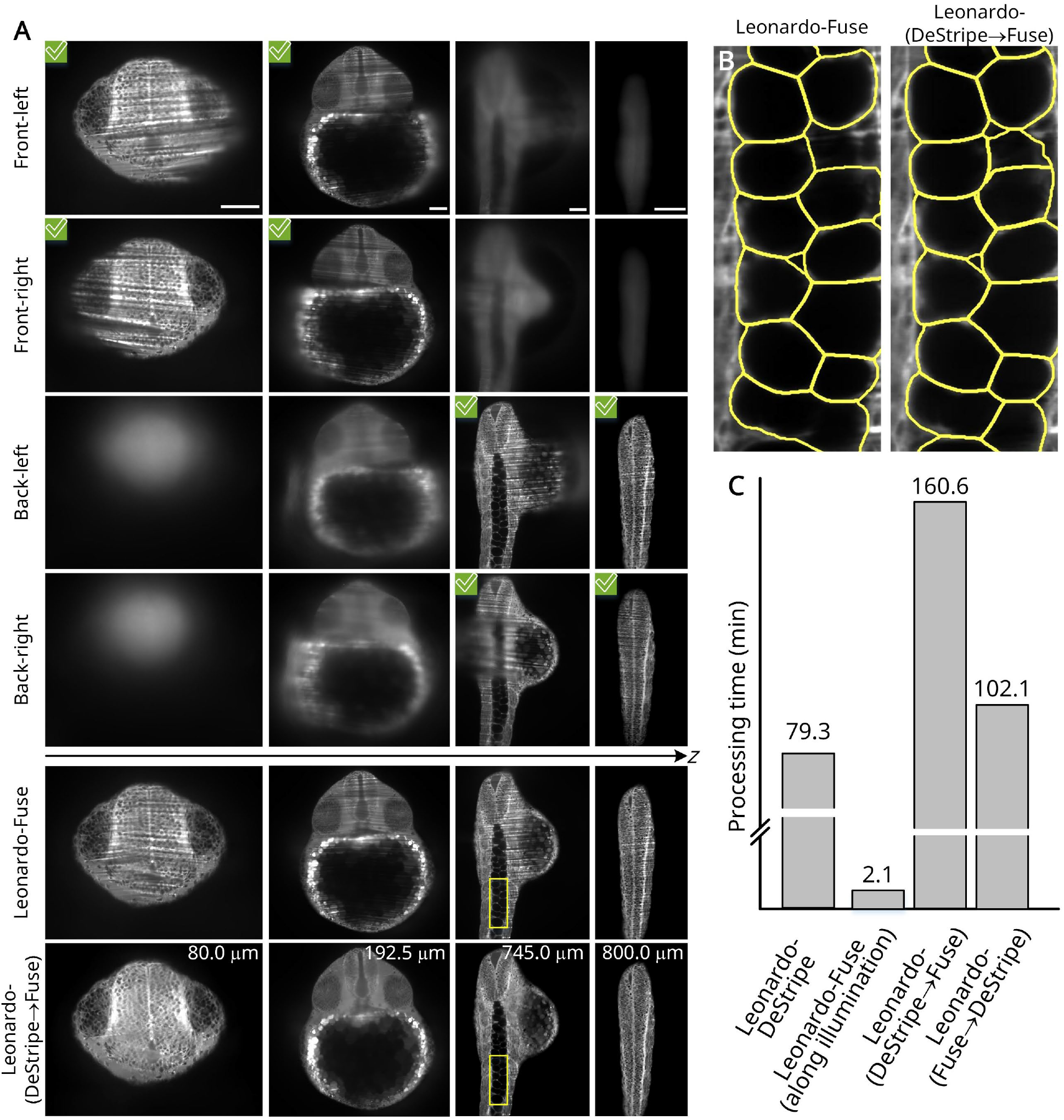
Leonardo-(Fuse→DeStripe), a synergistic integration of Leonardo-DeStripe and -Fuse. modules. (A) On a 1 dpf zebrafish labeled with Bodipy, Leonardo-(Fuse→DeStripe) firstly applies the - Fuse module to remove blurriness, ghost artifacts, and large shadows, followed by the -DeStripe-Fuse module to remove the remaining thin stripes. Inputs marked with green ticks in the top-left corner are with better image quality and selected by Leonardo. (B) Comparison between Leonardo-Fuse and Leonardo-(Fuse→DeStripe) in a zoomed-in region, showing that sample structures remain intact after stripe removal, as validated by membrane segmentation using Cellpose<u>^22^</u>. (C) Compared to a Leonardo-(DeStripe**→**Fuse) workflow which applies Leonardo-Fuse after Leonardo-DeStripe, Leonardo-(Fuse→DeStripe) saves processing time by a huge margin. Scale bars: 100 μm.

## Discussion

We present Leonardo, a pioneering set of AI-based image processing tools, to reduce sample-induced artifacts in SPIM datasets. Our toolbox comprises two modules: Leonardo-DeStripe and Leonardo-Fuse. Unlike traditional methods, Leonardo-DeStripe’s use of a GNN-parameterized 2D band-pass filter is adaptable to the specific characteristics of each sample, ensuring more effective and consistent stripe removal across diverse imaging conditions, particularly in complex multi-directional SPIM setups. This adaptive approach not only addresses a longstanding challenge in SPIM but also opens new insights into structured noise removal.

Leonardo-Fuse introduces a globally optimized fusion strategy that integrates datasets from dual-sided illumination and/or dual-sided detection, surpassing traditional methods that often struggle with ghost artifacts and blur. Further equipped with an image-based registration workflow, Leonardo-Fuse is adaptable to a wide range of SPIM-based systems and compatible with rotatable sample holders. Optimized for efficiency, Leonardo-Fuse requires only modest computational resources, facilitating its adoption by users who seek to achieve the most contrasted projections of extremely large, complex tissues with minimal human intervention. Notably, Leonardo-Fuse also delivers substantial speed advantages for image fusion, reducing the processing time of fusing a typical zebrafish embryo with dual-sided illumination from several hours with the existing tool to under ten minutes.

Leonardo was designed with a moderate parameter size, making it both efficient to train and execute. Unlike large, computationally intensive networks, Leonardo’s design allows it to be trained effectively in a self-supervised manner using only the current volumes being processed. This design choice ensures that Leonardo can run on standard workstations without requiring high-end computational resources, thereby improving accessibility for a wider range of users. This approach contrasts with prevailing trends in computer vision, particularly in generative foundation models that rely on large datasets and significant computational power for training. The difference likely stems from the distinct scopes of biological image analysis and general computer vision tasks. In general computer vision, models must perform well across a wide variety of test images, necessitating large and diverse training datasets. In contrast, Leonardo focuses on specific types of biological images, allowing it to deliver robust and reliable results tailored to the unique challenges of SPIM while maintaining computational efficiency.

While Leonardo is designed as a standalone toolbox, it can complement and enhance other image processing toolboxes, particularly in the context of the prevalent generative foundation models. These models, which are publicly available for image restoration, can be integrated as post-processing steps following Leonardo to push the limits even further (Supplementary Fig. 18)^30^. One aspect not directly addressed by Leonardo in SPIM is multi-view reconstruction^15,19^ (fusion or deconvolution), which is particularly useful for overcoming the inferior axial resolution of SPIM by leveraging better lateral resolution from other views (Supplementary Fig. 19). We found that multi-view reconstruction has been extensively explored and is well-supported in several image processing toolboxes, such as BigStitcher and Huygens. Although these tools perform reasonably well, preprocessing input volumes with Leonardo—removing stripes, blur, and ghost artifacts—can significantly enhance the quality of multi-view reconstruction. By providing clearer, artifact-free inputs, Leonardo enables these existing tools to deliver even better results.

Although initial results with Leonardo are promising, we envision several extensions to our work. First, for Leonardo-DeStripe, it would be valuable to consider inter-slice information when removing stripe defects. In practice, we found that biological specimens typically exhibit smooth continuity across adjacent slices, in comparison to stripes that abruptly change between slices. Therefore, the inherent structural consistency of the reconstructed specimen in 3D can be useful when regularizing Leonardo-DeStripe. In addition, since the current GNN is retrained for each individual slice, one important future direction is to explore reusing the model trained on one slice to initialize or guide the optimization on adjacent slices. This may help reduce the retraining overhead without sacrificing per-slice adaptiveness. Notably, since each slice is processed independently, the retraining procedure is in parallel and can be efficiently distributed across multiple GPUs or compute nodes. We plan to make such distributed inference pipelines available in future software releases to improve scalability on large datasets. Finally, while the dedicated post-processing module significantly improves Leonardo-DeStripe’s ability to preserve fine sample structures, further refinements are still warranted and represent a valuable avenue for future research.

Meanwhile, we anticipate potential improvement regarding the processing speed of Leonardo-Fuse. For example, the feature extraction step within the Fuse module can be further accelerated by utilizing single-precision floating-point numbers. Ideally, it would be particularly advantageous if Leonardo-Fuse could be deployed on-the-fly, such that the terabyte-scale datasets, after fusion, can be condensed and saved with fewer storage requirements. If Leonardo-Fuse achieves real-time operation, it could be potentially integrated with image acquisition systems to enable smart microscopy^31^. In addition, although the improved coarse-to-fine registration workflow achieved satisfying registration automatically, its performance highly depends on the amount of overlapped information in the specimen before and after rotation. When only a limited amount of overlapping information exists between datasets, the performance of the registration will inevitably be compromised. Therefore, we recommend that users of Leonardo-Fuse manually register the input stacks beforehand or consider using bead-based registration plugins^18^, which require embedding fluorescent beads in the mounting medium around the specimen.

The Leonardo toolset is not limited to SPIM-based large and/or living images illustrated here. In particular, stripe artifacts can plague micrographs across any length scale, e.g., from Focused Ion Beam Scanning Electron Microscopy (FIB-SEM)^32^ to atomic force microscopes (AFM)^33^. Therefore, Leonardo-DeStripe holds significant potential for broad application across various microscopy techniques. Meanwhile, Leonardo-Fuse has the potential to be incorporated with lifelong learning, enabling it to adapt quickly and promptly in time-lapse experiments. In conclusion, with its versatility, extensibility, and user-friendly GUI design, we anticipate that Leonardo will become a powerful tool in the biologist’s arsenal for biological imaging, particularly in SPIM applications.

## Acknowledgments

Y.L. is supported by the China Scholarship Council (No. 202106020050). G.F.M. is supported by the MWK (Niedersächsisches Ministerium für Wissenschaft und Kultur, 6707040) and is member of the Hertha Sponer College of the Cluster of Excellence Multiscale Bioimaging (MBExC; EXC 2067/1-390729940), University of Göttingen, Germany. T.C. is supported by the Helmholtz Association under the joint research school “Munich School for Data Science - MUDS”. J.H. is supported by the Alexander von Humboldt Foundation (Alexander von Humboldt Professorship; J.H.) and the German Research Foundation (Germany’s Excellence Strategy EXC 2067/1-390729940; J.H.).

## Author contributions

Conceptualization: J.H., T.P.; Machine learning methodology: Y.L.; Software: L.K., J.C., Y.L., G.F.M.; Microscope design: G.F.M., J.H.; Validation: Y.L., G.F.M., L.K., T.C.; Formal analysis: Y.L., G.F.M.; Investigation: Y.L., G.F.M.; Resources: G.F.M., K.W., P.M., M.R., M.S., A.G., J.L., J.P., A.E., J.H.; Data Curation: G.F.M., Y.L., L.K., J.C.; Writing - Original Draft: Y.L., G.F.M.; Writing - Review & Editing: Y.L., G.F.M., T.C., K.W., J.L., N.N., C.M., J.C., J.H., T.P.; Visualization: Y.L., G.F.M.; Supervision: J.H., T.P.

## Data availability

A sample transgenic zebrafish dataset required using X-SPIM (shown in Figs. 5) is available at https://zenodo.org/records/14215090. The remaining samples are either publicly available at the link provided or can be requested from the original investigators:

- wild-type mouse body (https://discotechnologies.org/wildDISCO/atlas/index.php)
- PEGASOS-cleared mouse brain (https://zenodo.org/records/5639726)

## Code availability

Leonardo is a pip installable Python package and is available at the following GitHub repository: https://github.com/peng-lab/leonardo_toolset, with documentation at https://leonardo-toolset.readthedocs.io/en/latest/. Tutorials regarding the use of Leonardo as Notebooks and Napari plugins can be found https://leonardo-toolset.readthedocs.io/en/latest/tutorials.html.

## Methods

### Leonardo-DeStripe

#### Graph neural network regularized with anisotropic total variation unit

In SPIM, stripe removal is to restore the underlying stripe-free volume *X* from its stripe-corrupted observation *Y*:

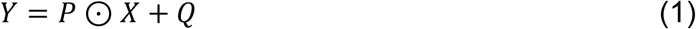

where 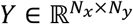 is the SPIM input at an arbitrary *z*-plane, with a size of *N*_*x*_ × *N*_*y*_, ⊙ is element-wise product, 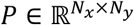 and 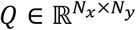 are the multiplicative distortions caused by stripes and additive noise, respectively.

Additionally, Leonardo-DeStripe notes that adjacent entities in *P*, within a stretched window along the stripe orientation, tend to be similar, as the attenuating obstacles that block the illumination light and cause the stripes are sparse. Thus, Eq. (1) can be rewritten as:

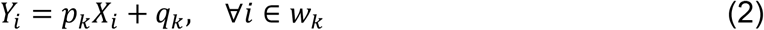

where *w*_*k*_ is a local window centered at pixel *k, p*_*k*_ and *q*_*k*_ are shared within *w*_*k*_. Moreover, considering the oblong morphology of the stripes, Eq. (2) holds only when *w*_*k*_ of size 1 × *w*, assuming a horizontal illumination direction.

In general, completely removing stripes while preserving all sample information in SPIM is challenging, particularly when stripe artifacts and sample signals are entangled. However, it is much more feasible to first obtain a suboptimal estimation 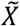 (with stripes removed completely but sample details compromised) of *X*, and later refine it to achieve a closer approximation of the true stripe-free *X*. This refinement can be achieved using guided filtering (GF)^34^, a recent edge-preserving filtering technique:

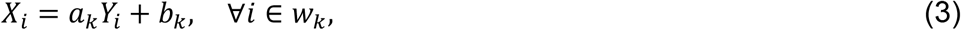

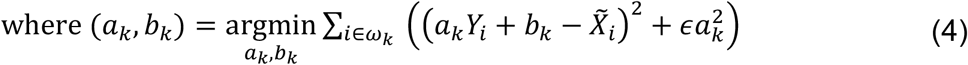

where Eq. (3) is the inverse of Eq. (2) with *a*_*k*_ = 1/*p*_*k*_ and *b*_*k*_ = −*q*_*k*_/*p*_*k*_, and ε is a penalization parameter preventing *a*_*k*_ from being too large. Suboptimal 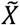 is hence refined in two aspects: first, by ensuring better data integrity through the local linear transformation in Eq. (3), which transfers visual details from *Y* to *X*; and second, by maintaining a stripe-free property, as the learned *X* is visually similar to the stripe-free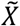, ensured by the mean squared error (MSE) in Eq. (4). To transfer as many textures from *Y* to *X* as possible, the hyperparameter ε is set to be small (1*e*^−6^), and the row-wise window size *w* is kept large and conditioned on the size of the specimen (Extended Data Fig. 2A).

Next, Leonardo-DeStripe aims to estimate 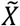, which can be efficiently realized in a down-sampled space, inspired by the local smoothness of attenuation matrix *P*. Specifically, Leonardo-DeStripe translates the estimation of 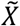 into logarithmic space, where stripe degradation becomes additive, rather than a multiplicative one, with respect to the underlying 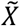. This transformation simplifies the problem and allows for the optimization of the following image restoration framework using the split Bregman algorithm^21^:

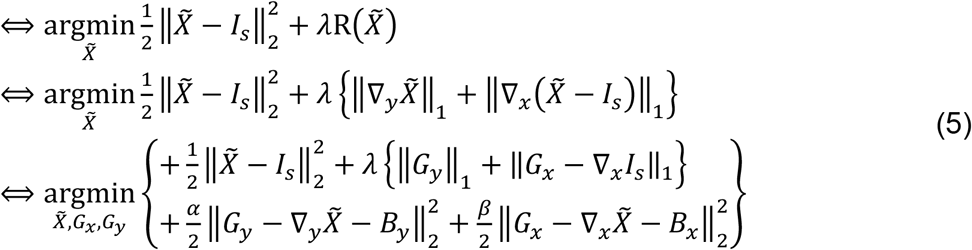

where *I*_*s*_ = logDS(*Y*) is the stripe-corrupted *Y* after downsample DS(·) and logarithm log(·), ‖·‖_2_ is the *l*_2_-norm to encourage similarity between the prediction 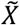 and *I*_*s*_, R(·) is the regularizer that enforces desirable properties onto 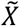, and *λ* is the trade-off parameter. For the stripe removal, it is prevalent to employ anisotropic total variation (TV) for the regularizer, as stripes exhibit strong directional characteristics, causing significant gradient variations across the stripes while gradients along the stripes remain relatively unaffected. Thus, for the horizontal stripes, 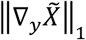 and 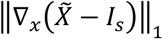 are to suppress the gradient across the stripes and to preserve gradients along the stripes, respectively, where ‖·‖_1_ is *l*_1_-norm, ∇_*y*_ and ∇_*x*_ denote the vertical and horizontal derivative operators, respectively. Furthermore, the intractable optimization of 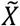 can be simplified by decoupling it into three alternating and simpler minimizations over 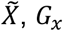, *G*_*x*_ and *G*_*y*_, by introducing auxiliary variables 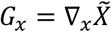 and 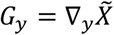, along with the Bregman variables *B*_*x*_ and *B*_*y*_:

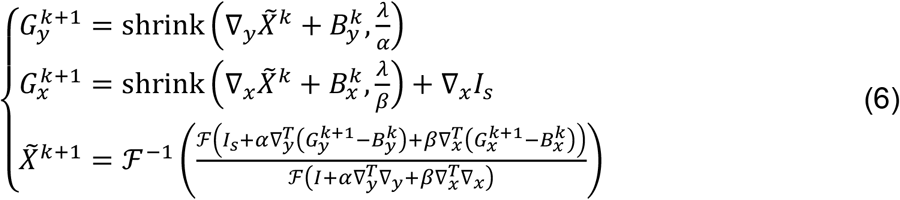

where subscript *k* denotes the *k*-th iteration, 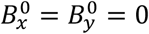, shrink(*r, ξ*) = *r*/|*r*| max (*r* − *ξ*, 0) is a shrinkage operator, *I* is an identity matrix, ℱ(·) and ℱ^−1^(·) are the Fourier transform and its inverse, respectively (Supplementary Note 2.1). Here, Leonardo-DeStripe identifies two significant limitations in such an anisotropic TV-based stripe removal framework. First, the term 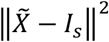 in Eq. (5), which was originally intended to ensure data fidelity under the assumption of Gaussian noise, is inappropriate for stripe noise, as the zero-mean and pixel independence of the Gaussian distribution do not hold for structured noise. The spatially correlated nature of stripes, moreover, renders the shrinkage operator ineffective, as the global threshold *ξ* cannot be adjusted at the pixel level (Supplementary Note 2.2). Thus, Leonardo-DeStripe enhances the conventional stripe remover described earlier as (Extended Data Fig. 2B-C):

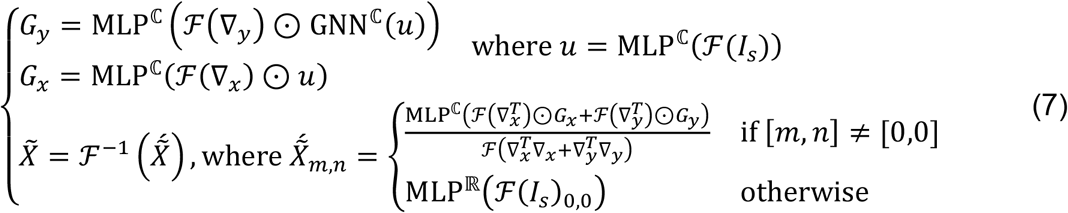

where each row corresponds to the respective row in Eq. (6), with subscripts omitted, as Leonardo-DeStripe operates as a one-iteration split Bregman solution enhanced by deep learning, GNN(·) and MLP(·) are deep learning components proposed in Leonardo-DeStripe. Note that the element-wise multiplication of ℱ(∇_x_) in Fourier, i.e., ℱ(∇_x_) *⊙*, is equivalent to the convolution operation of ∇_*x*_ in the pixel space (the same applies to ∇_*y*_, 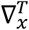 and 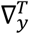).

Specifically, Leonardo-DeStripe first optimizes *G*_*y*_, to suppress the gradient in the direction perpendicular to the stripes, using the proposed GNN-based 2D band-pass Fourier filter (Fig. 2A). Inspired by the directionality of the stripes, a band-pass filter in the Fourier domain that suppresses specific frequencies affected by the stripes should be effective for stripe removal. Typically, the stripe-related Fourier coefficients are concentrated within a narrow wedge-shaped region perpendicular to the stripes. However, the angular coverage of the wedge region, ideally large enough to remove stripes completely, is limited to avoid removing sample information within the filter. Hence, Leonardo-DeStripe constructs a significantly larger wedge mask with an angular coverage of ±29°(predefined but user-adjustable, Supplementary Fig. 20). Within this mask, Fourier coefficients are replaced by a combination of their uncorrupted neighbors—those outside the wedge mask—on a polar coordinate system. This approach is inspired by the ideally homogeneous Fourier projection of the specimen without stripes (Supplementary Note 3, Supplementary Table 4)^35^. Specifically, the Fourier projection ℱ(∇_*y*_) *⊙ u* can be reformulated as a graph 𝒢 = {𝒱, *H, A*}, where 𝒱 is the vertex set of |𝒱| = *N*_*x*_ × *N*_*y*_ nodes, *H* ∈ ℂ^|𝒱|×*d*^ is the Fourier coefficients associated with each node (*d* is the dimension of node attributes), *A* ∈ ℝ^|𝒱|×|𝒱|^ is the adjacency matrix with (p, *q*)-th entity denoting the connection from node *q* to *p*. Additionally, we impose sparsity on 𝒢, by randomly selecting a total of *N* neighbors based on polar coordinates for each node falling within the wedge mask, while reserving only self-loop for nodes outside the mask:

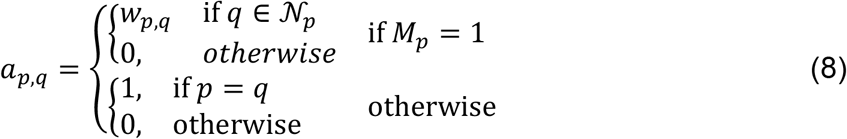

where *M* is the wedge-shaped mask with 1 and 0 denoting within and outside the mask, respectively, *w*_*p*,q_ ∈ [0,1] is the “strongness” that node *q* is influencing *p*, and is set as learnable weights in GNN, 𝒩_p_ denotes the neighboring set of the node *p*. As a result, GNN selectively propagates messages across 𝒢:

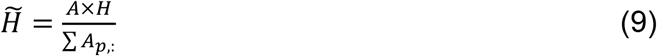

where stripe-affected nodes 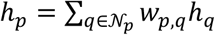 are replaced as a weighted and normalized combination of their neighbors (Extended Data Fig. 2D).

Next, Leonardo-DeStripe models *G*_*x*_ using a multilayer perceptron (MLP). Given that *G*_*x*_ inherently acts as a fidelity term to minimize the difference between the desired and striped images in their first-order derivatives, solving *G*_*x*_ is equivalent to optimizing a *G* with structures similar to *u* along the *x*-axis and reduced noise. In this regard, an MLP operating in the Fourier space is particularly effective:

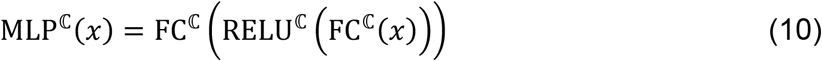

where FC^ℂ^(x) = *xW*^*T*^ + *b* is a fully connected (FC) layer in the complex domain where 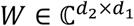and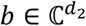, whereas RELU^ℂ^(x + *yi*) = RELU(x) + RELU(y)*i* is a rectified linear unit (ReLU) activation function in the complex domain, with *i* being the imaginary unit. Here, FC^ℂ^(·), which has a 1 × 1 receptive field in the Fourier domain but a global perspective of the entire image in the pixel space, is efficient for denoising^36^. It is also sufficient for ensuring data fidelity, as it can easily degrade into an identity mapping when 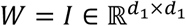 and *b* = 0, where *I* is an identity matrix in the real domain. To further preserve fidelity, the same MLP architecture is incorporated into the process of solving *G*_*y*_ right after the GNN module.

Finally, given *G*_*x*_ and *G*_*y*_, Leonardo-DeStripe loosely follows the conventional split Bregman to generate 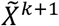, which jointly considers the similarity of the desired 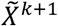 with 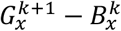 along the *x*-axis, 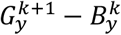 along the *y*-axis, and *I* in the Fourier space. The major difference is that Leonardo-DeStripe, as a single-pass estimation of the stripe-free solution, does not consider the contaminated striping *I*_*s*_ when constructing 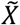. Consequently, the zero frequency in Fourier, representing the direct current (DC) component of 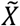, needs to be estimated separately, as the two derivative operators ∇_*x*_ and ∇_*y*_ are zero-mean. Thus, we formulate a separate DC branch to infer 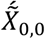, where 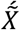 is the Fourier projection of 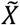, using an MLP in the real domain, since 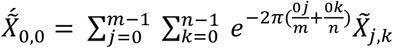 should always be real. The DC component of stripe-corrupted *I*_*s*_ is fed as input to the MLP. Additionally, the process of composing 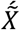 is parameterized using a complex MLP, where Bregman variables 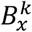 and 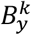are initialized as zero and thus can be omitted in Leonardo-DeStripe.

Additionally, inspired by the nature of discrete Fourier transform—the Fourier projection of a real-valued signal, e.g., our SPIM image, exhibits conjugate symmetry— Leonardo-DeStripe only learns the positive frequencies in the last dimension of 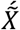. The full 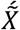 can be obtained by concatenating the conjugate-flipped positive frequencies, the DC component, and the learned positive frequencies in the last dimension of 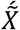accordingly.

During the training of Leonardo-DeStripe, the loss function is given by:

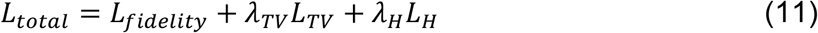

where *L*_*fidelity*_ is the fidelity term, *L*_*TV*_ and *λ*_*H*_ are the TV-based and Hessian-based regularization terms, respectively, and *λ*_*TV*_ and *λ*_*H*_ are the corresponding trade-off parameters. Specifically, given that MSE has already been proven to be an inappropriate criterion for measuring similarity in stripe removal tasks, a GF-based similarity criterion is proposed:

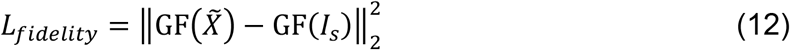

where stripes in 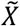 and *I*_*ss*_ are averaged out/blurred separately using GF with large penalization *ε* and column-wise *w*_*k*_ (Supplementary Note 2.3).

Next, anisotropic TV is reused in the loss term:

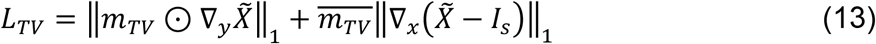

where 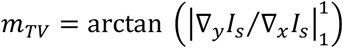 is a spatially adaptive weighting map designed to emphasize smoothing across stripes in regions where the underlying structure exhibits strong *y*-axis features. arctan(·) is the element-wise arc tangent. 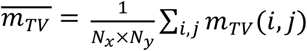 is the spatial average of the mask.

In addition to the TV-based regularizer, which penalizes first-order gradients to suppress stripe-like noise, Leonardo-DeStripe also incorporates a second-order regularization term based on the Hessian. While anisotropic total variation encourages piecewise constancy and is effective at removing stripes, it may inadvertently suppress fine structural details. The Hessian-based regularizer, on the other hand, enforces continuity of second-order derivatives, which better preserves smoothly varying biological structures such as curved membranes and neurites (Supplementary Fig. 21):

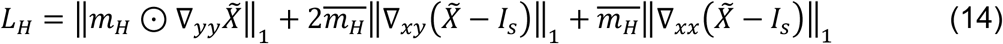

where ∇_*xx*_, ∇_*xy*_ and ∇_*yy*_ are the second-order derivative operators, 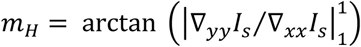 is a spatially adaptive weighting map similar to *m*_*TV*_ in the TV-based term.

### Post-processing module for sample structure preservation

#### Guided upsampling

By using the trained graph neural network regularized by anisotropic total variation unit, Leonardo-DeStripe can effectively remove the stripe artifacts. However, two limitations remain. The first is that the network, which operates in a downsampled space, requires an upsampling strategy to reconstruct the full-resolution output without losing fine sample details. To address this, we introduce a guided upsampling operator, which first upsamples the graph neural network’s output 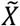:

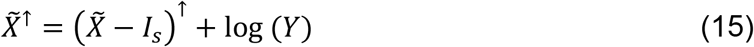

where ↑ is the upsampling operation, for which we use bilinear interpolation. Here, rather than directly upsampling the predicted 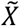, we upsample the residual component 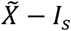. This is motivated by the observation that the stripe artifacts tend to be smooth and low frequency—properties that make the residual particularly well-suited for interpolation. In contrast, directly interpolating 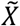 may blur fine sample features.

Next, as discussed earlier, 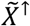 is only a suboptimal estimation, as the GNN is primarily trained to remove stripes and may sacrifice subtle sample details in the process. This is particularly evident when sample structures align with the stripe direction. This limitation arises from the unidirectional total variation regularization term in the loss function, whose receptive field is so local that it fails to distinguish stripe shadows and similarly oriented sample structures. Thereby, the guided upsampling is to recover subtle structural variations by refining 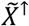 using the GF defined in Eqs. (3) and (4). Since we are still operating in the logarithmic domain, the refinement takes the form:

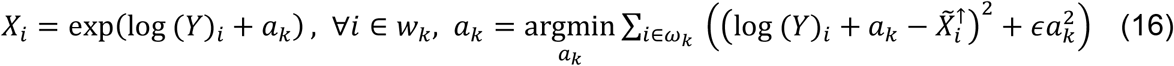

where *b*_*k*_ in Eqs. (3) and (4) is omitted, as the additive noise in SPIM is typically negligible. There is a closed-form solution 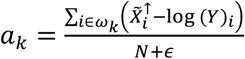 where *N* = |*ω*| denotes the number of pixels in the local window *ω*_k_. However, in practice, we found that replacing the mean-based solution with a median filter yields better results, as it more effectively preserves edges and suppresses outliers in the residual. As a result, the final refinement becomes:

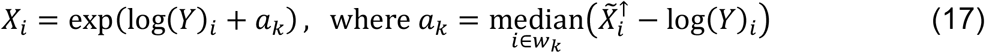

where median(·) is the median filter. Conceptually, a larger |*ω*_k_| helps better preserve sample structural continuity. However, it may also increase the risk of retaining residual stripe artifacts, as the filter becomes less sensitive to small intensity variations. In Leonardo-DeStripe, |*ω*_k_| is set to be 49 by default (Supplementary Fig. 22). By using the guided upsampling operator, Leonardo-DeStripe can restore small sample structures that are erroneously removed previously by the GNN (Extended Data Fig. 1).

#### Global post-processing using illumination prior knowledge (ill. prior)

As discussed earlier, to prevent residual stripes, the filtering window used by guided upsampling is limited in terms of size. Hence, relatively large directional sample structures that align with the stripe orientation cannot be restored after guided upsampling (Extended Data Fig. 1B-C). As a result, to further suppress stripe artifacts while preserving underlying sample structures, we introduce a dedicated post-processing module by modeling the directional and global propagation of the stripes along illumination (ill. prior). Specifically, this module is motivated by a physical observation in the SPIM system: true stripe artifacts exhibit consistent directional propagation through the sample volume:

- Dark stripes correspond to consistent absorption or occlusion, and thus become progressively darker.
- Bright stripes result from the accumulation of light along the illumination direction and become progressively brighter.
- In contrast, biological structures typically exhibit non-monotonic, localized intensity patterns that do not persist uniformly along any fixed direction.

Therefore, when post-processing the residual prediction 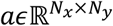 (which represents stripes and erroneously removed sample structures, in logarithmic space) with each entity estimated based on Eq. (17) via guided upsampling, ill. Prior first extracts its horizontal directional gradient (to process horizontal stripes) and splits it into two channels: the positive part *a*^+^ corresponding to bright stripes and the negative part *a*^−^ corresponding to darkening (attenuation):

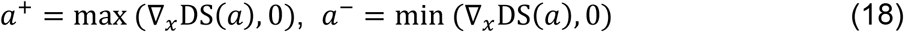

where DS(·) is the downsampling operator along the *x*-axis. Next, *a*^+^ and *a*^−^ are scaled by two learnable parameters, *w*^+^ and *w*^−^, to adjust intensity:

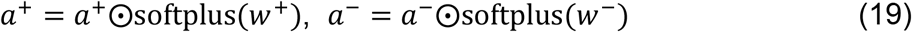

where softplus(·) ensures positivity. A cumulative summation can then be applied along the stripe direction, i.e., the *x*-axis, to simulate directional propagation:

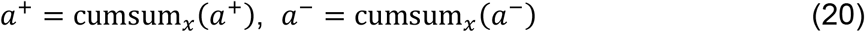

*a*^+^ and *a*^−^ are then blended to mimic the composition of bright and dark stripes:

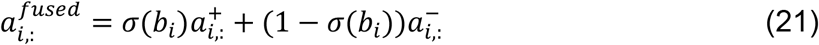

where the *i*-th row is composed via learnable parameter *b*_*i*_, *σ*(·) is the sigmoid function. Additionally, directional propagation through cumulative summation may introduce global intensity drift, especially when residuals consistently propagate in one direction. To mitigate this:

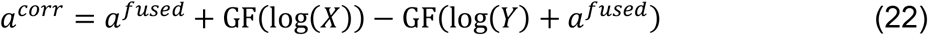

where GF(·) uses large *ε* = 10 and large window of 49× 49to smooth out details. The final stripe prediction *a*^*final*^ can be obtained via bilinear upsampling.

To train ill. prior, unidirectional TV-based loss is reused again:

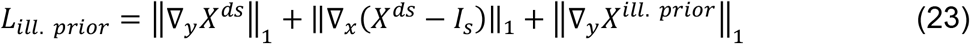

where *X*^*ds*^ = *a*^*corr*^ + DS(log (Y)) and DS(·) is the same downsampling in Eq. (18), *X*^*ill*.*prior*^ = *a*^*final*^ + log (Y) is the stripe-free reconstruction in full-resolution, *I*_*s*_ = DS(log (Y)). As a result, only monotonic stripes are retained by ill. prior, while true sample structures are preserved.

### Leonardo-DeStripe for advancing cases

Beyond removing horizontal stripes, Leonardo-DeStripe is extensible to more challenging applications, such as stripes with arbitrary angular orientation and even stripes along multiple directions. Specifically, when removing stripes along, neither *x*-nor *y*-axes, but arbitrary degree *θ*, the only adjustments made to the network of Leonardo-DeStripe are to rotate the angular orientation of the wedge-shaped mask and the anisotropic TV unit (Supplementary Note 2.4). Additionally, for multi-directional SPIM, the anisotropic TV unit in Leonardo-DeStripe is replicated multiple times according to the total number of stripe directions Θ, each of which is to resolve stripes along a specific orientation only. Outputs from these units are then concatenated channel-wise and merged using a fully connected layer with weight *W* ∈ ℂ^(Θ×*d*)×*d*^ and bias *b* ∈ ℂ^*d*^, and finally fed into the subsequent operations (Fig. 3A).

In parallel to the network design, the loss function is also extended in a direction-aware manner. Specifically, the TV-based and Hessian-based regularization terms are computed independently along each candidate stripe orientation using rotated derivative kernels (Supplementary Note 2.4). In the multi-directional SPIM case, where *θ* ∈ {*θ*_1_, *θ*_2_, ⋯, *θ*_*K*_} = Θ, for each pixel, the direction with the strongest directional response— quantified via *m*_*TV*_ in Eq. (13) and *m*_*H*_ in Eq. (14)—is selected as the dominant stripe orientation. As a result, let 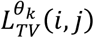 and 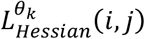 denote the TV-based and Hessian-based loss at each pixel (*i, j*) along direction *θ*_*k*_ ∈ Θ, the final regularization losses are aggregated as:

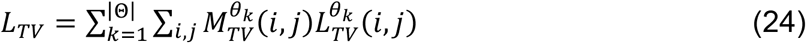

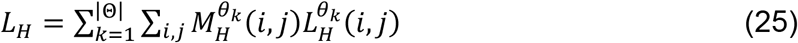

where *M*_*TV*_(*i, j*) ∈ {0,1}^|Θ|^ and *M*_*H*_ (*i, j*) ∈ {0,1}^|Θ|^ are one-hot selection masks indicating the dominant stripe direction at each pixel based on the weighting maps *m*_*TV*_ and *m*_*H*_, respectively. This strategy ensures that each regularization term is enforced only along the most relevant direction at each location.

### Leonardo-Fuse (along illumination)

Given two opposing illumination sources *a* and *b*, image fusion in SPIM is to reconstruct a single stack 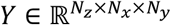with high image quality from two inputs 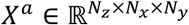 and 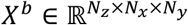 with *N*_*z*_ slices of size *N*_*x*_ × *N*_*y*_. To do so, image fusers are to quantify local saliency levels of the images and select or give more importance to pixels that have more structures/less blur. Here, Leonardo-Fuse (along illumination) moreover incorporates prior knowledge of SPIM and sample’s orientation, that is given a specific pixel at (z, *x, y*) in the image space, we are empirically in favor of 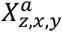 over 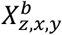, if illumination light travels through fewer scattering tissues from light source *a* to get to (z, *x, y*) than source *b*. Moreover, the degradation of image quality can be decomposed into two parts, one along illumination which is non-positive, and one along detection which is shared among *a* and *b*. Thus, using left-right illumination orientation as example and *a* and *b* being sources on the left and right, respectively, transition of trust on *X*^*a*^ to *X*^*b*^ can be rigidly defined as a series of curves associated with row indexes:

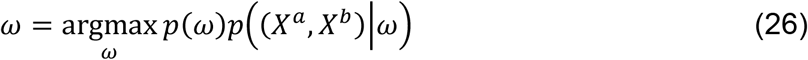

where 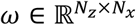is the image quality changeover, termed fusion boundary, with the *z*-th row corresponding to the curve fusion boundary in-between *z*-th slice of (X^*a*^, *X*^*b*^). After logarithm, Eq. (26) can be further decomposed:

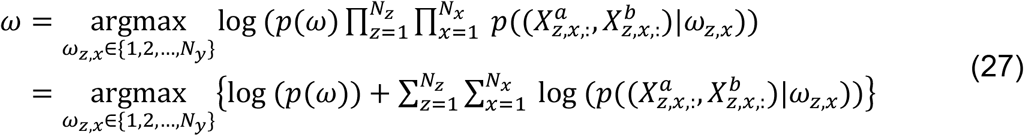

where *X*_*z*,x,:_ is the *x*-th row of *X* in the *z*-th slice, *ω*_z,x_ is the 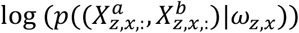 are defined as the smoothness of the fusion boundary *ω* and the row-wise saliency level, respectively.

Next, we define 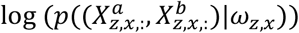, which is interpreted as the saliency level of the fusion result at the *x*-th row in the *z*-th slice, 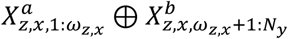, where ⊕ is the concatenating operator, given that image quality changeover is currently determined as *ω*_z,x_. We directly define row-wise saliency level as a summation of pixel-level saliency level measured using non-subsampled contourlet transform (NSCT) (Extended Data Fig. 5A). Here, NSCT is a multi-scale, multi-directional, and shift-invariant image decomposition that can capture critical transitions like edges and corners, and we perform NSCT slice-by-slice independently. In detail, we use 3, 3, and 3 directions in the scales from coarse to fine to get a series of band-limited depiction 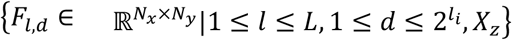 for the *z*-th slice in each SPIM input with *l*_*i*_ = 3 with *i* = 1,2, ⋯, *L* = 3. As a result, pixel-level saliency level is defined as:

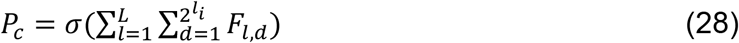

where *σ*(·) is a 3 × 3 standard deviation operator after NSCT coefficients at various levels and scales are summed up. Next, corresponding row-wise saliency level at the *x*-th row in the *z*-th slice conditioned on *ω*_z,x_, i.e., 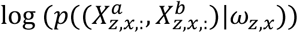 is defined as:

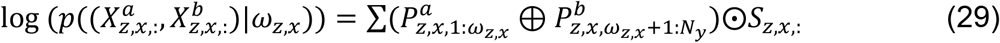

where *P*^*a*^ and *P*^*b*^ are pixel-level saliency level for respective stack, ⊕ is the concatenating operator, ⊙ is the element-wise multiplication, and *S* is a foreground segmentation mask we derived in advance (Extended Data Fig. 5A). Here, this segmentation mask is to incorporate a trustworthy visualization of illumination light propagating inside of the specimen. To segment, a combination of OTSU thresholding (calculated on the whole volume) and watershed is applied to every slice in each input *X*^*a*^ and *X*^*b*^. Since the segmentation results become less reliable as the illumination light diverges at deeper layers of the specimen, we derive the segmentation mask as the region between left and right boundaries extracted from *X*^*a*^ (illuminated from left-hand side) and *X*^*b*^ (lighted from right-hand side), respectively. We additionally encourage inter-slice continuity of the segmentation mask by using `remove_small_objects` and `remove_small_holes` along depth, two functions provided by skimage package in Python.

In parallel, inspired by the continuity of a biological specimen, Leonardo-Fuse (along illumination) considers smoothness of the fusion boundary *ω*:

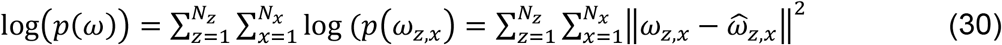

where 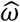 is the fusion boundary *ω* after window-based smoothing. In our previous publication^37^, we obtained 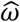via Savitzky–Golay (SL) filter, which was to fit subsets of adjacent data points with a low-degree polynomial by the method of linear least squares. Analytically, this can be implemented via convolving all data subsets with a single set of SL filter coefficients 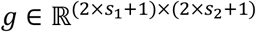. Thus, for *ω*_z,x_, given its neighboring set 𝒩(z, *x*) = {(z + *i, x* + *j*)| − *e*_1_ ≤ *i* ≤ *e*_1_, −*e*_2_ ≤ *i* ≤ *e*_2_}, its smoothed estimate is 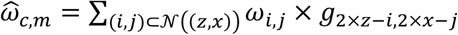. Although the SL filter can preserve the trend of *ω*, dynamic abruption related to changes in sample shape is not considered. Therefore, we reuse the segmentation mask 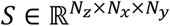, extract its left boundary 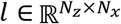 and right boundary 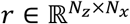, and re-define the neighboring set for the pixel at (z, *x*) as:

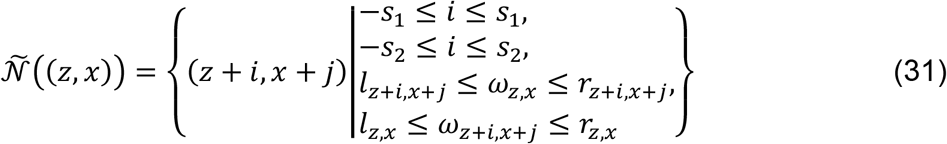

Correspondingly, smoothed 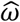 is changed as:

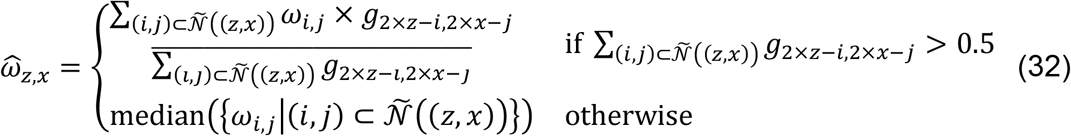

where smoothing is performed and then normalized based on the neighboring set 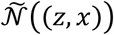. For the sake of stabilization, SL filter is only applied to *ω*_z,x_ accompanied with a relatively large 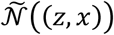, whereas median filter, i.e., median(⋅), is adopted for the rest.

As a result, the overall energy function to be optimized by Leonardo-Fuse (along illumination) is:

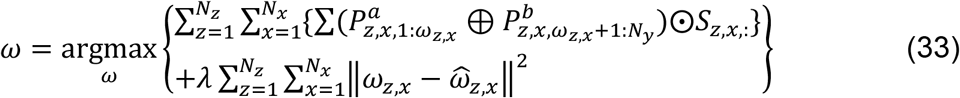

where *λ* is the trade-off parameter. To optimize, Leonardo-Fuse (along illumination) uses the Expectation–Maximization (EM) algorithm. Specifically, in the *k*-th iteration, *ω*^(k)^ is solved by iterating over {y|1 ≤ *y* ≤ *N*_*y*_} to maximize Eq. (33), meanwhile with 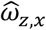 substituted by 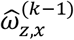 from the previous iteration. Iteratively, 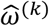 is updated based on Eq. (32). Additionally, *ω*^(0)^ is initialized by maximizing Eq. (29), followed by a median filter. Moreover, since the specimen may not occupy the entire image space, several entities in *ω*^(0)^ may be missing, which means there is no pixel detected as sample-relevant after segmentation along this specific row. Missing values are filled using a first-order polynomial interpolation.

Next, after a total of *K* iterations of EM, two image stacks can be stitched based on the fusion boundary *ω*^(*K*)^, which will be referred to as *ω* in the following for the sake of simplification. However, the smoothed fusion boundary *ω* may not fit specimen details nicely. Thus, we reuse GF^34^, which we already described in the Leonardo-DeStripe subsection to refine the weight maps and enhance correlations between neighborhood pixels—spatial consistency (Extended Data Fig. 5B). Specifically, two binary weight maps are directly derived out of the fusion boundary *ω* for *X*^*a*^ and *X*^*b*^ respectively:

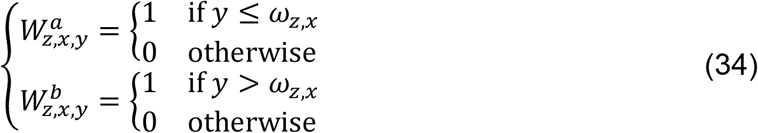

where 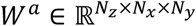and 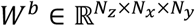are the weight maps for sources *a* and *b*, respectively. Next, each weight map *W* is refined using GF with the corresponding source image *X* serving as the guidance image:

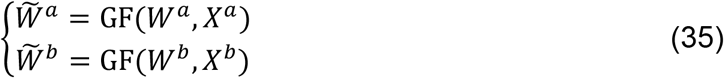

where linear transformation is now assumed within a 3D window rather than a 2D one, such that both intra- and inter-slice spatial consistencies can be considered simultaneously. Additionally, values of the two weight maps are normalized such that they sum to one at every pixel position. As a result, source images with opposing illumination orientations are fused together by weighted averaging:

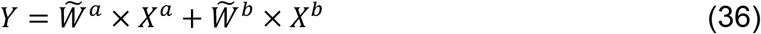

### Leonardo-Fuse (along detection)

Basically, fusing two image stacks with opposing detection devices and a shared illumination source is equivalent to fusing two datasets illuminated from opposing orientations but acquired using the same detecting camera. In other words, suppose *X*^*a*^ and *X*^*b*^ are illuminated via the same light source but recorded from opposite directions, Leonardo-Fuse (along detection) is to estimate a fusion boundary 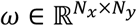whose (x, *y*)-th entity is the quality changeover between 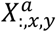 and 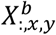. Thus, in a complete workflow where dual-sided illumination and dual-sided detection are deployed and outputs four stacks in total, the first step in Leonardo-Fuse (along detection) is to perform fusion boundary estimation four times (Extended Data Fig. 9A):

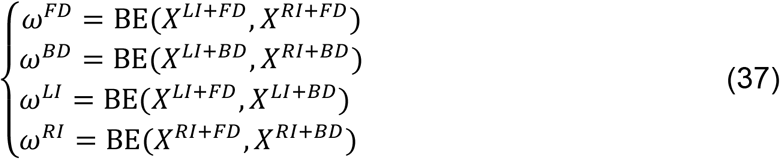

where BE(·) is the method of fusion boundary estimation that we already described in the Leonardo-Fuse (along illumination) subsection, superscript *LI, RI, FD* and *BF* denote illumination from the left side, illumination from the right side, acquired from the front side, and acquired from the back side, respectively.

Next, four fusion boundaries can be binarized as weight maps *W*^*FD*^, *W*^*BD*^, *W*^*LI*^ and *W*^*RI*^, respectively, similar to Eq. (34). The weight maps need to be integrated before serving as criteria for dataset stitching, as the process of estimating corresponding boundaries is done in parallel. Specifically, there potentially can be two scenarios where weight maps may conflict/mismatch each other (Extended Data Fig. 9B). First, one region is chosen by two weight maps simultaneously. It is noteworthy that this happens only between two datasets with both different illumination directions and opposite detection manners, i.e., between *X*^*LI*+*FD*^ and *X*^*RI*+*BD*^, or *X*^*RI*+*FD*^ and *X*^*LI*+*BD*^. Second, a region is not covered by any weight map. In both cases, we re-quantify the NSCT-based saliency level provided by different illumination-detection combinations, sum over the whole region, merge the region into the weight map corresponding to the highest saliency level. Additionally, in case that inter-slice weight maps may be discontinuous, we apply `remove_small_objects` and `remove_small_holes` along depth. As a result, GF-based weight map refinement, followed by dataset stitching, can be performed, similar to Leonardo-Fuse (along illumination).

### Sample preparation

#### Zebrafish

Adult zebrafish (*Danio rerio)* and embryos were maintained following established protocols^38^, in compliance with EU directive 2010/63/EU and the German Animal Welfare Act. The transgenic line Tg (H2B:GFP) was used, where GFP is expressed ubiquitously in all nuclei. The stained in vivo zebrafish embryos at 1 dpf were incubated in Bodipy (20 µl stock solution in 980 µl E3, Invitrogen) for 1 h before embedding and imaging. Zebrafish embryos were kept at 28.5°C in E3 medium with 0.003% 1-phenyl-2-thiourea (PTU) to suppress pigmentation. Before imaging, embryos were selected for their expression pattern and checked for potential malformations using a fluorescence stereomicroscope. Zebrafish embryos (<120h) are mounted in Fluorinated ethylene propylene (FEP) tubes for *in vivo* light sheet imaging. FEP tubes (ProLiquid; refractive index 1.338, inner diameter 0.8 mm, outer diameter 1.2 mm) were straightened by heating in an oven at 180°C for 2 hours, then cooled slowly to room temperature. Tubes were flushed with isopropanol, followed by double-distilled water (ddH_2_O), and immersed in 0.5 M NaOH. They were ultrasonicated for 10 minutes, rinsed again with ddH_2_O, and flushed with 70% ethanol. Tubes were then placed in fresh ddH_2_O and ultrasonicated for another 10 minutes. Finally, they were flushed and stored in ddH_2_O. Gloves and/or forceps were used to handle the cleaned tubes. Low-melting-point agarose (Sigma) was prepared in E3 medium. For embedding in 1.5% low-melting agarose, embryos were suspended in liquid agarose and gently drawn into FEP tubes using a blunt needle and syringe.

#### Tissue clearing

The murine hind legs, murine hearts, and zebrafish brain tissue were optically cleared with ethyl cinnamate (ECi, Sigma Aldrich, Cat. 112372-100G) as previously described^28,39^. Briefly, samples were fixed with 4% PFA/PBS and dehydrated with an ascending ethanol series (50%, 70% and twice in 100% EtOH) gently shaking at room temperature for 12 hours per step.

The fully dehydrated samples were transferred to ECi and gently shaken at room temperature until they were optically transparent. The fully dehydrated samples were transferred to ECi and gently shaken at room temperature until they were optically transparent.

### Data acquisition

#### SPIM Data

Images were acquired with a home-built light sheet system developed in the Huisken lab. It’s hardware and software components are standardized and interchangeable between different configurations^40^. We used a Toptica iChrome CLE fibre-coupled laser with 488nm excitation laser, Nikon water immersion lenses with 10x/0.3 and 16x/0.8 as illumination and detection lenses, Physik Instrumente (PI) compact linear stages for sample positioning and a PCO pco.panda sCMOS camera to collect the fluorescence signal. Two light sheet configurations were employed, a T-SPIM setup with two illumination arms and one detection arm and an X-SPIM setup with a second, opposing detection arm. The X-SPIM configuration allows simultaneous image stack acquisition using two cameras. Both systems create a static light sheet using a cylindrical lens in the illumination arm. A volumetric z-stack is created by moving the sample through the light sheet (488 nm excitation, band-pass 525/50 nm emission filter, 25 ms exposure time, 40 frames/s, 2.5 µm z-steps). Sequential, two-sided illumination is done with a scan mirror that also allows for multi-directional illumination by pivoting the light sheet (mSPIM)^41^. Imaging was performed on live zebrafish embryos between one and four days post-fertilization (dpf). The custom imaging chamber was filled with E3 fish medium. Images were 16-bit TIFF stacks for each view and every illumination and detection pair. In an X-SPIM configuration, this results in four data sets. Cleared zebrafish brain and murine heart samples were imaged on a modified version of this instrument utilizing Nikon 4x/0.13 air lenses and a quartz cuvette to hold the clearing media and sample.

#### Blaze data

The optically cleared samples were imaged using a LaVision BioTec Blaze ultramicroscope (Miltenyi/LaVision BioTec, Bielefeld, Germany) with a supercontinuum white light laser providing excitation from 460-800 nm and 7 individually combinable excitation and emission filters defining excitation and emission profiles from 450 nm to 865 nm. In addition, an AndorNeo sCMOS camera with a pixel size of 6.5µm2 and a 4x objective (NA 0.35) with a magnification changer set to 0.6x was used for image acquisition.

The optically cleared samples were immersed in ECi in a quartz cuvette and excited at 560/40 nm to visualize tissue autofluorescence and detected with a 620/60 nm bandpass emission filter at a Z-step of 4 µm.

## Extended Data Figures

**Extended Data Fig. 1.**
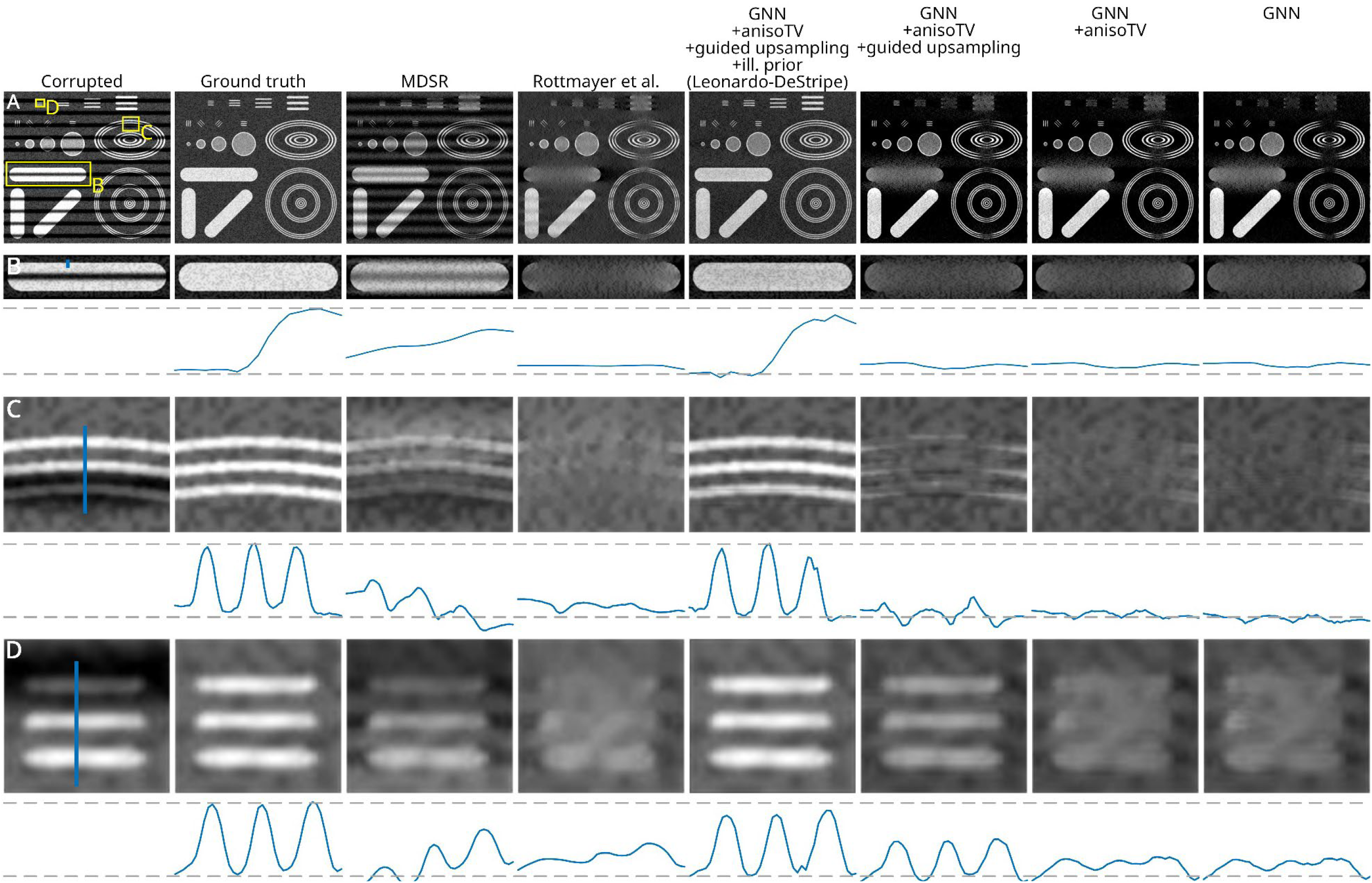
Comparison between existing methods, Leonardo-DeStripe, and ablation studies on a simulated spherical phantom image. (A) Two benchmark methods, MDSR and Rottmayer et al., are shown. MDSR tends to leave stripe artifacts partially, while Rottmayer et al. often suppresses both the stripes and underlying sample structures. In contrast, Leonardo-DeStripe effectively removes stripes while preserving sample structures to the greatest extent. To assess the contribution of each component in Leonardo-DeStripe, we performed ablation studies. The graph neural network (GNN) plays a central role in stripe removal, although it may also compromise fine structural details. The anisotropic total variation unit (anisoTV) helps in preserving sample structures, though its effect is sometimes minor. The guided upsampling module, and the post-processing module for sample structure preservation (denoted as ill. prior), are applied on top of the trained GNN and achieve the best overall structure preservation. Specifically, the guided upsampling module helps maintain sample structure in (D), while the ill. prior in (B–D) consistently preserves sample features of various sizes and orientations with a significant margin.

**Extended Data Fig. 2.**
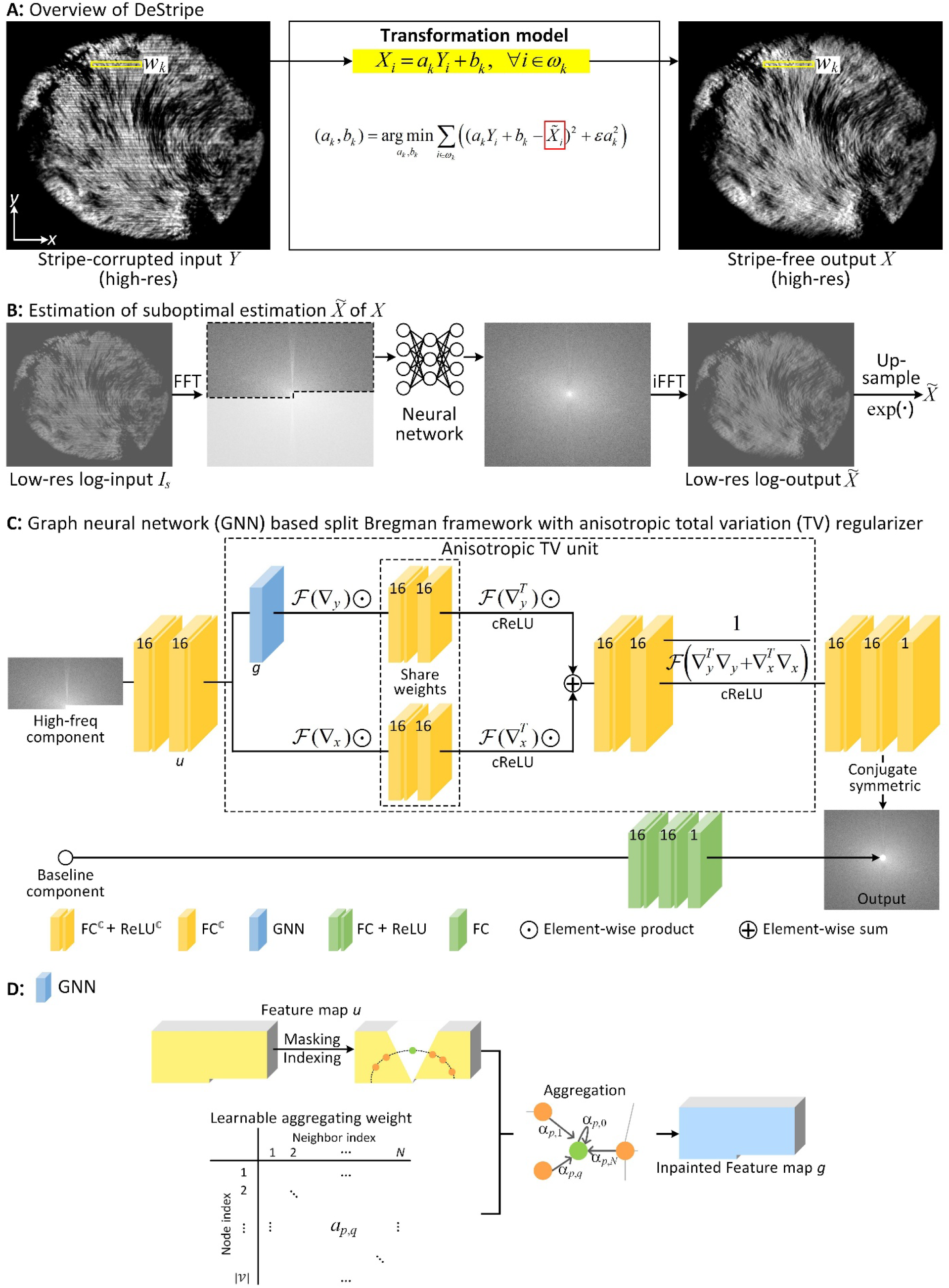
Architecture of Leonardo-DeStripe. (A) Overall, Leonardo-DeStripe treats the stripe removal task in SPIM as mapping stripe-corrupted input to stripe-free output via a local linear transformation, supported by the observation that stripes are considered locally consistent along illumination direction. (B) The guidance map used by the local linear transformation, which can be low-resolution and even detail compromised, is learned via a deep neural network (NN) in log space after Fourier transformation, where multiplicative stripes are converted into additive noise and thus easier to be modeled. (C) Proposed NN follows a split Bregman framework regularized via anisotropic total variation (TV) in latent feature space. Specifically, a graph neural network in (D) is nested, in which stripe-corrupted Fourier coefficients are inpainted as a combination of their neighbors on a polar coordinate system.

**Extended Data Fig. 3.**
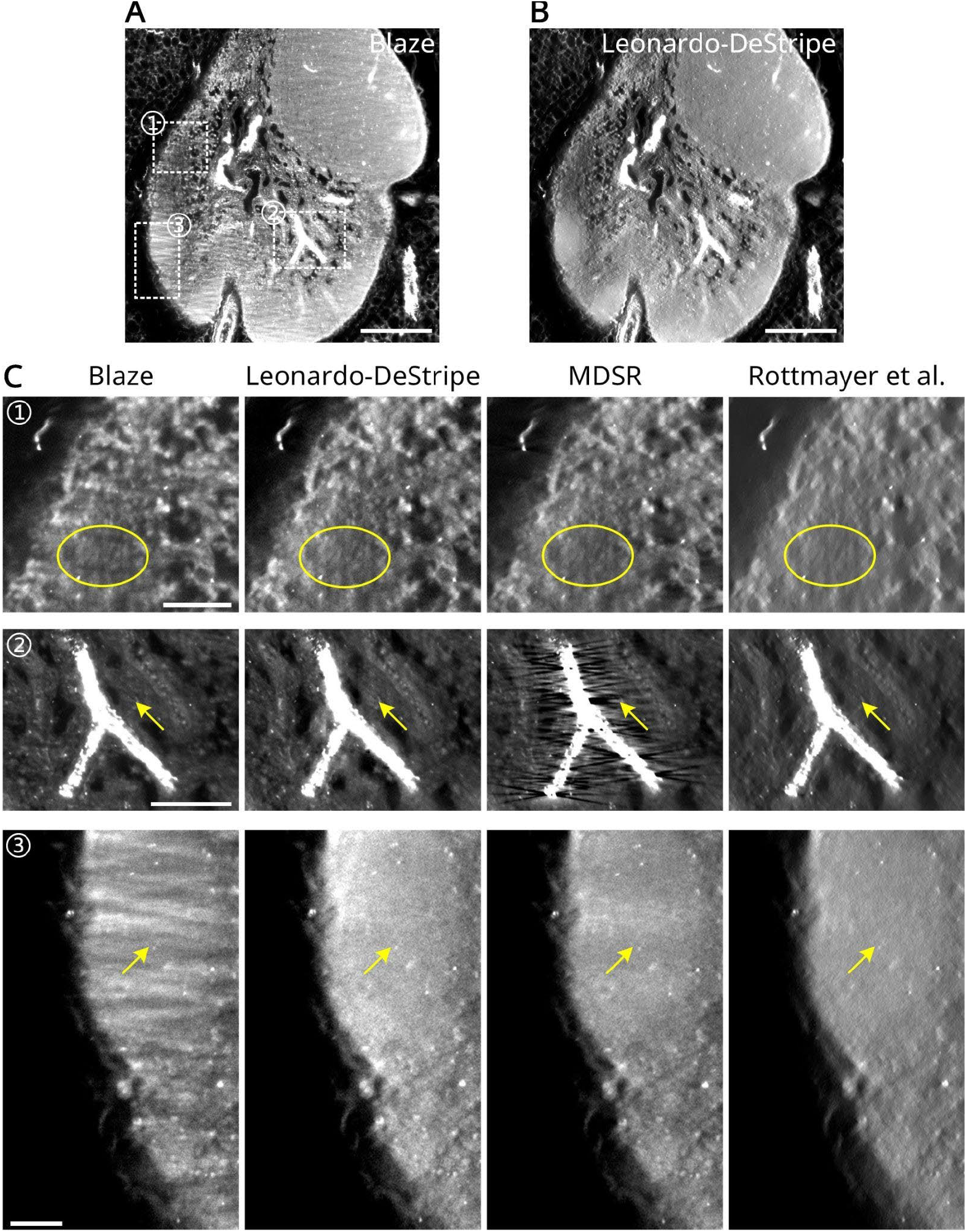
Comparison on a mouse stained with antibodies against the TH antibody in (A). (B) Leonardo-DeStripe consistently resolved stripes of all three directions. (C) In comparison, MDSR left residual stripes (arrows in patch ➂), or introduced additional artifacts (arrows in patch ➁). Meanwhile, Rottmayer et al. removed sample details (circles in patch ➀). Scale bars, 200 μm (A, B), 50 μm (C).

**Extended Data Fig. 4.**
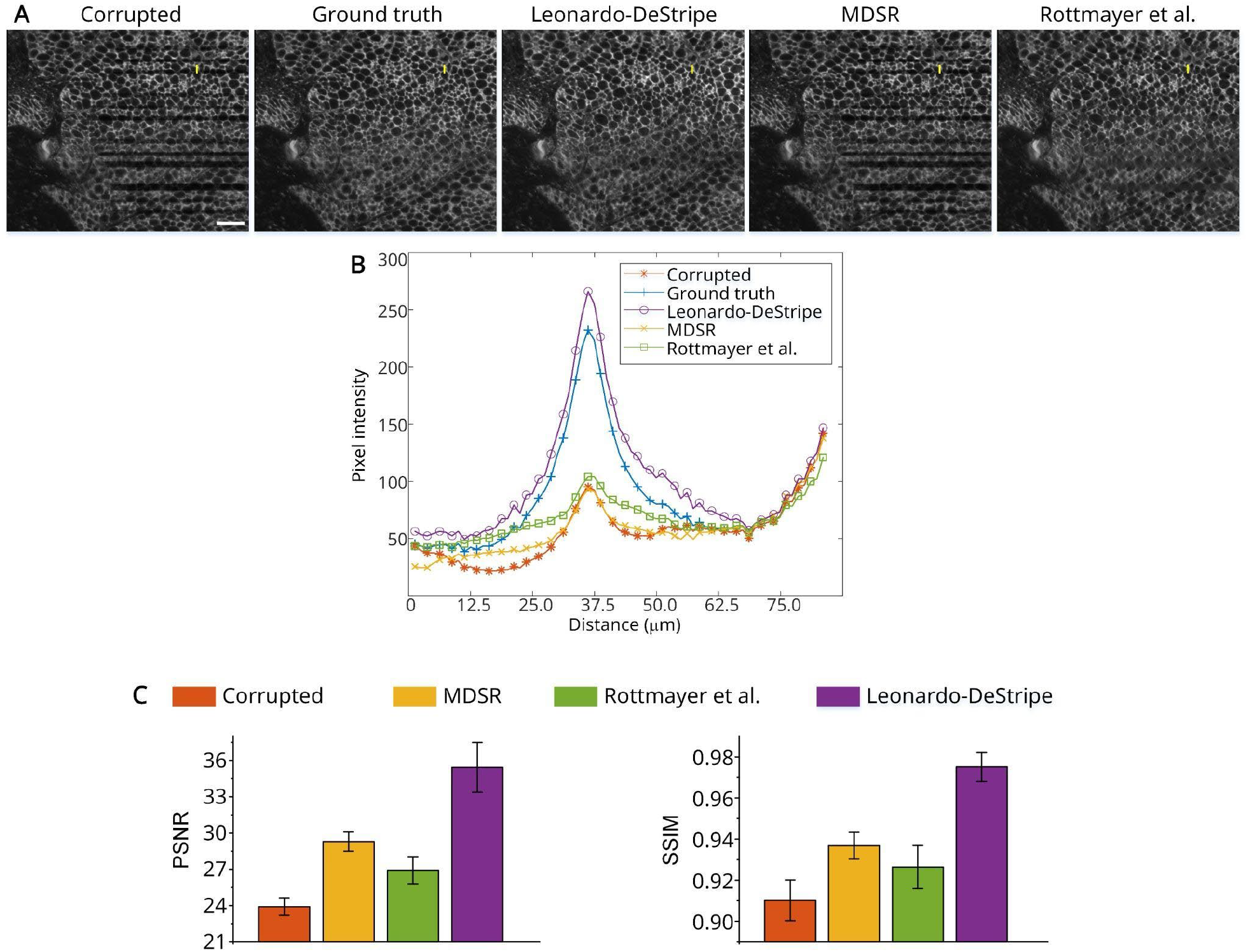
Simulation results in adipose tissue. A stripe-free stack of adipose tissue with 95 × 2076 × 1728 voxels was used as the ground truth, onto which synthetic stripes were simulated. (A) Leonardo-DeStripe produces results that closely match the ground truth, whereas MDSR tends to merge thin stripes into thicker ones rather than effectively removing them and Rottmayer et al. leaves residual stripes. (B) Intensity profiles along the yellow line segments in (A) show that Leonardo-DeStripe more faithfully recovers the ground truth compared to MDSR and Rottmayer et al. (C) Quantitative evaluation demonstrates that Leonardo-DeStripe outperforms the benchmark methods. Means and standard deviations from the total 95 slices are shown for PSNR and SSIM. Scale bars, 400 μm (A).

**Extended Data Fig. 5.**
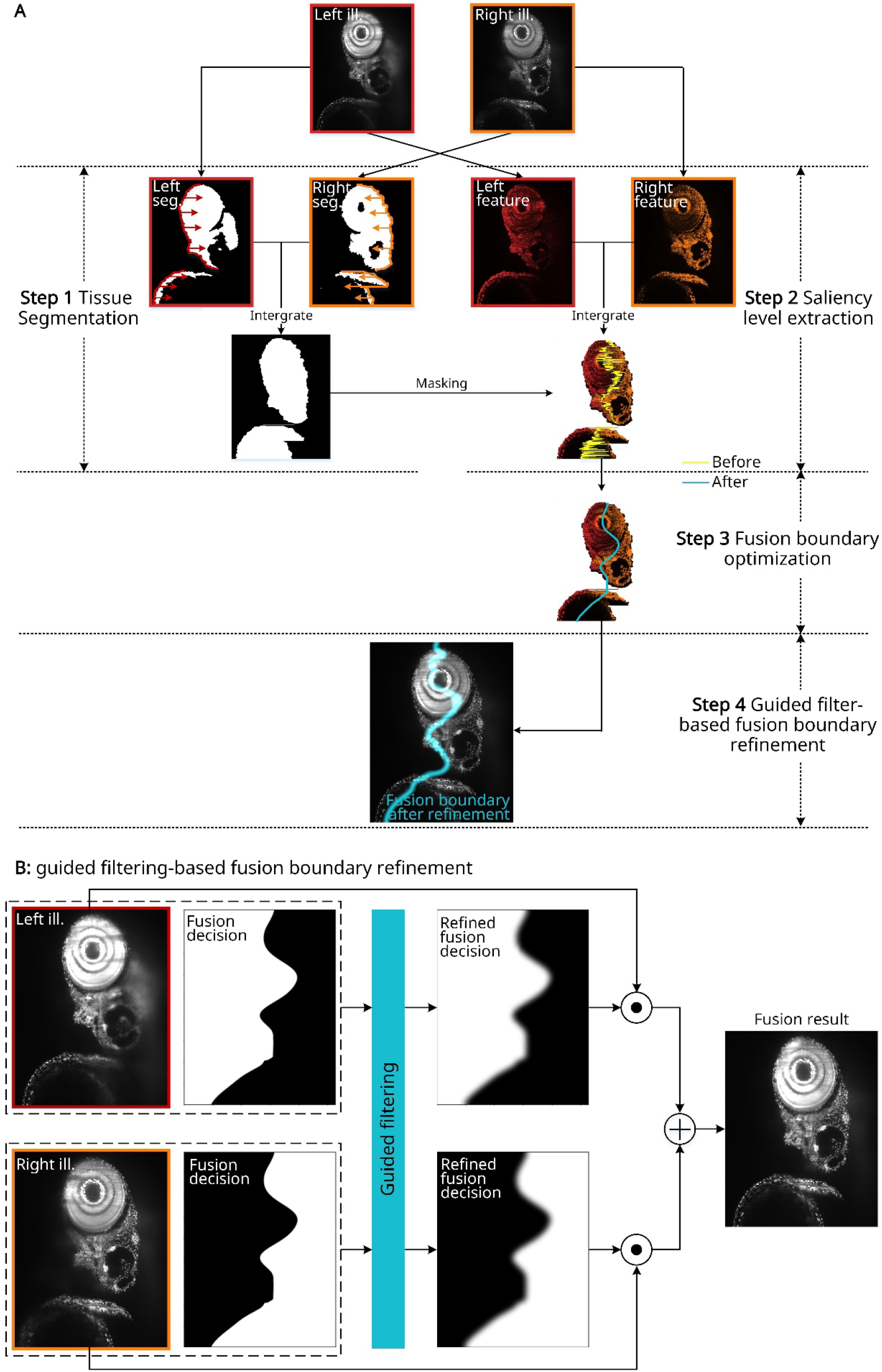
Architecture of Leonardo-Fuse (along illumination). (A) In parallel, Leonardo-Fuse (along illumination) segments sample-related pixels and simultaneously quantifies pixel-level image qualities using non-subsampled contourlet transform (NSCT). The mask used to estimate photon propagating path is therefore integrated as regions between left and right boundaries extracted from left seg. and right seg., respectively. Only sample-related saliency levels are then fed into an expectation-maximization algorithm to estimate the fusion boundary, where the smoothness of the fusion boundary, together with the quality of the fused result, are considered jointly. Before stitching the two SPIM inputs based on the fusion boundary, Leonardo-Fuse (along illumination) additionally refines the fusion boundary using guided filtering (GF) to ensure smooth transitions near the fusion boundary while maintaining distinct selections from each stack in areas farther away. The process of refining the fusion boundary using GF is explained in detail in (B).

**Extended Data Fig. 6.**
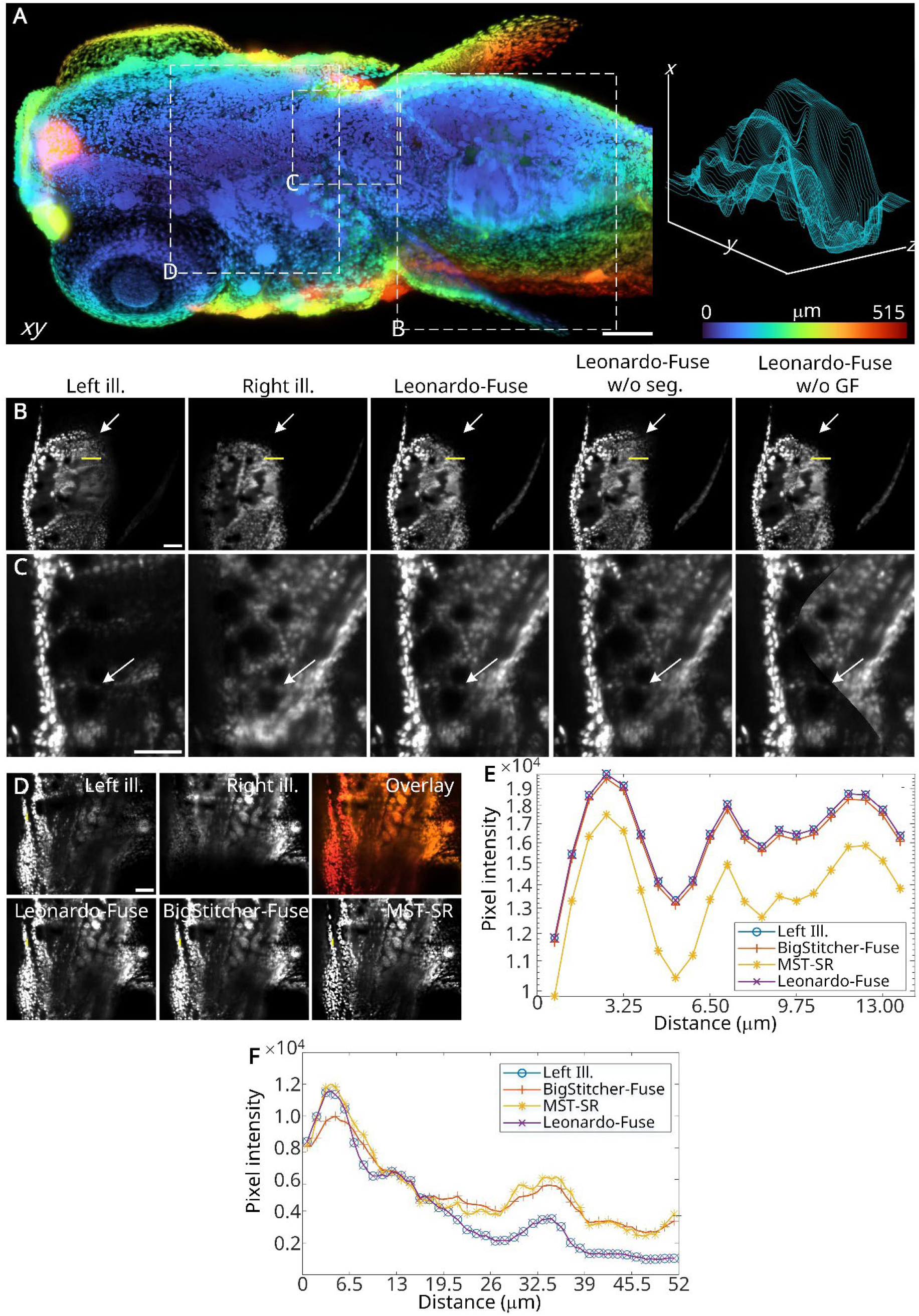
Ablation studies of Leonardo-Fuse (along illumination). (A) A depth-colored maximum intensity projection from Leonardo-Fuse (along illumination) applied to an H2B-GFP-expressing transgenic zebrafish, together with the estimated fusion boundary at various depths plotted on the right. (B) The consideration of prior knowledge of light travel path (enabled by the tissue segmentation) helps Leonardo-Fuse (along illumination) better reject ghost artifacts (white arrows in Leonardo-Fuse w/o seg. panel). Meanwhile, the guided filtering (GF)-based fusion boundary refinement in (C) allows us to seamlessly stitch multiple datasets without creating artifacts (white arrows in Leonardo-Fuse w/o GF panel). (D) Information is integrated by using both baseline methods and our method. (E) Intensity profiles of segments indicated by yellow lines in (D) are given, where Leonardo-Fuse (along illumination) matches the good input (Left ill.) perfectly. (F) Line plot extracted from the region indicated by yellow lines in (B). Only Leonardo-Fuse (along illumination) allows the best data fidelity (perfectly overlaps with Left ill.). Scale bars: 100 μm (A), 50 μm (B-D).

**Extended Data Fig. 7.**
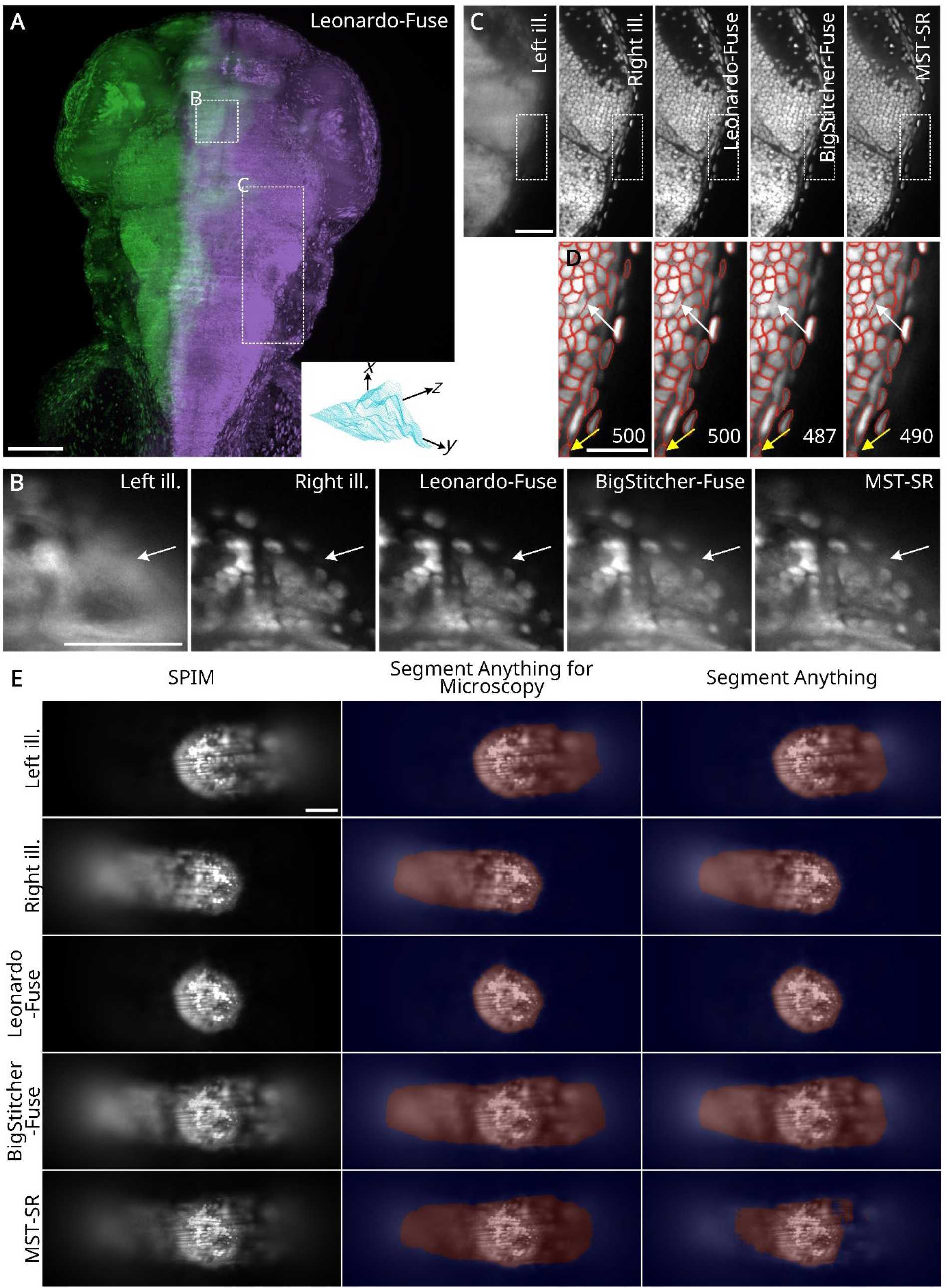
Leonardo-Fuse (along illumination) facilitates the downstream segmentation task. (A) Leonardo-Fuse (along illumination)’s result on a calcium imaging specimen is shown, together with the estimated fusion boundary. Two zoom-in regions are given in (B) and (C), where benchmark methods, especially MST-SR, suffer from halo artifacts (white arrows in (B)). (D) Therefore, when counting cell numbers using Cellpose, only Leonardo-Fuse (along illumination) gives the exact same result and thus the best data fidelity as input with illumination source placed on the right. In comparison, baseline approaches fail to detect several cells (white and yellow arrows for BigStitcher-Fuse and MST-SR, respectively). (E) Leonardo-Fuse enables accurate representation of the outline of a 1 dpf zebrafish labeled with Bodipy. Due to ghost artifacts, both the Segment Anything model and the Segment Anything for Microscopy variant fail to capture the correct outline when using reconstructions from BigStitcher-Fuse or MST-SR. In contrast, Leonardo-Fuse provides a reconstruction that allows both models to accurately recover the zebrafish boundary. Scale bars: 100 μm (A, E), 50 μm (B, C), 25 μm (D).

**Extended Data Fig. 8.**
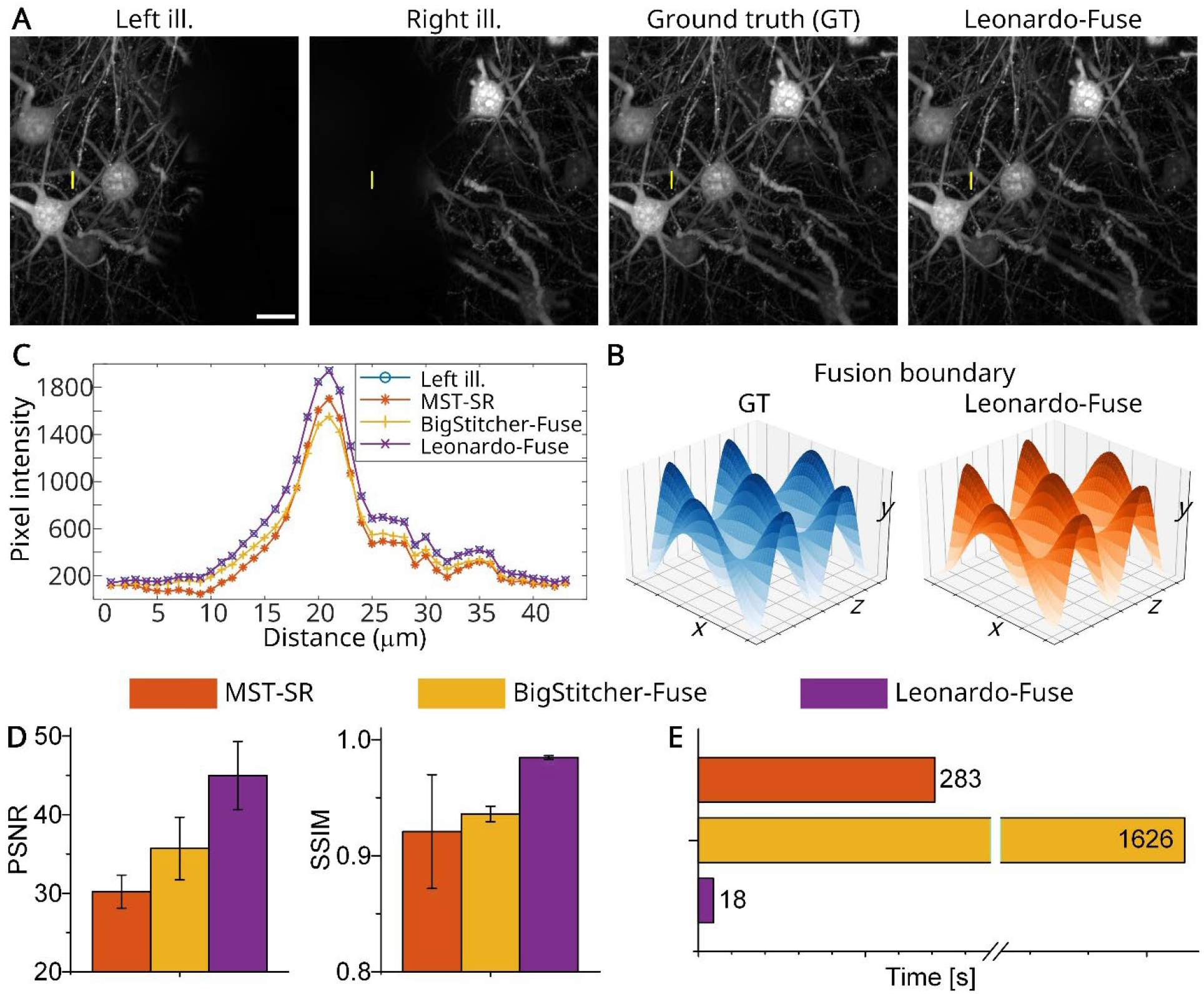
Leonardo-Fuse (along illumination) quantitatively outperforms benchmark methods. (A) Degradation along illumination is simulated on a previously published PEGASOS-cleared mouse brain with 158 × 512 × 512 voxels. (B) Leonardo-Fuse (along illumination) successfully learns the fusion boundary, very much resembling the ground truth (GT). (C) A line plot of a segment (indicated as yellow line in (A)) is given, where only Leonardo-Fuse (along illumination) preserves data fidelity (perfectly overlaps with left ill.). (D) Quantitative comparison. Means and standard deviations from the total of *N* = 158 slices are shown for PSNR and SSIM. (E) Leonardo-Fuse is significantly faster than the benchmark methods, requiring only 18 seconds to process the entire volume. Scale bars: 100 μm (A).

**Extended Data Fig. 9.**
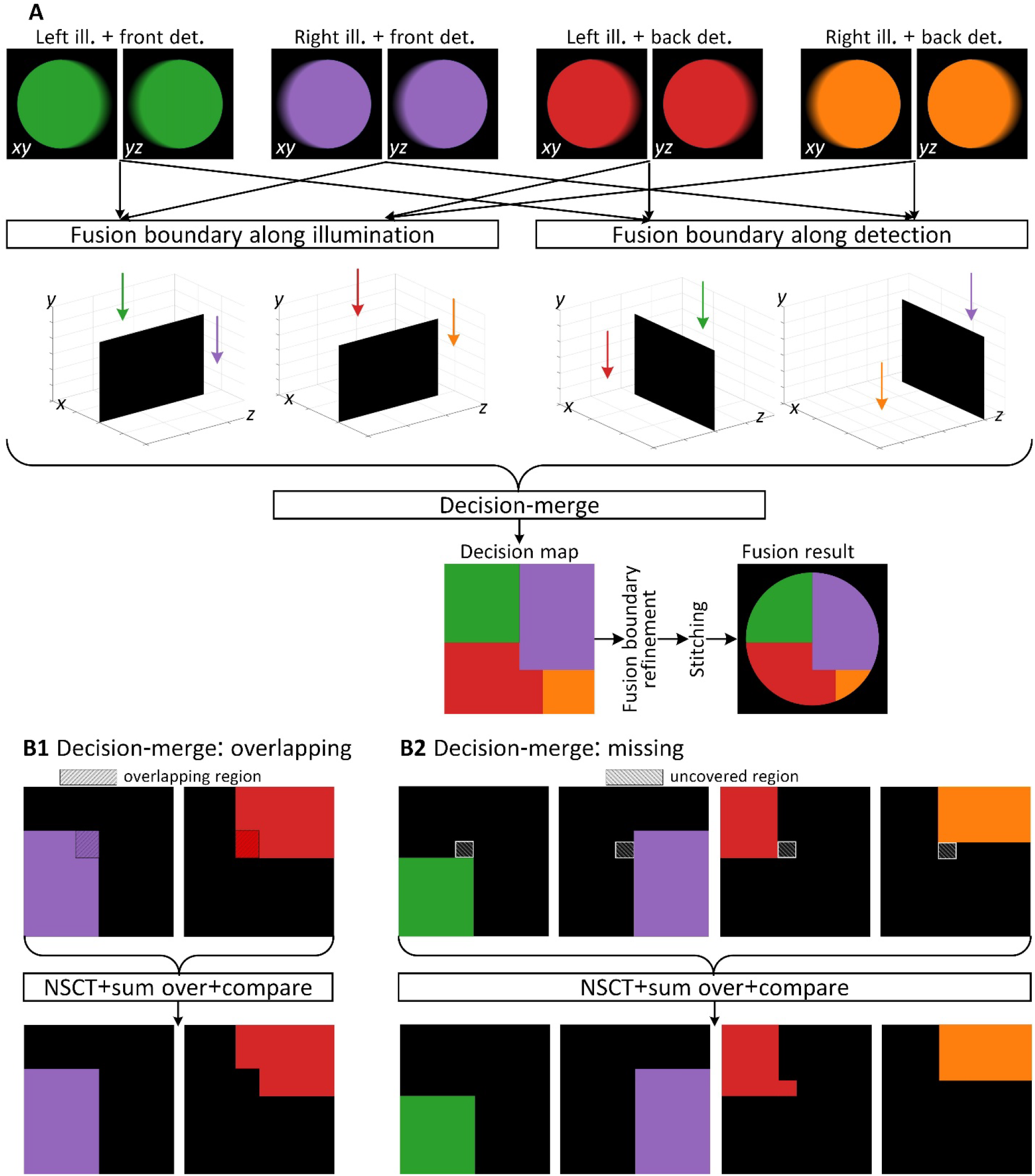
Architecture of Leonardo-Fuse (along detection). (A) Leonardo-Fuse (along detection) reuses the estimation of the fusion boundary four times, two of which along the illumination direction and the others along the detection direction. Since these four “good”-to-”bad” quality boundaries are estimated independently, they need to be merged into one before being fed into a fusion boundary refinement module and used as guidance for dataset stitching. Decision merge includes two scenarios: (B1) when the same region is covered twice and (B2) when a patch is not considered as “good” by any dataset.

**Extended Data Fig. 10.**
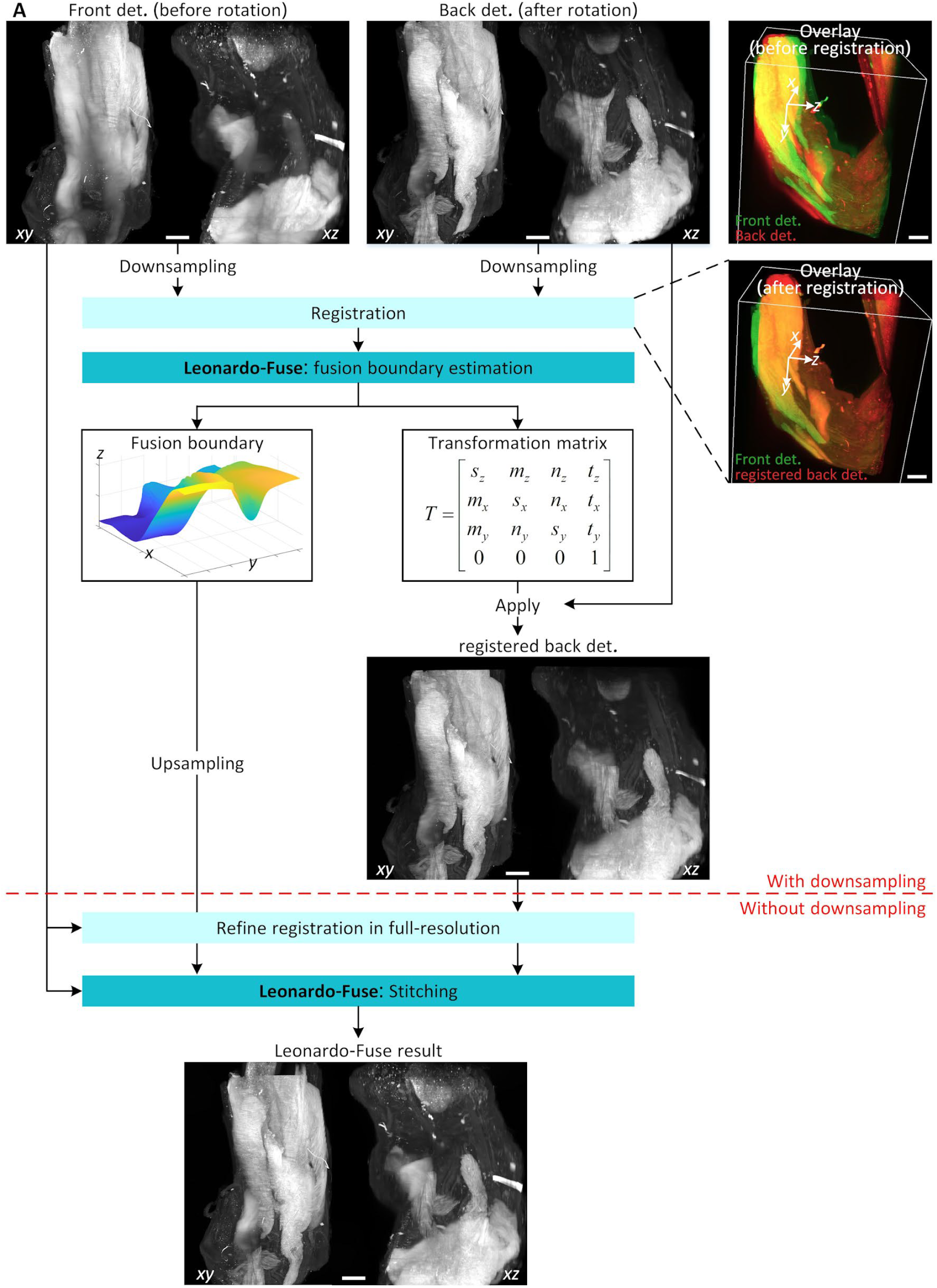
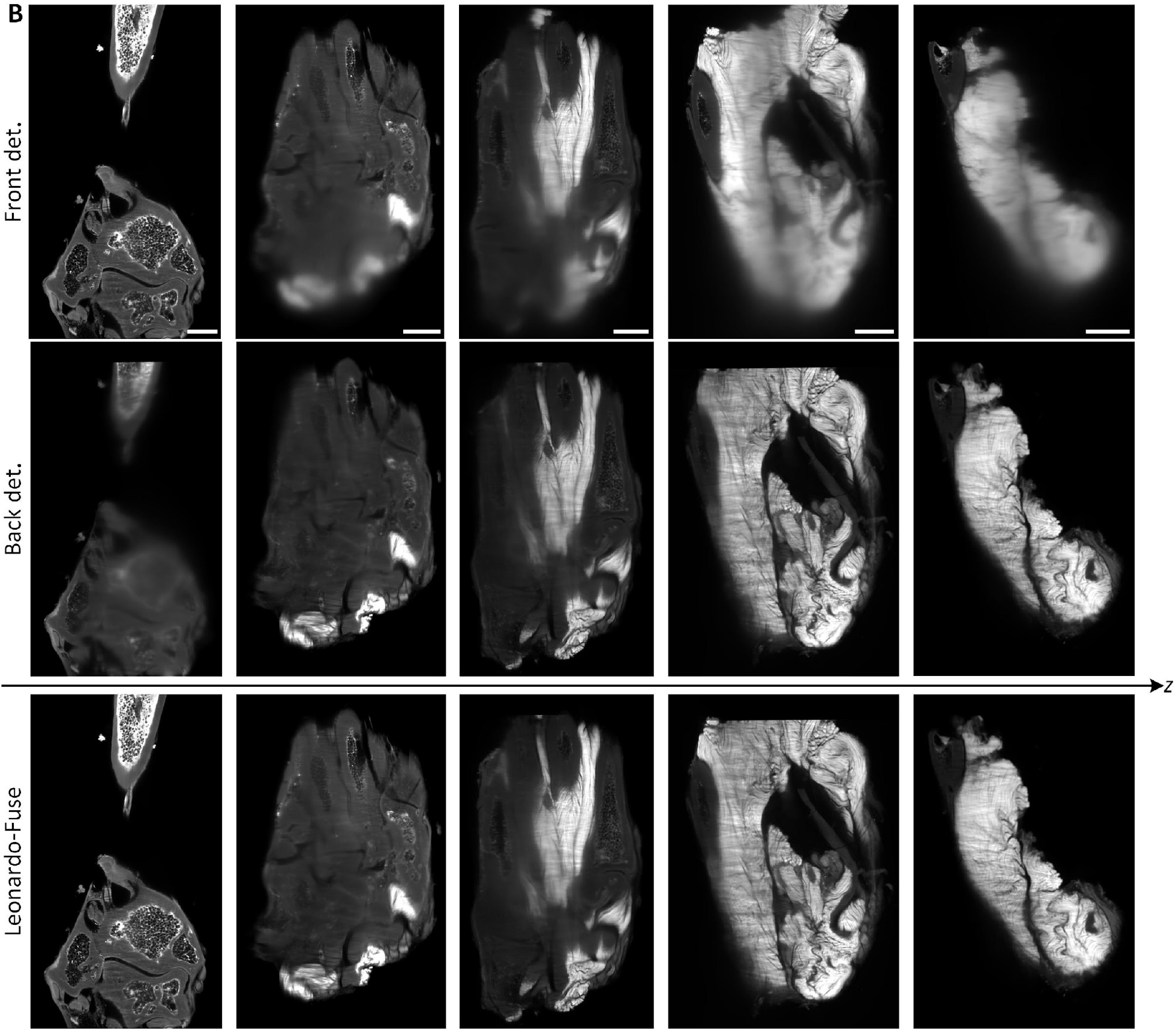
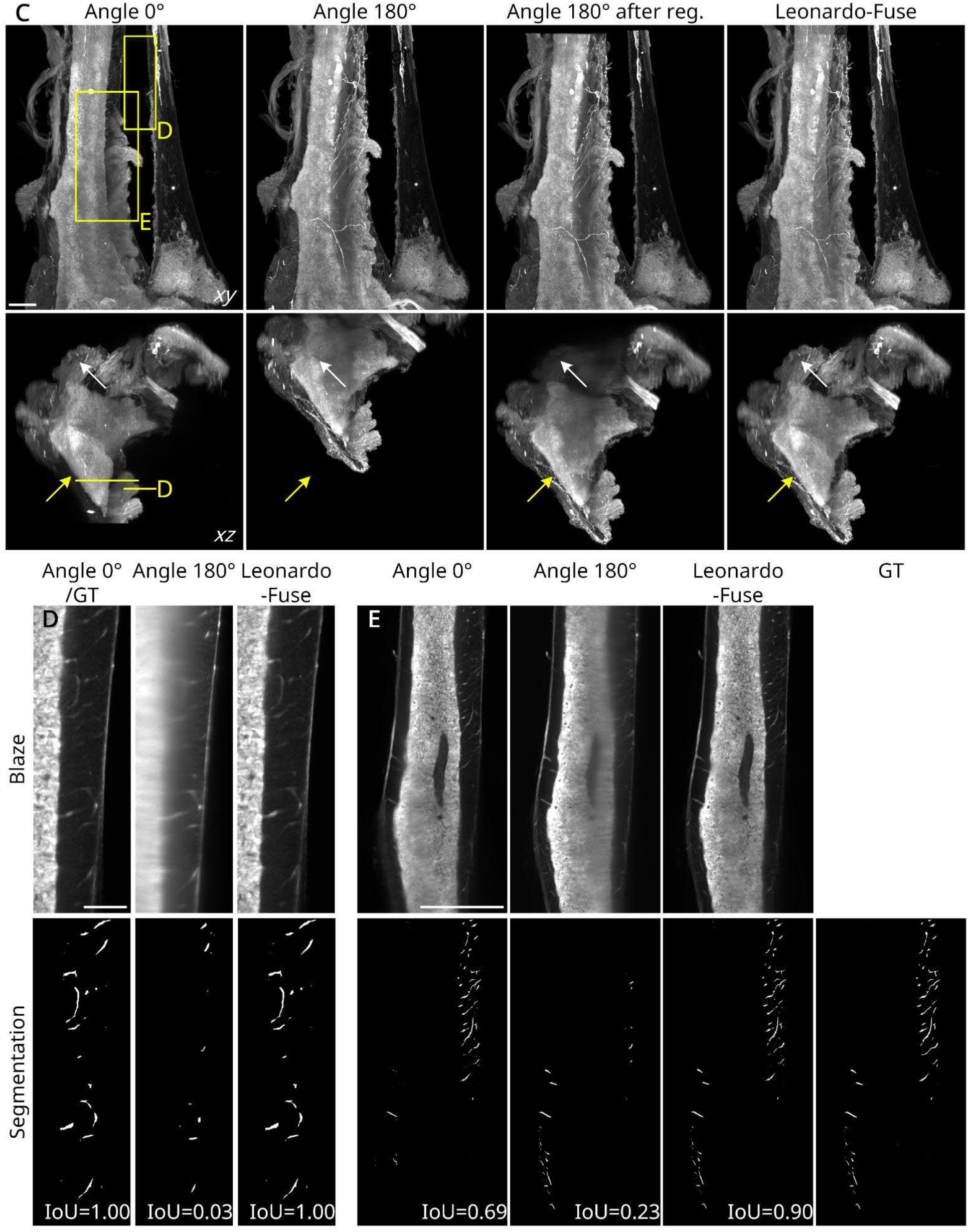
Leonardo-Fuse performs fusion on extremely large specimens using manageable computational requirements. (A) Leonardo-Fuse can be optimized to fuse extremely large tissues by estimating the fusion boundary, together with the transformation matrix, in a down-sampled space, and later refining the registration matrix at the original resolution and applying a stitching-based fusion technique in high resolution. (B) Leonardo-Fuse gradually transfers attention from front volume to back stack as it gets deeper into the tissue. (C) A murine tibia dataset, which is also shown in Fig. 5G-I. The data were acquired using an Ultramicroscope Blaze equipped with a rotatable sample holder, enabling imaging from two opposing angles by rotating the specimen 180°. Leonardo-Fuse first registers the rotated dataset (angle 180°) to the one before rotation (angle 0°), followed by fusion. Complementary information is successfully integrated: informative regions (white arrows) are transferred from angle 0°, while sharp features from angle 180° after registration (yellow arrows) are also preserved in the fused result. (D–E) Trans-cortical vessels (TCVs) segmented using a vesselness filter. In (D), Leonardo-Fuse achieves segmentation nearly identical to that of the higher-quality acquisition (angle 0°), as indicated by the Intersection over Union (IoU) values. In (E), where both inputs are partially degraded, Leonardo-Fuse effectively integrates complementary data to produce a segmentation most consistent with the manually annotated ground truth (IoU = 0.90 vs. 0.69 and 0.23). Scale bars: 500 μm (A-C).

