## Supplementary Materials for "Leonardo: a toolset to correct sample-induced artifacts in light sheet microscopy images"

### Supplementary Figures

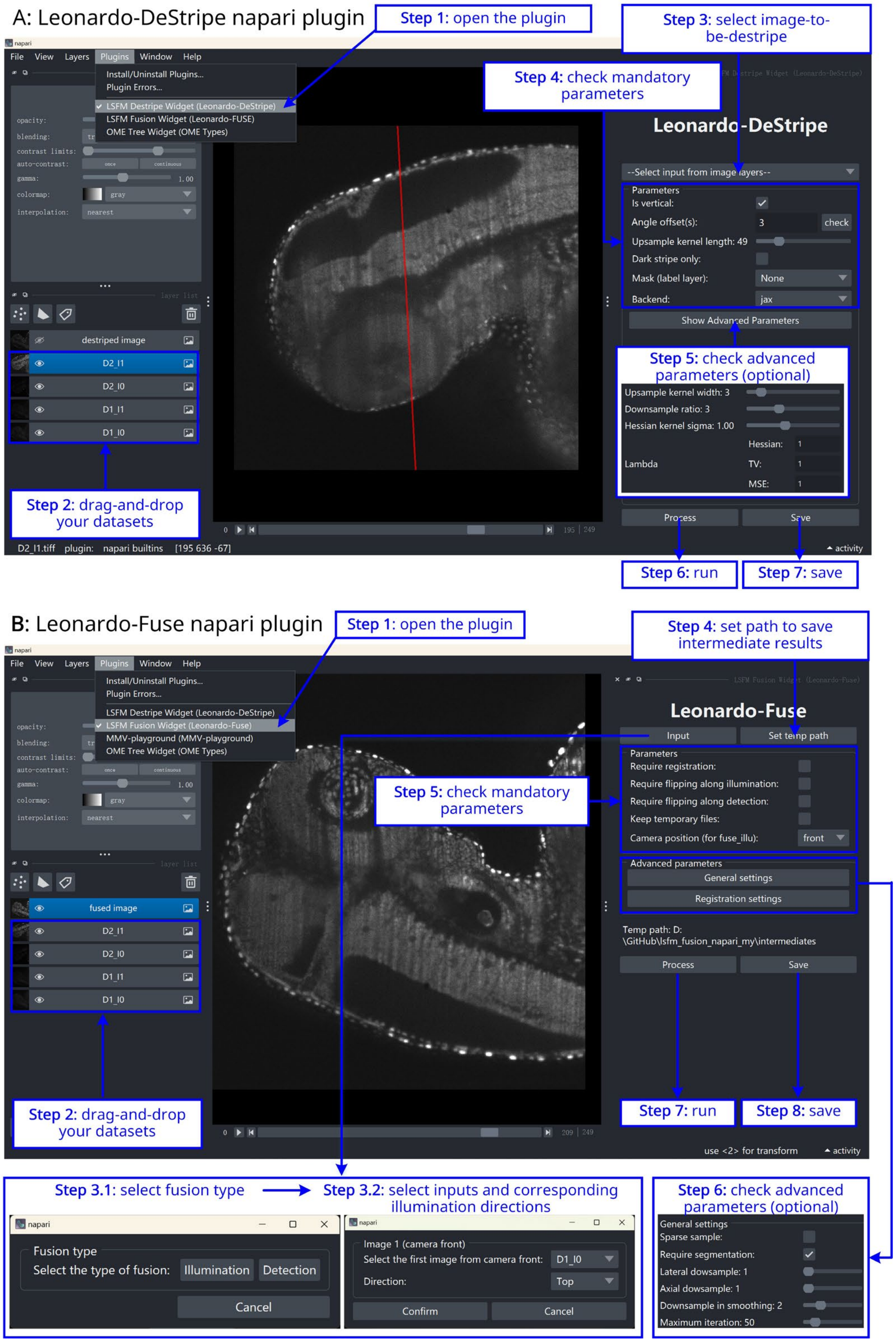

**Supplementary Fig. 1. Graphical user interface (GUI).** Leonardo is modular and encapsulated individually in Napari and can adapt to different SPIM variants and workflows. Specifically, Leonardo-DeStripe and Leonardo-Fuse are nested in two Napari widgets separately.

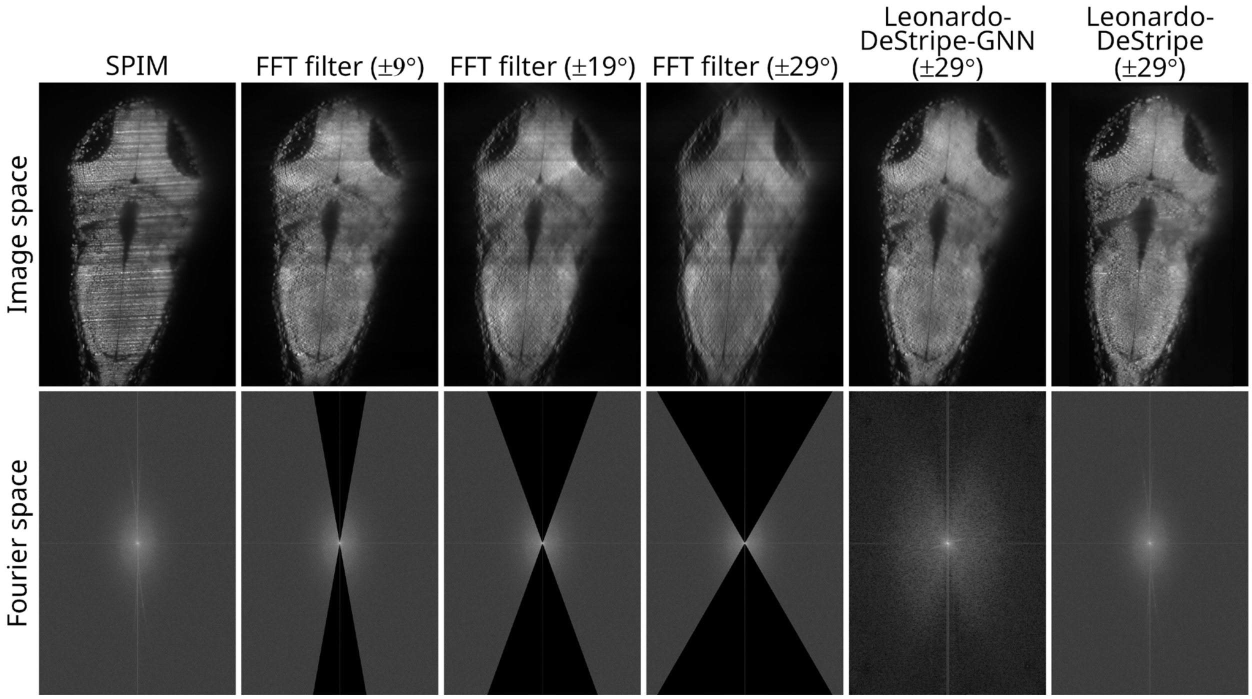

**Supplementary Fig. 2. A comparison between Leonardo-DeStripe and a conventional 2D band-pass FFT filter with different angular coverages of the wedge-shaped mask.** From left to right, the angular coverage of the masking region in the FFT filter increased from $\pm 9°$ to $\pm 29°$, during which sample structures were increasingly distorted. In comparison, with the wedge-shaped mask in Leonardo-DeStripe set to be $\pm 29°$, Fourier coefficients falling within this mask were in-painted by the graph neural network (denoted as Leonardo-DeStripe-GNN), and fully recovered after further post-processing, i.e., the complete Leonardo-DeStripe.

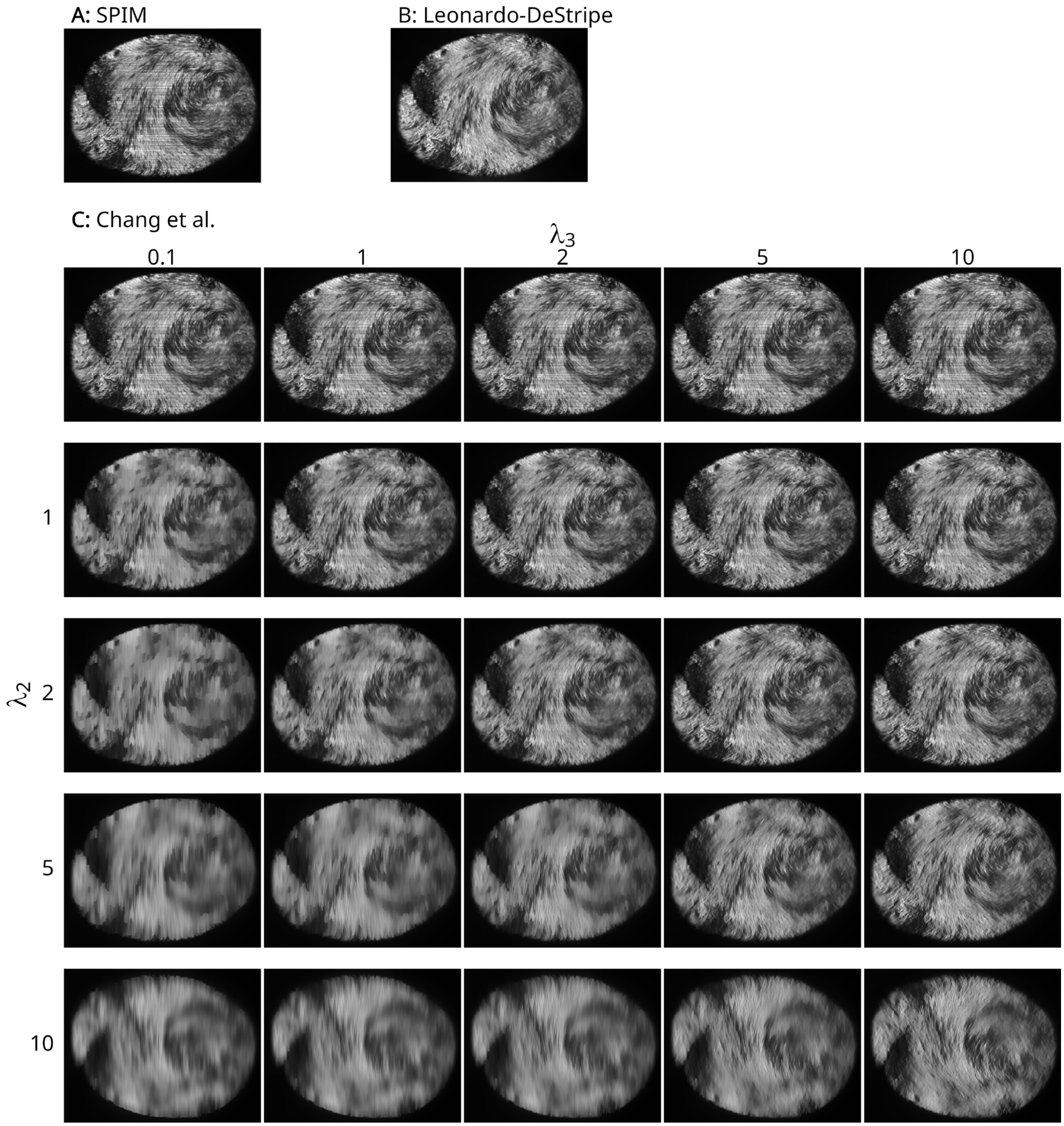

**Supplementary Fig. 3. Evaluation of the method by Chang et al. on a murine heart dataset.** (A) Original SPIM image. (B) Leonardo-DeStripe's result. (C) Outputs of the method by Chang et al., a typical optimization-based method, are given under different parameter settings (without framelet regularization). In their formulation, increasing $\lambda_2$ enhanced stripe removal, whereas increasing $\lambda_3$ helped preserve sample structures. A visually balanced result was achieved at $\lambda_2 = 5$ and $\lambda_3 = 5$. However, careful parameter tuning was required for satisfactory performance.

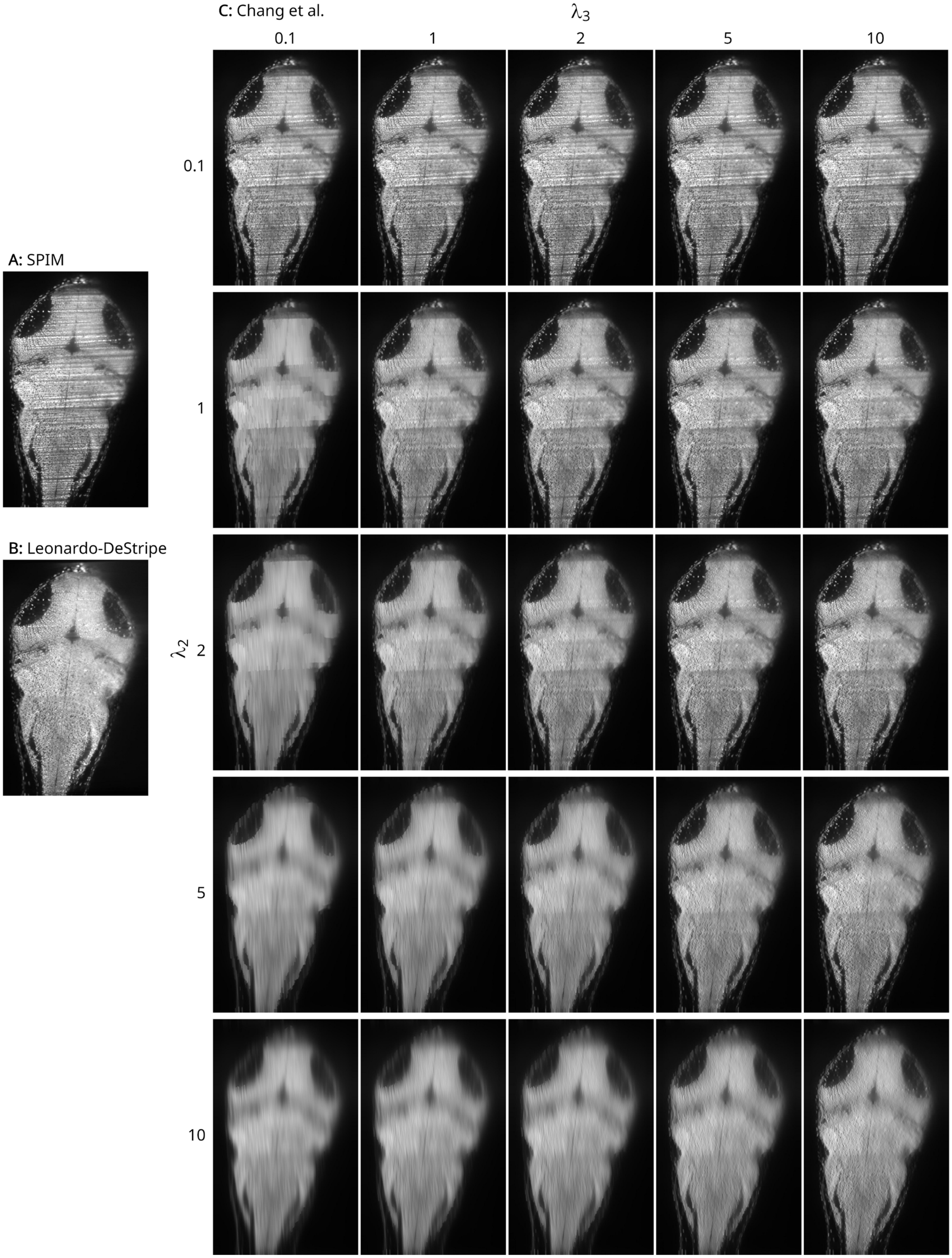

**Supplementary Fig. 4. Evaluation of Chang et al. on a zebrafish dataset.** (A) Raw SPIM image. (B)

Result of Leonardo-DeStripe. (C) Outputs of the method proposed by Chang et al., a typical optimization-based method, are given under different parameter settings (without framelet regularization). In their configuration, higher values of $\lambda_2$ led to stronger stripe removal, while larger $\lambda_3$ values better preserved structural details. However, even with extensive grid search, the method failed to yield satisfactory results across the parameter space.

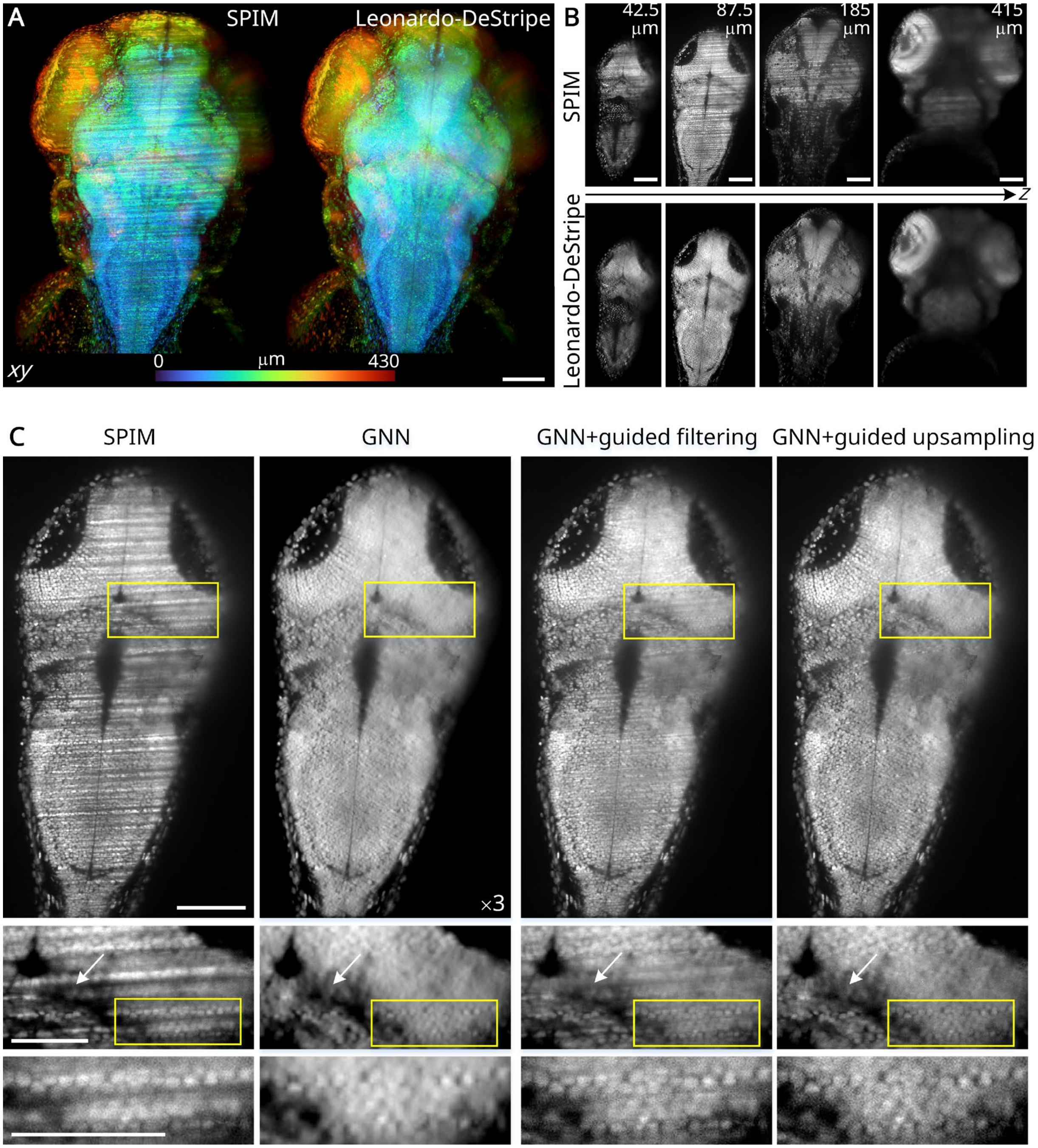

**Supplementary Fig. 5. The effect of guided upsampling in Leonardo-DeStripe.** (A) Stripes were visible in *xy* maximum projection (color-coded along *z*), whereas after Leonardo-DeStripe, they were largely resolved. (B) Four typical slices were displayed, where Leonardo-DeStripe resolved diverse stripes successfully. (C) Although the GNN in the Leonardo-DeStripe could remove the stripes completely, it took the risk of compromising sample details (GNN panel). Here, "×3" indicated that the GNN was trained in a space downsampled by a factor of 3, and its output was subsequently upsampled by a factor of 3 via bilinear interpolation for visualization alongside the other columns. When upsampling the output of the GNN and preserving sample structures, our modified guided upsampling module (GNN+guided upsampling panel) worked better than the original guided filtering (GNN+guided filtering panel), where stripes were better suppressed (white arrows). Scale bars: 100 μm (A, B, first row in C), 50 μm (last two rows in C).

125

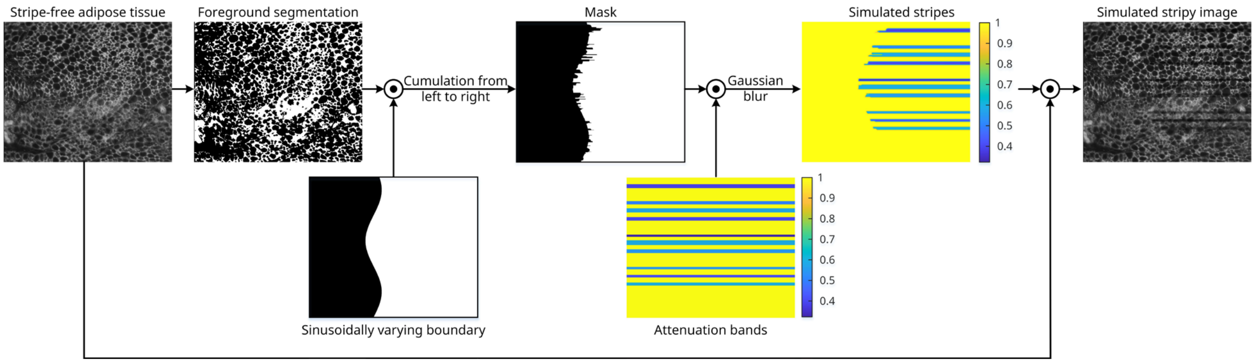

126

127 **Supplementary Fig. 6.** Simulation of stripe artifacts in SPIM. Adipose tissue was used as a stripe-free
128 ground truth image. Structured noise was simulated to mimic horizontal and multiplicative stripes.

129

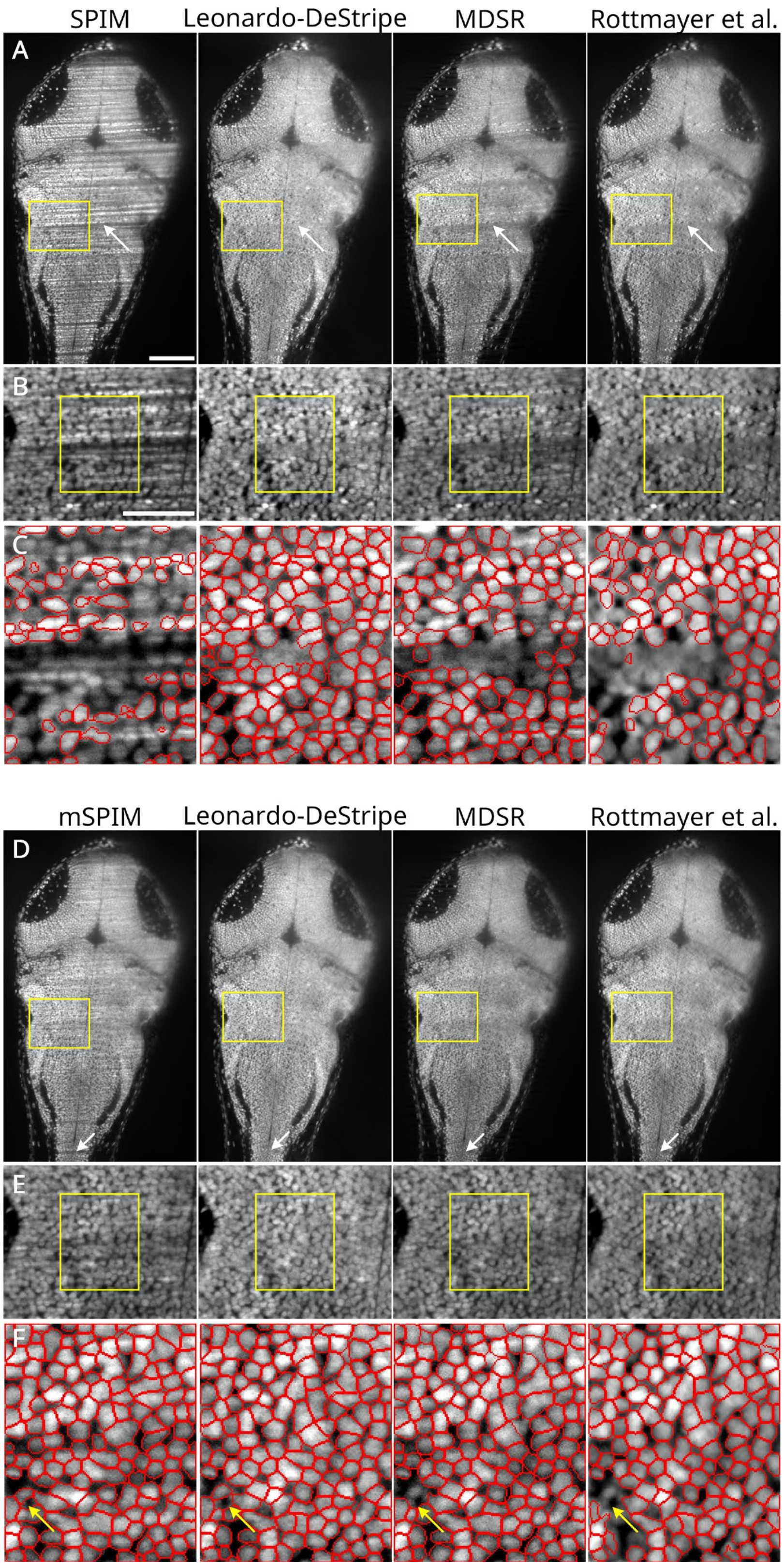

**Supplementary Fig. 7. An extended version of Fig. 2B-D in the main text with a full comparison including Leonardo-DeStripe, MDSR, and the method by Rottmayer et al.** On an H2B-GFP zebrafish,

Leonardo eliminated stripes more completely (white arrows in (A) and (D)), especially on data that was heavily affected by stripes in (B) and (E). (C) Stripe removal helped downstream processing, e.g., cell segmentation using Cellpose. Cellpose segmentation after Leonardo-DeStripe produced more continuous cell boundaries compared to MDSR-enhanced or Rottmayer et al.-enhanced data, whereas segmentation largely failed on the unprocessed SPIM image. Notably, Rottmayer et al. tended to remove sample structures. As a result, Cellpose segmented fewer cells based on Rottmayer's result than based on the original mSPIM data (yellow arrows in (F)). Scale bars are shared with Fig. 2 in the main text.

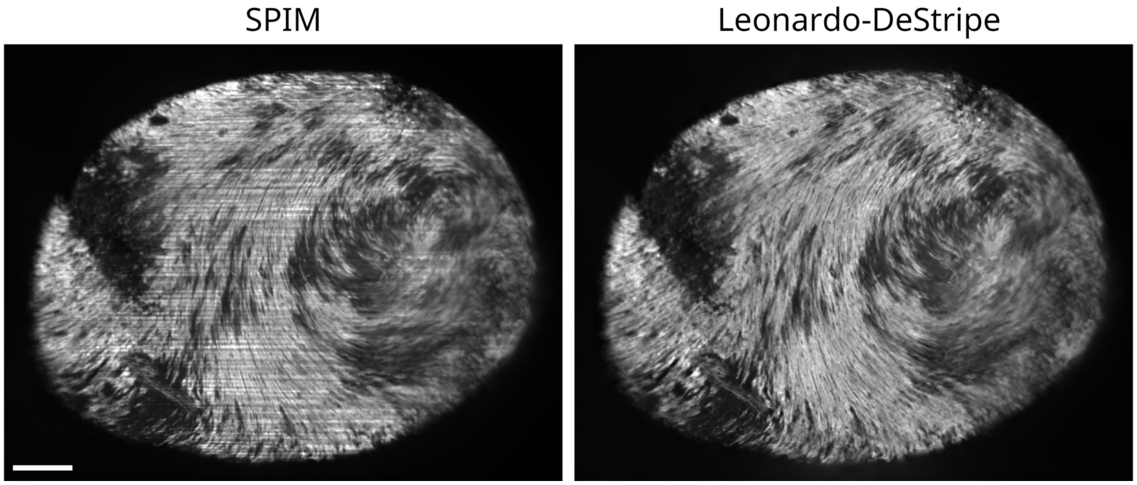

**Supplementary Fig. 8.** Leonardo-DeStripe was applied to murine heart tissue. Periodic stripe artifacts were effectively removed. Scale bar: 200 μm.

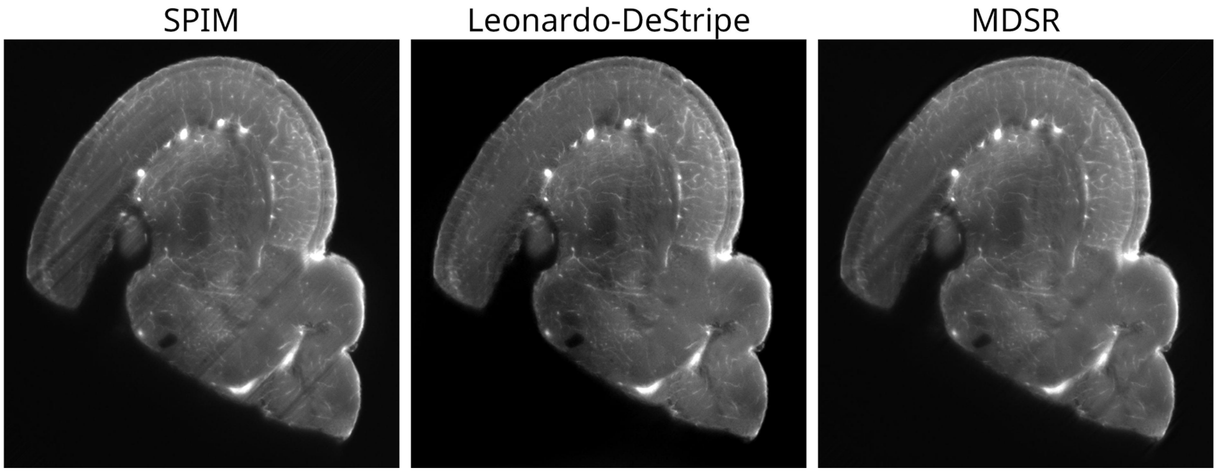

**Supplementary Fig. 9.** Removing stripes in an arbitrary direction using Leonardo-DeStripe on a zebrafish brain dataset. The stripes in the input dataset were originally horizontal (Fig. 2E in the main text). We manually rotated the stack $45°$ to test the extensibility of Leonardo-DeStripe.

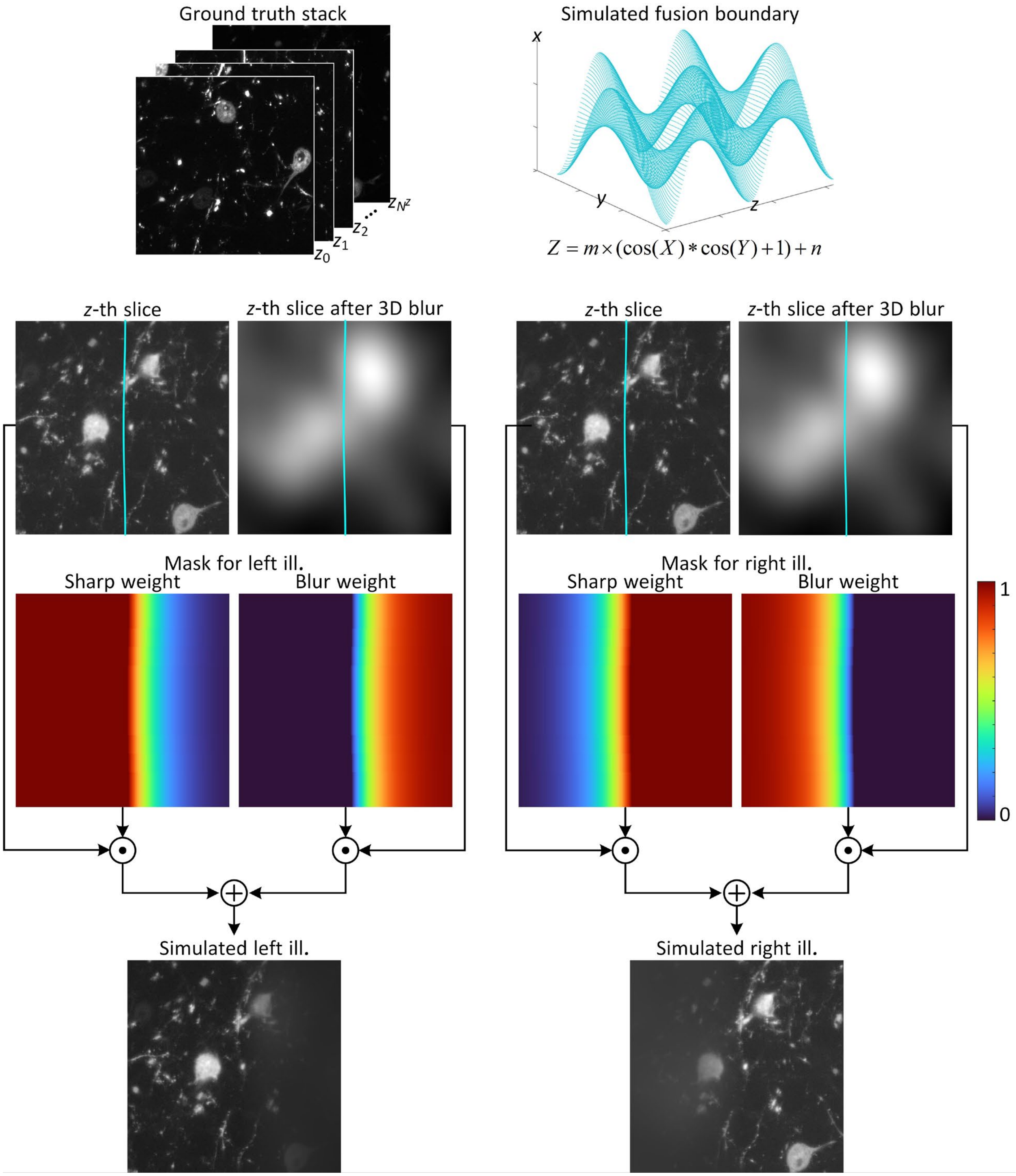

**Supplementary Fig. 10.** Simulation of two partially degraded SPIM volumes. Deteriorated stacks were simulated to become more degraded as they moved farther from the illumination sources.

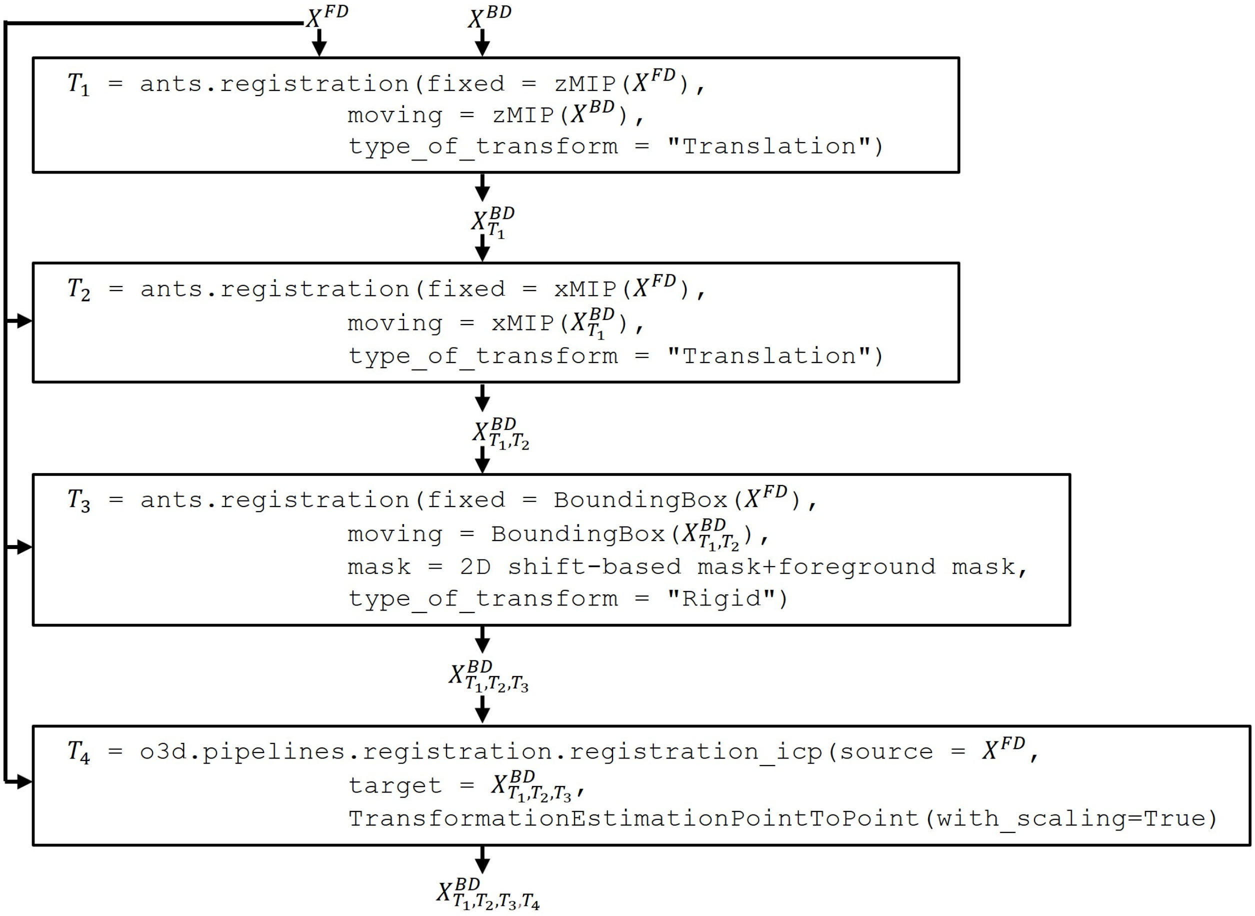

**Supplementary Fig. 11.** An illustration of the registration workflow in Leonardo, which is based on ANTsPy (ants). From coarse to fine, Leonardo registers input datasets using their maximum intensity projections (along the *z*-axis, i.e., zMIP, or along the *x*-axis, i.e., xMIP), 3D downsampled and cropped volume stacks (BoundingBox), and extracted point anchors at full resolution, step by step.

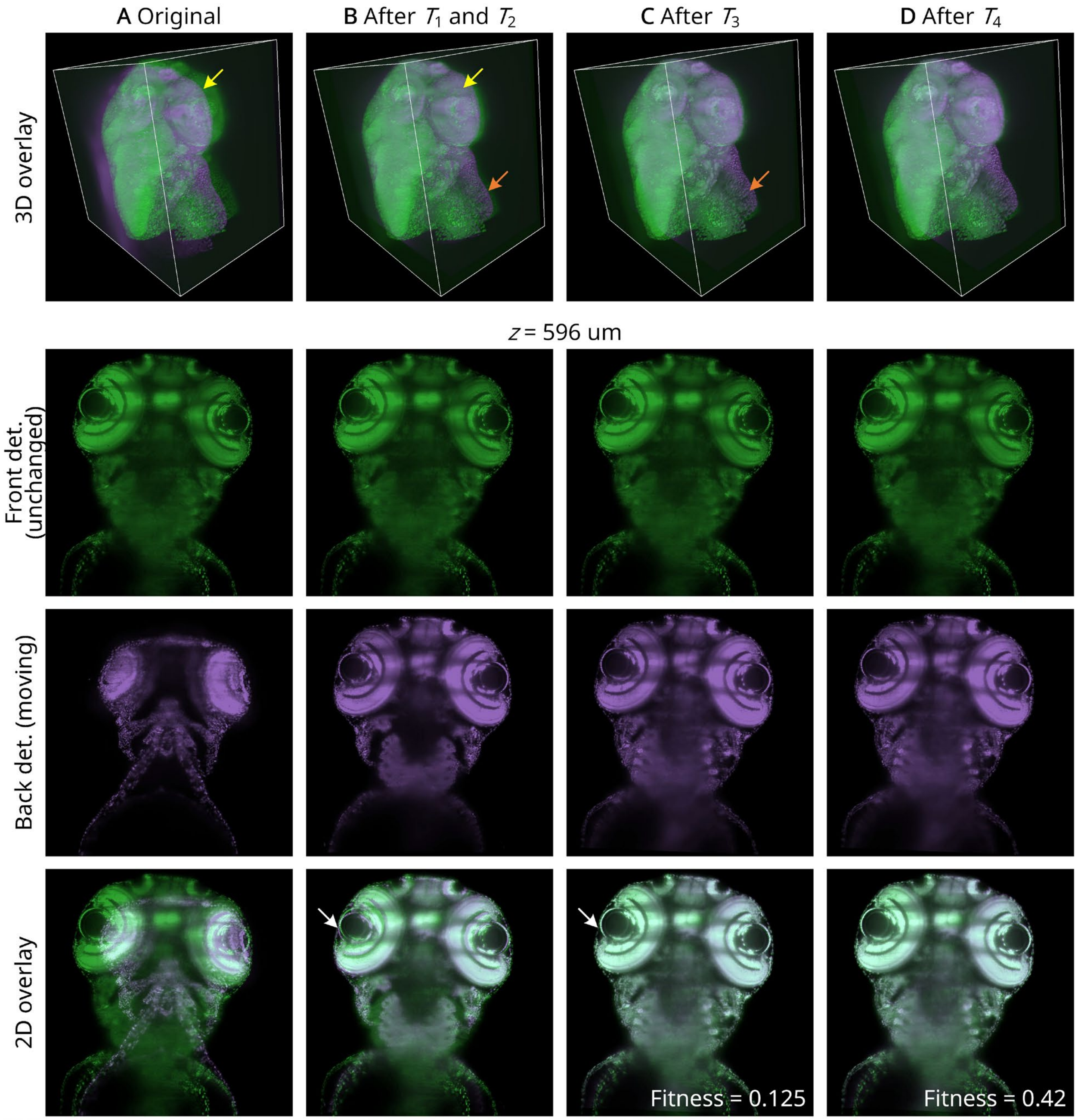

**Supplementary Fig. 12.** Registration performance from coarse to fine. The stack acquired using the camera in the back was gradually aligned with the volume captured with the camera in the front (yellow and orange arrows in the 3D overlay, and white arrows in the 2D overlay at a depth of 596 μm).

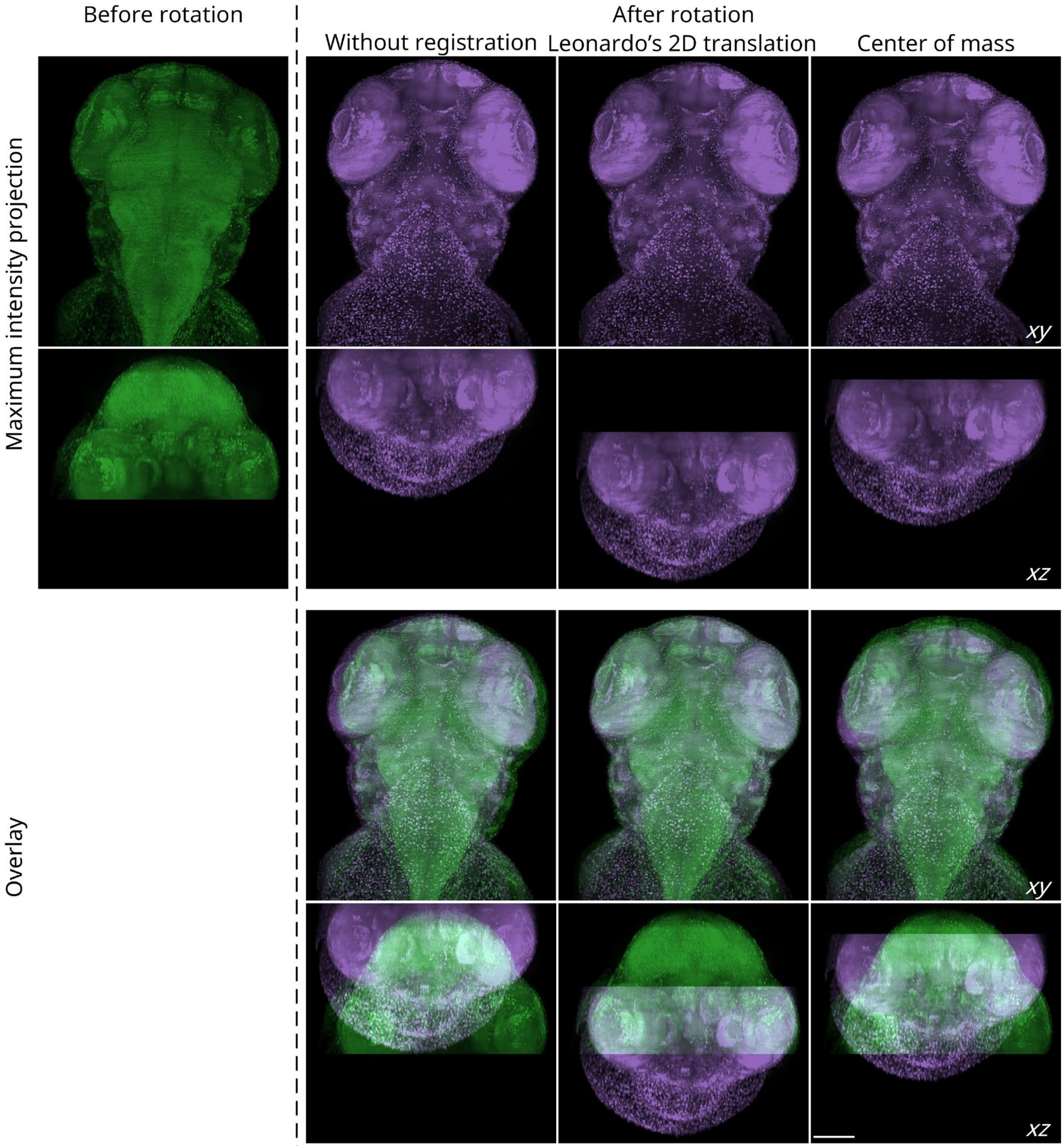

**Supplementary Fig. 13.** A comparison between Leonardo’s 2D MIP-based initial alignment and alignment of center-of-mass. When only a portion of the zebrafish was imaged each time, registration was extremely challenging. Nevertheless, Leonardo successfully mapped the rotated dataset to the same coordinate space as the dataset before rotation, whereas center-of-mass alignment failed. Scale bar: 100 μm.

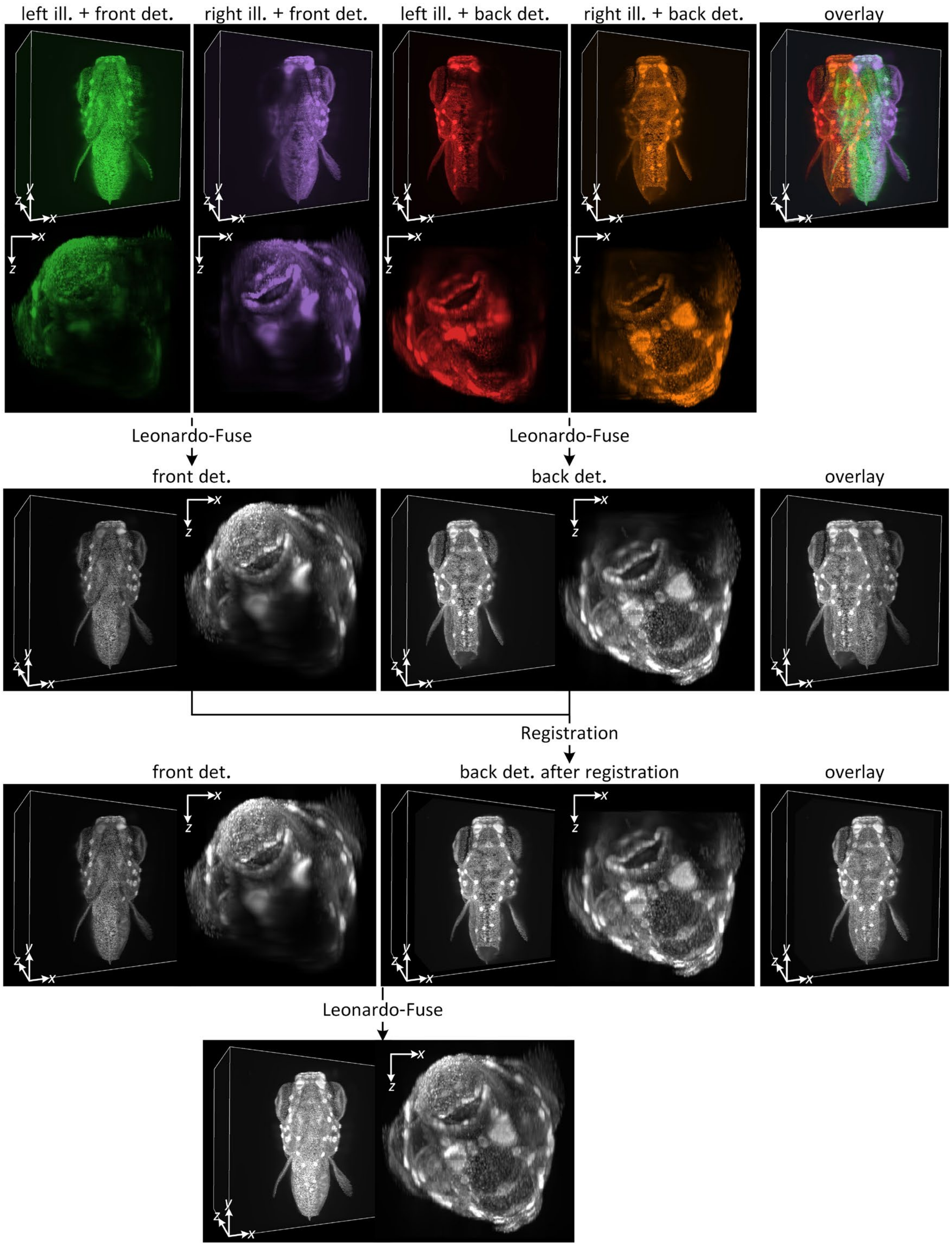

**Supplementary Fig. 14.** The complete Leonardo-Fuse workflow when registration is required on an H2B-GFP-labeled transgenic zebrafish. Leonardo-Fuse (along illumination) is first performed independently for stacks from different detection cameras. Registration is then optimized based on the fusion results along illumination. Using the registered stacks, Leonardo-Fuse (along detection) is performed as the final step.

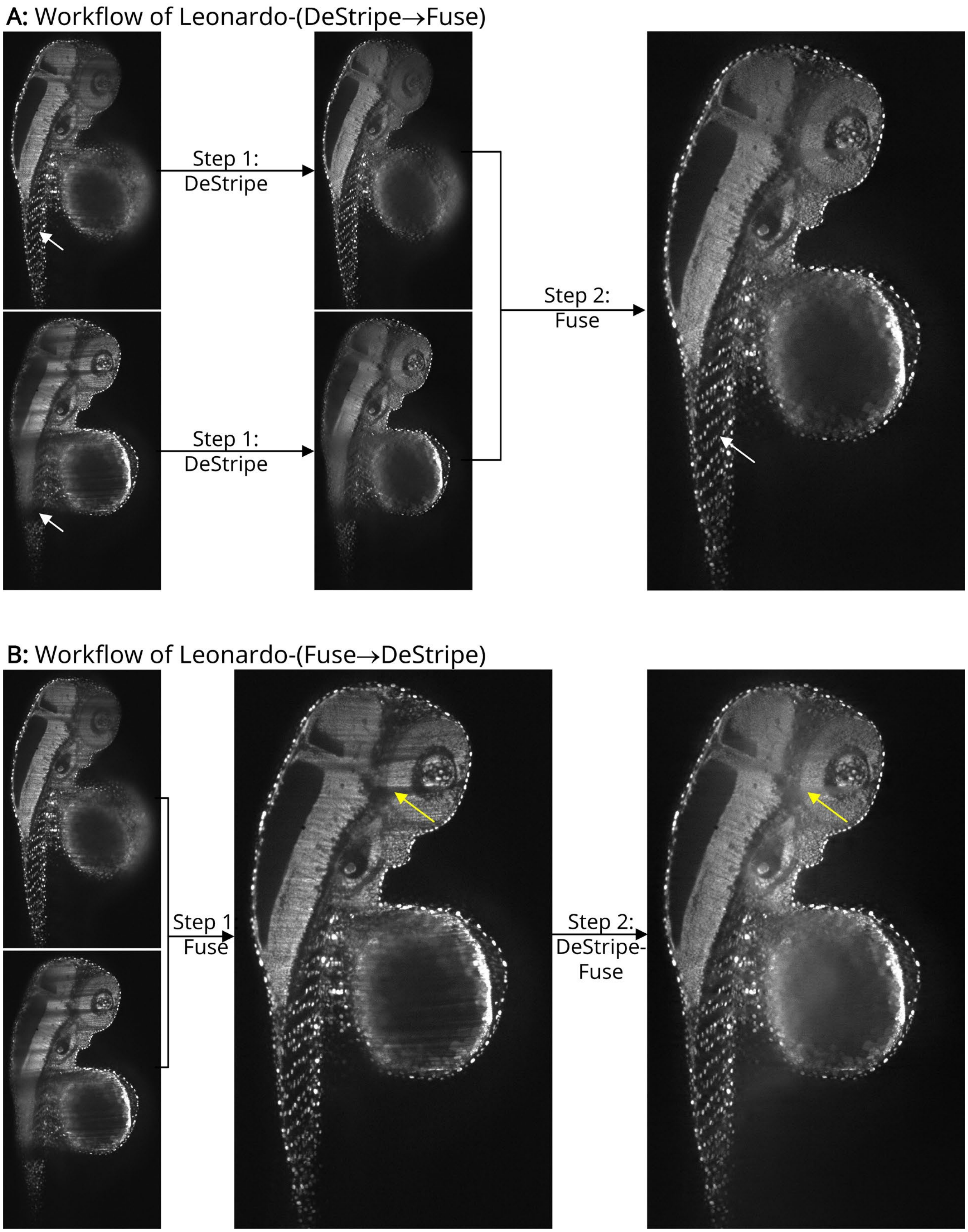

**Supplementary Fig. 15.** Two ways to formulate Leonardo workflow. (A) The first way, called Leonardo-(DeStripe→Fuse), starts with performing the DeStripe module on each dataset separately. The Fuse module is then used for dataset integration, which not only fuses stacks to maximize optical coverage but also resolves remaining extremely thick and dark stripes after DeStripe (white arrows). (B) Alternatively, we propose Leonardo-(Fuse→DeStripe), where the Fuse module is first used to remove huge and dark shadows, and Leonardo-DeStripe-Fuse, a variant of Leonardo-DeStripe, is then used to remove the remaining thin stripes (yellow arrows). Overall, Leonardo-(Fuse→DeStripe) is significantly faster than Leonardo-(DeStripe→Fuse) (see Fig. 6 in the main text).

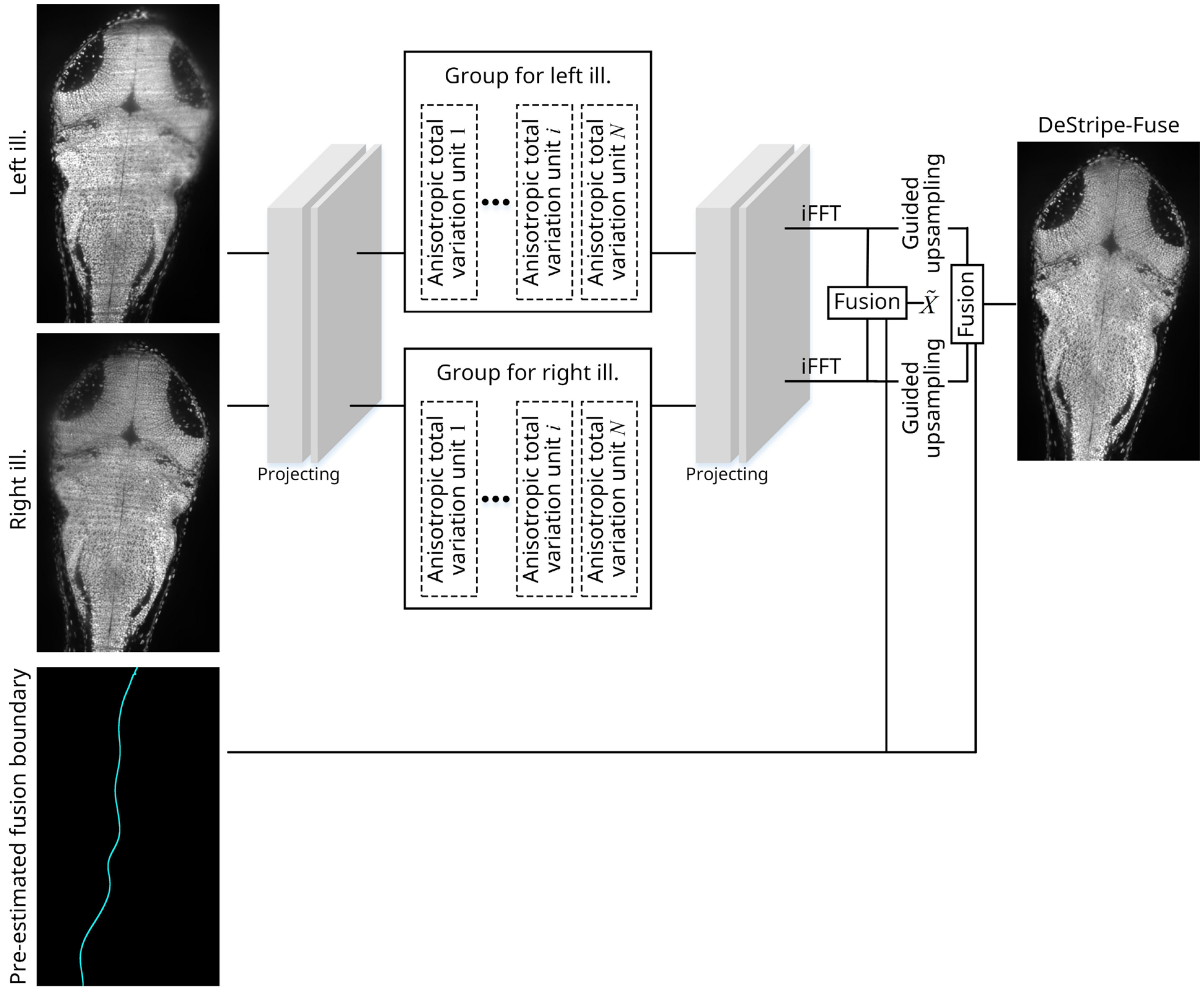

**Supplementary Fig. 16.** Workflow of Leonardo-DeStripe-Fuse. Compared to the normal Leonardo-DeStripe, Leonardo-DeStripe-Fuse can perform stripe removal for patches that are to be included in the fused result.

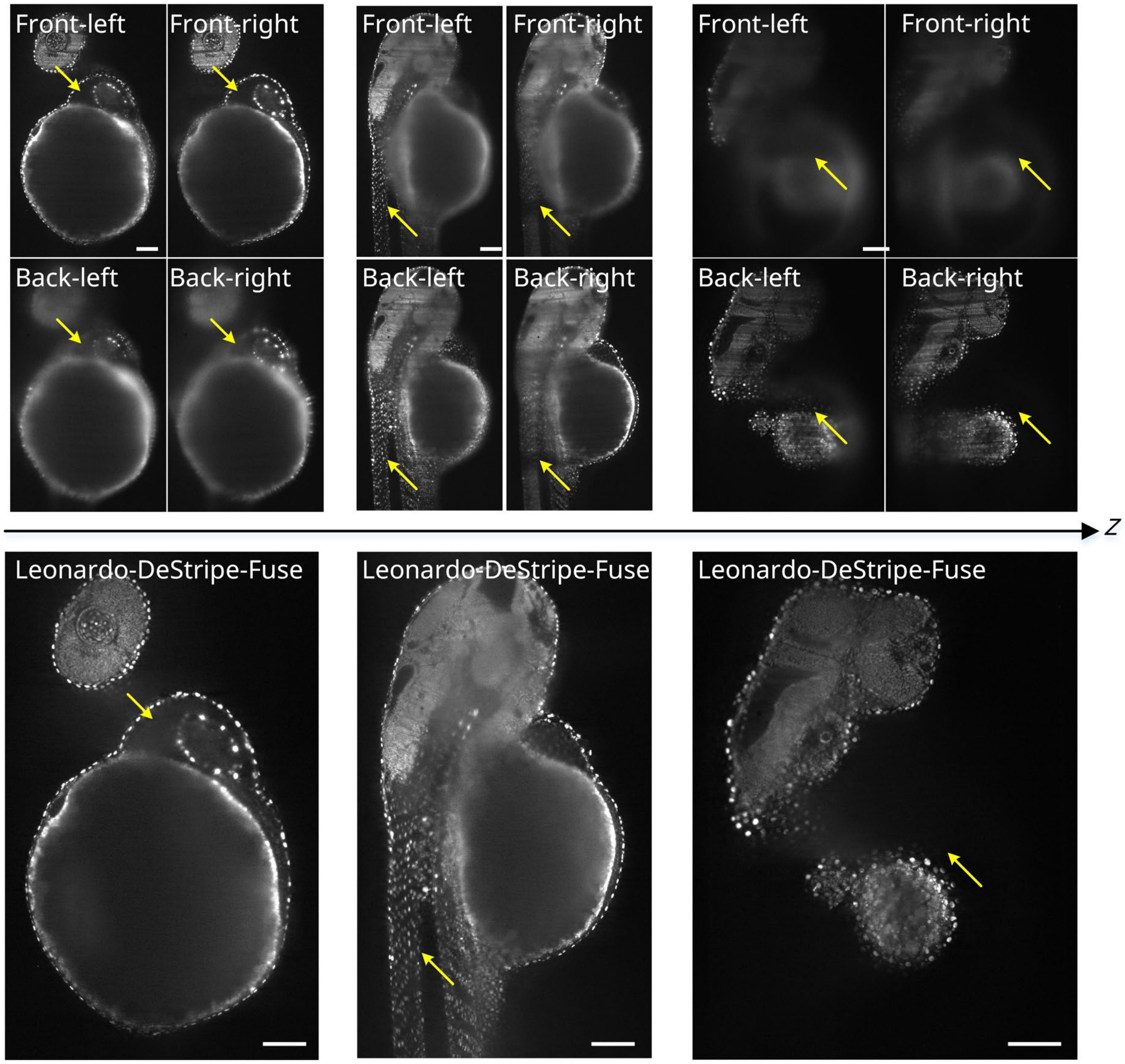

**Supplementary Fig. 17.** On a zebrafish dataset (Fig. 5A-E in the main text), Leonardo-DeStripe→Fuse restored the optimal SPIM image with minimal sample-induced artifacts, i.e., light absorption (yellow arrows in the middle panel), scattering (yellow arrows in the left panel) and refraction (yellow arrows in the right panel). Scale bars: 100 μm.

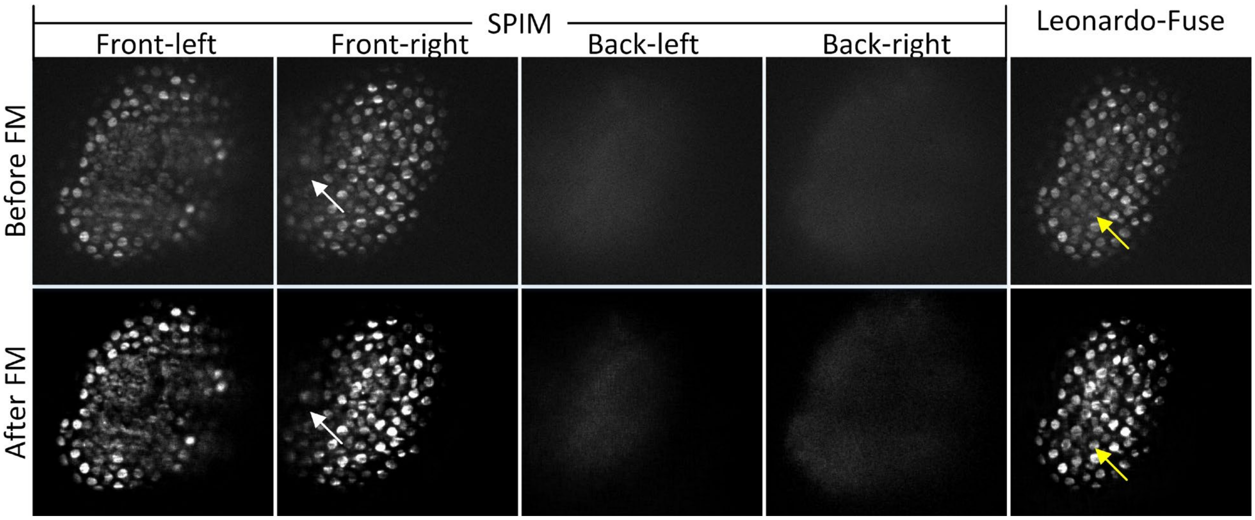

**Supplementary Fig. 18.** Restoration results using the foundation model UniFMIR, a recent universal image restoration model for fluorescence microscopy. The top row showed raw inputs before foundation model (FM) processing, including four SPIM acquisitions with single-sided illumination and detection (front-left, front-right, back-left, and back-right) and their fusion using Leonardo-Fuse. The bottom row showed the corresponding results after UniFMIR. Specifically, UniFMIR was applied to the fused volume after Leonardo-Fuse (rightmost column), and directly to the raw SPIM data for the other columns. Both the white and yellow arrows highlighted enhanced resolution and sharper nuclei following FM-based restoration.

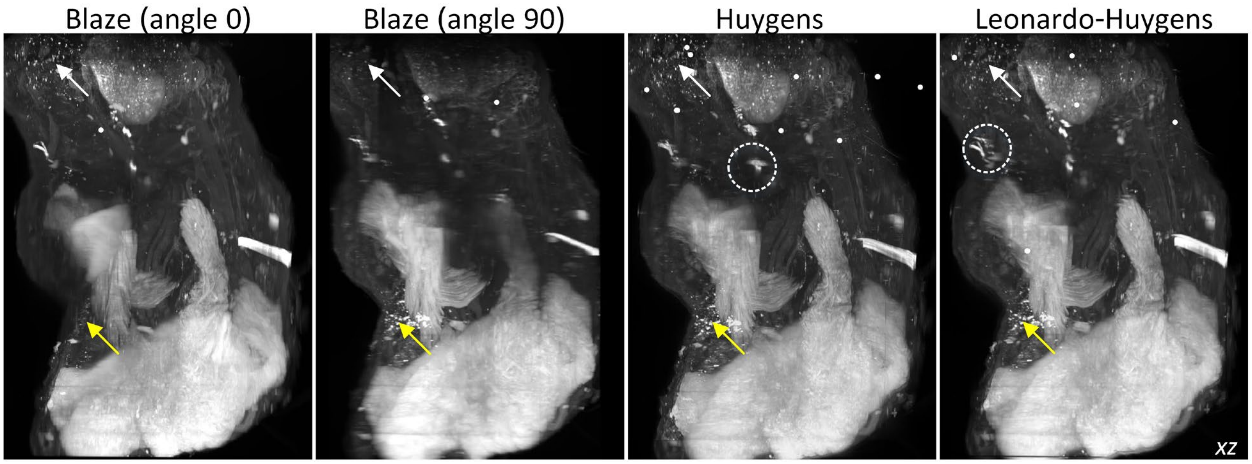

**Supplementary Fig. 19.** Multi-angular fusion results from Huygens. After applying Huygens to the fusion results from Leonardo-Fuse, sample information was further gathered (white and yellow arrows). However, misalignment existed (circled regions), as manual registration in Huygens was extremely challenging for large datasets.

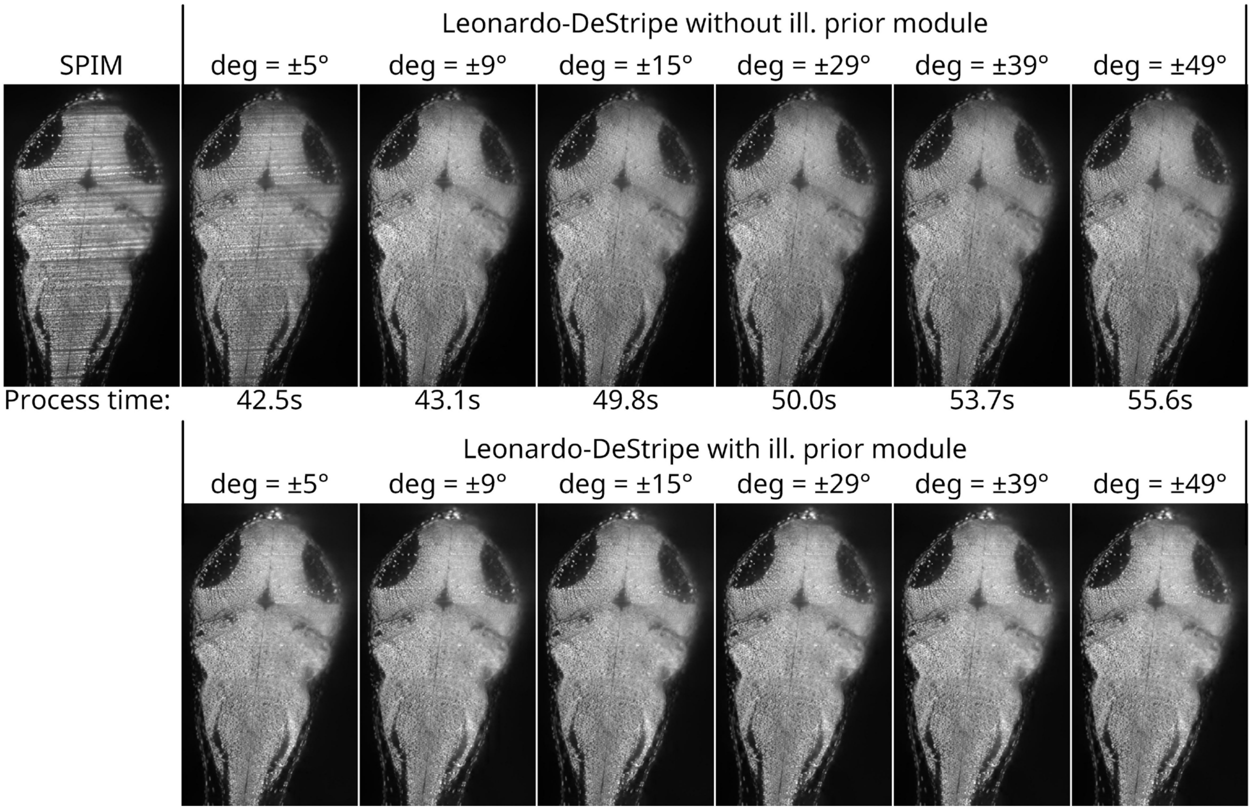

**Supplementary Fig. 20.** Impact of the angular coverage of the wedge-shaped mask in the Fourier domain in Leonardo-DeStripe. Since the mask is used by the graph neural network, we first compared the impact of different angular coverages without the post-processing module for sample structure preservation (without ill. prior, top row). As the angular range increased from $\pm 5°$ to $\pm 49°$, stripe removal became more effective, but the processing time also increased (from 42.5 seconds to 55.6 seconds). As a trade-off, Leonardo-DeStripe adopts $\pm 29°$ as the default setting, which is consistently used across all datasets in this study. Notably, the ill. prior module further improved robustness (bottom row), enabling effective stripe removal even with a narrow angular mask of $\pm 5°$.

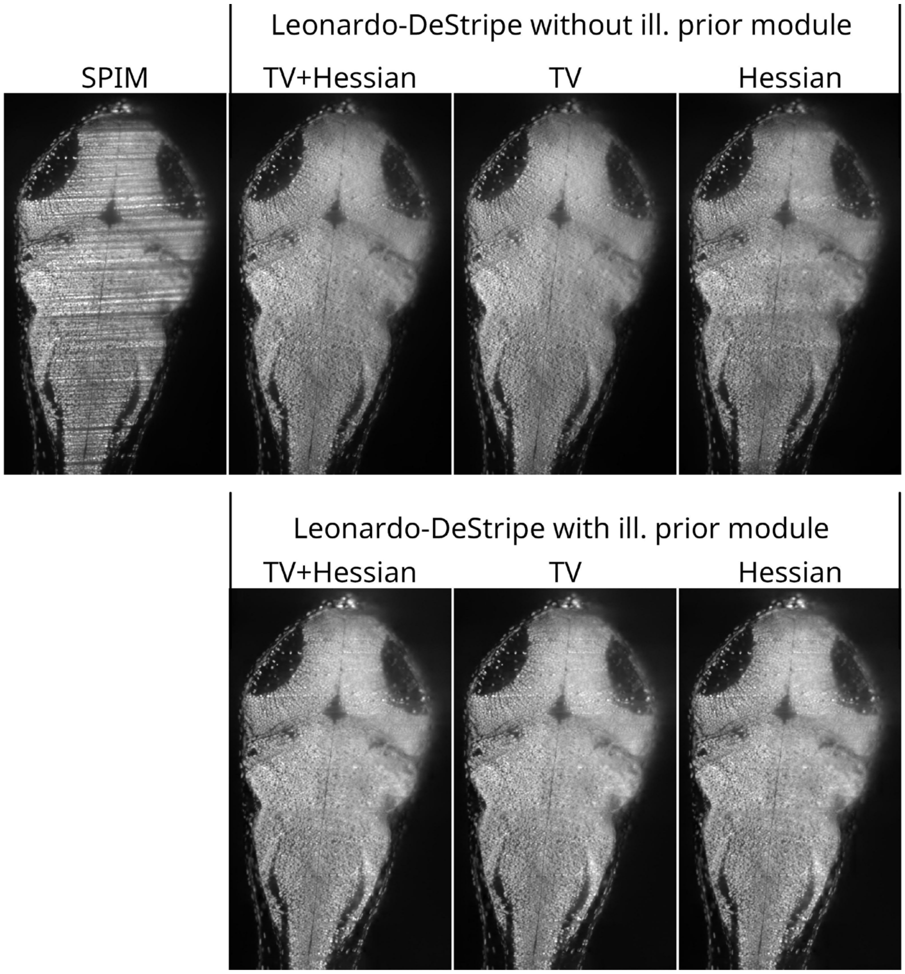

**Supplementary Fig. 21.** Impact of the regularization terms in the loss function of Leonardo-DeStripe. Since the loss function is designed to train the graph neural network, we first compared the effect of different regularization terms without the ill. prior post-processing module (top row). To better visualize the differences, the kernel size of the guided upsampling was set to 1×9, which was smaller than the default value of 49. It was evident that the TV-based loss term strongly suppressed stripe artifacts, but at the expense of removing fine sample details. In contrast, the Hessian-based loss term better preserved structural features, although some residual stripes remained. Overall, combining both TV and Hessian regularizers yielded the best performance. Notably, the ill. prior module further enhanced robustness (bottom row), enabling effective stripe removal even when only the Hessian-based regularizer was used.

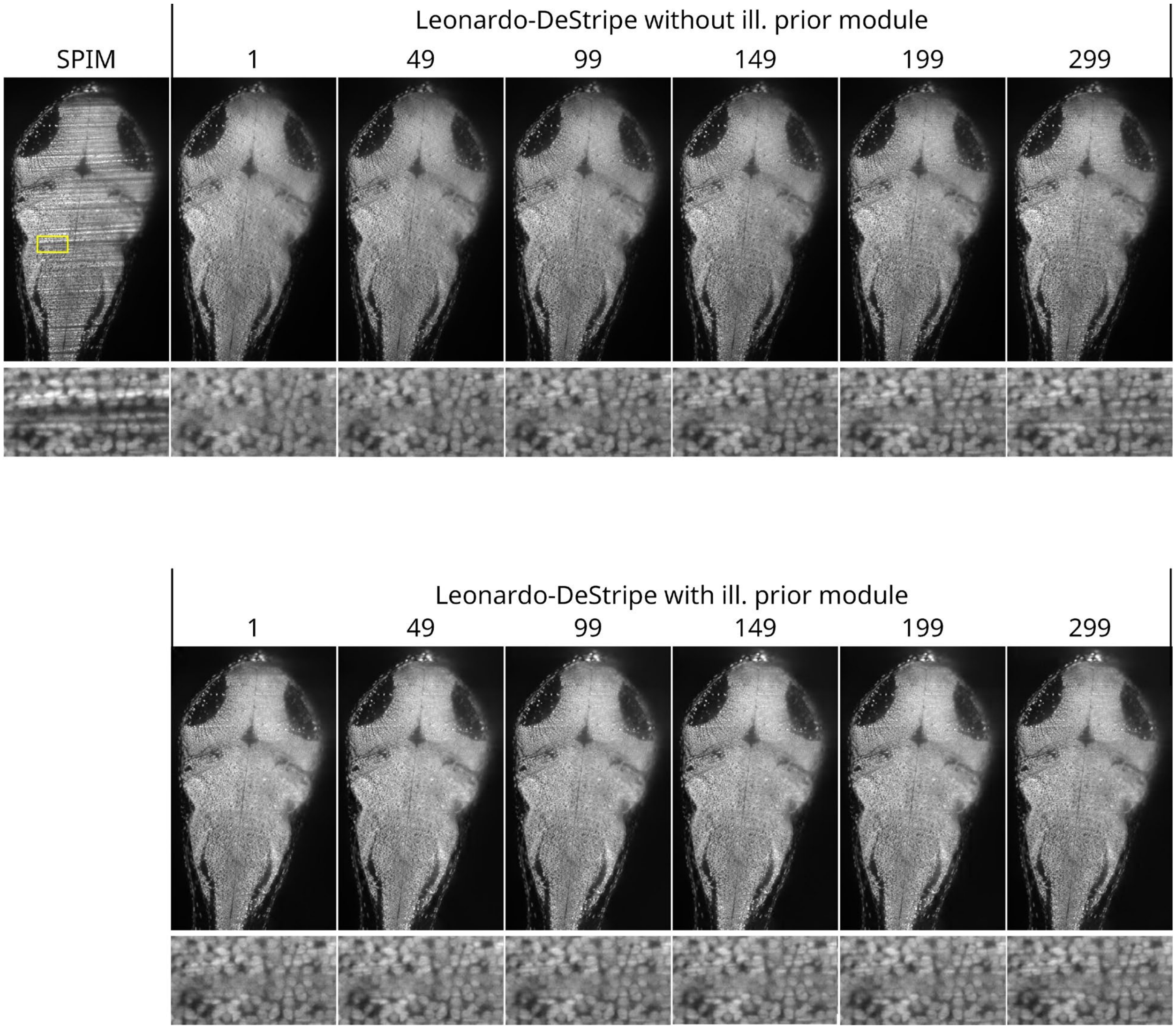

**Supplementary Fig. 22.** Impact of the window size of the guided upsampling. To better see the effect, we first applied guided upsampling with different window sizes without ill. prior module (top row). As the window size increased, sample structures could be better preserved; however, residual stripes also became more apparent (see zoomed-in region). Notably, the ill. prior module (bottom row) enhanced robustness and effectively suppressed residual stripes across all window sizes.

#### Supplementary Tables

**Supplementary Table 1**. Comparison between recent stripe removal methods and Leonardo-DeStripe

| Features | Leonardo-DeStripe | MDSR | Rottmayer et al. |
|---|---|---|---|
| Code availability | ✓ | ✓ | ✓ |
| Structure preser-vation | ✓ | x | x |
| Multi-directional stripes | ✓ | ✓ | x |
| Interactive inter-face | ✓ | x | x |

**Supplementary Table 2**. Comparison between recent image fusion methods and Leonardo-Fuse

| Features | Leonardo-Fuse | BigStitcher-Fuse | MST-SR |
|---|---|---|---|
| Code availability | ✓ | ✓ | ✓ |
| Ghost artifact removal | ✓ | x | x |
| Signal fidelity[a] | ✓ | x | x |
| Speed | Fastest | Slowest | Slower |
| Registration inside | ✓ | ✓ | x |
| Interactive interface | ✓ | ✓ | x |

[a] Signal fidelity refers to the preservation of the original signal from the high-quality side without any manipulation, rather than altering or reconstructing the original measurements.

**Supplementary Table 3**. Runtime and memory footprint comparison between baseline ANTsPy-only registration and the lightweight Leonardo pipeline.

| Methods | Speed [min] | Peak memory [GB] |
|---|---|---|
| ANTsPy-only | 17.6 | 36.8 |
| Leonardo | 4.9 | 21.3 |

**Supplementary Table 4**. Key differences between Destripe and Leonardo-DeStripe.

| Feature | DeStripe | Leonardo-DeStripe |
|---|---|---|
| Definition of corrupted Fourier coefficients | Identified via Rayleigh distribution fit; assumes noise model and may be unstable | Uses wedge-based mask; robust, model-agnostic, and direction-aware |
| Treatment of unmasked Fourier coefficients | All coefficients (including uncorrupted) are updated; slower | Only wedge-masked region is updated; faster |
| Structure-preserving regularization | Unfolded Hessian prior via Bregman iterations; iterative and costly | Lightweight anisotropic TV unit; Hessian used as loss term; effective |
| Fidelity loss (self-supervision) | Self2Self denoising with handcrafted isotropy regularizer; relies on statistical priors | Guided filtering-based loss using pseudo-target; better separates structure from artifacts |
| Training resolution and scalability | Requires training on high-resolution data; computationally intensive | Supports downsampled training with guided upsampling; faster |
| Multi-directional stripe support | Not available | Supports; handles dual-side or rotational setups |
| DC component modeling | Not available | Dedicated branch to estimate DC term; resolves mathematical gap and improves low-frequency reconstruction |
| Illumination-aware postprocessing | Not available | Postprocessing using illumination priors; improves sample structure preservation |

### Supplementary Notes

#### Supplementary Note 1: Stripe removal in SPIM

Stripe artifacts are not unique to SPIM-based imaging but are a common issue across various microscopy modalities. They can appear at multiple length scales, from large-scale volumetric imaging to high-resolution techniques such as Focused Ion Beam Scanning Electron Microscopy (FIB-SEM)[1] and Atomic Force Microscopy (AFM)[2]. In this context, computational strategies, which attempt to remove stripe artifacts after acquisition, are highly attractive.

Inspired by the desirable properties of Fourier transform in condensing stripes into isolated values in Fourier space, one line of destriping studies[3,4] suppressed stripe noises by constructing a Fourier filter on a transformed domain, e.g., wavelet[4] or contourlet domains[3]. However, filtering-based methods risked removing structural information of the sample which falls within the same filter band, resulting in undesirable image blurring[5,6].

On the contrary, another line of works treated the destriping issue as an ill-posed inverse problem in the standard image space. These approaches attempted to minimize an energy function composed of a data term—encouraging consistency between the prediction and the observed degraded image—and a regularization term to impose desirable properties on the solution. The resulting energy function could be effectively solved using optimization techniques such as split Bregman[5] or primal-dual gradient hybrid method with extrapolation of the dual variable (PDHGMp)[7]. Crucially, the effectiveness of these methods hinged on the design of an appropriate energy function that faithfully modeled the destriping problem—a nontrivial task in practice. For example, regarding the regularization term, Chang et al. imposed low-rank assumption for the stripes, which only held true when the stripes covered the entire field of view but not the case in LSFM imaging[8]. Anisotropic total variation (TV) was one of the most prevalently used priors in the stripe removal task, which leveraged the directionality of the stripe artifacts[5,7]. However, its receptive field was inherently limited, preventing it from capturing the global propagation of stripe artifacts. As a result, it might inadvertently suppress genuine sample structures that aligned with the stripe direction.

Meanwhile, for the data term, most methods adopted a pixel-wise fidelity loss—typically an L1 or L2 norm—to enforce similarity between the predicted output and the observed degraded image[5,9]. While simple and computationally efficient, these point-wise terms often failed to capture the structured nature of stripe corruption, which typically affected extended regions in a spatially coherent manner. To address this limitation, more recent methods explored adaptive or structure-aware fidelity terms[7,10], though their integration into a well-behaved energy function remained challenging.

### Supplementary Note 2: Leonardo-DeStripe

#### 2.1 Stripe remover regularized by anisotropic total variation

Before Leonardo-DeStripe, several useful stripe removers treated the destriping issue as an ill-posed inverse problem. The isotropic total variation (TV) regularized split Bregman framework, which we mentioned in the **Methods** in the main text, was a typical example:

$$\begin{aligned}&\Leftrightarrow \underset{\tilde{X}}{\operatorname{argmin}} \frac{1}{2}\left\|\tilde{X}-I_s\right\|_2^2+\lambda \mathrm{R}(\tilde{X}) \\ &\Leftrightarrow \underset{\tilde{X}}{\operatorname{argmin}} \frac{1}{2}\left\|\tilde{X}-I_s\right\|_2^2+\lambda\left\{\left\|\nabla_y \tilde{X}\right\|_1+\left\|\nabla_x\left(\tilde{X}-I_s\right)\right\|_1\right\}\end{aligned} \tag{1}$$

where we follow all the definitions provided in the main text. Since the energy function on $\tilde{X}$ is intractable, the split Bregman is adopted to convert the unconstrained minimization problem on $\tilde{X}$ into a constrained one by introducing auxiliary variables $G_x=\nabla_x \tilde{X}$ and $G_y=\nabla_y \tilde{X}$:

$$\underset{\tilde{X}}{\operatorname{argmin}} \frac{1}{2}\left\|\tilde{X}-I_s\right\|_2^2+\lambda\left\{\left\|G_y\right\|_1+\left\|G_x-\nabla_x I_s\right\|_1\right\}\ s.t.\ G_x=\nabla_x \tilde{X}, G_y=\nabla_y \tilde{X} \tag{2}$$

Subsequently, by using the Bregman iteration to weakly enforce the constraints, Eq. (2) can be further transformed into an unconstrained minimization:

$$\underset{\tilde{X}, G_x, G_y}{\operatorname{argmin}}\left\{\begin{array}{l}+\frac{1}{2}\left\|\tilde{X}-I_s\right\|_2^2+\lambda\left\{\left\|G_y\right\|_1+\left\|G_x-\nabla_x I_s\right\|_1\right\} \\ +\frac{\alpha}{2}\left\|G_x-\nabla_x \tilde{X}-B_x\right\|_2^2+\frac{\beta}{2}\left\|G_y-\nabla_y \tilde{X}-B_y\right\|_2^2\end{array}\right\} \tag{3}$$

Here, the minimizations of Eq. (3) with respect to $\tilde{X}$, $G_x$ and $G_y$ can be decoupled and further converted into three separate sub-minimization problems:

- The $G_y$-related subproblem:

$$\underset{G_y}{\operatorname{argmin}}\left\{\lambda\left\{\left\|G_y\right\|_1\right\}+\frac{\alpha}{2}\left\|G_y-\nabla_y \tilde{X}^{(k)}-B_y^{(k)}\right\|_2^2\right\} \tag{4}$$

- The $G_x$-related subproblem:

$$\underset{G_x}{\operatorname{argmin}}\left\{\lambda\left\|G_x-\nabla_x I_s\right\|_1+\frac{\beta}{2}\left\|G_x-\nabla_x \tilde{X}^{(k)}-B_x^{(k)}\right\|_2^2\right\} \tag{5}$$

  where the $G_x$- and $G_y$-related subproblems can be solved as in Eq. (6) in the main text by using the shrinkage operator.

- The $\tilde{X}$-related subproblem:

$$\underset{\tilde{X}}{\operatorname{argmin}}\left\{\frac{1}{2}\left\|\tilde{X}-I_s\right\|_2^2+\frac{\alpha}{2}\left\|G_y^{(k+1)}-\nabla_y \tilde{X}-B_y^{(k)}\right\|_2^2+\frac{\beta}{2}\left\|G_x^{(k+1)}-\nabla_x \tilde{X}-B_x^{(k)}\right\|_2^2\right\} \tag{6}$$

which is a least-squares problem equivalent to:

$$\left(I + \alpha \nabla_y^T \nabla_y + \beta \nabla_x^T \nabla_x\right)\tilde{X} = I_s + \alpha \nabla_y^T\left(G_y^{k+1} - B_y^k\right) + \beta \nabla_x^T\left(G_x^{k+1} - B_x^k\right) \tag{7}$$

which can be efficiently solved using the fast Fourier transform, as shown in Eq. (6) in the main text.

Additionally, the Bregman variables $B_x^{(k)}$ and $B_y^{(k)}$ can be updated accordingly:

$$\begin{cases} B_x^{(k+1)} = B_x^{(k)} + \left(\nabla_x \tilde{X}^{(k+1)} - G_x^{(k+1)}\right) \\ B_y^{(k+1)} = B_y^{(k)} + \left(\nabla_y \tilde{X}^{(k+1)} - G_y^{(k+1)}\right) \end{cases} \tag{8}$$

We additionally visualize the aforementioned split Bregman process during the first iteration in **Figure SN 2.1**. It is clear that the deep learning architecture in Leonardo-DeStripe (**Extended Data Fig. 2**) mimics the first iteration of split Bregman optimization, which supports the interpretability of Leonardo-DeStripe.

The only difference is that Leonardo-DeStripe, empowered by the DC branch, is able to ignore $\tilde{X}^{(0)}$, i.e., the stripe-corrupted $I_s$, when composing $\tilde{X}^{(1)}$, thus requiring only one iteration of learning to remove the stripes.

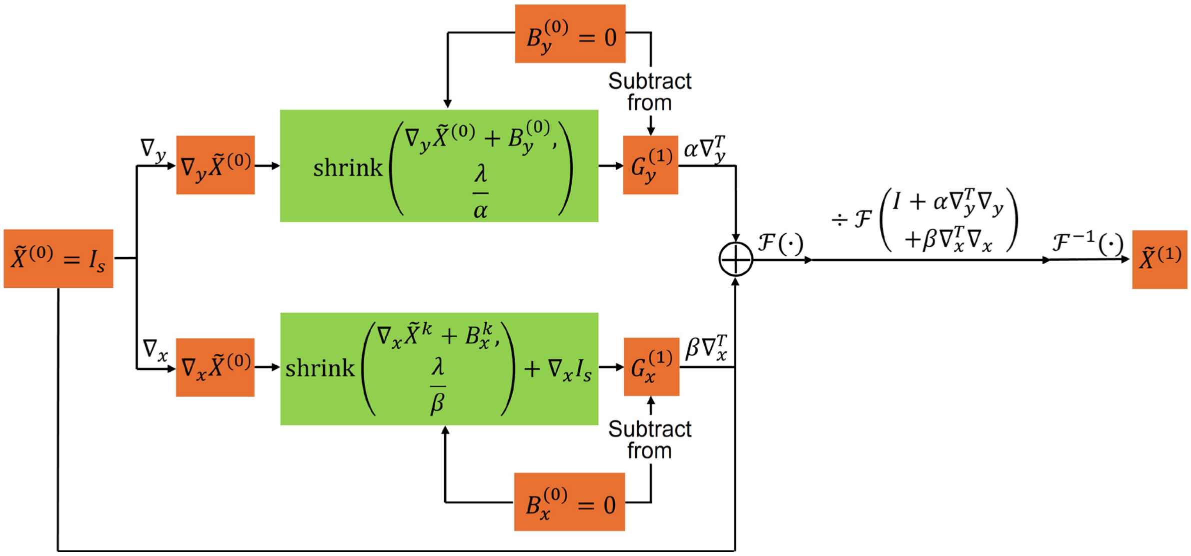

**Figure SN 2.1.** First iteration of the split Bregman framework regularized with anisotropic total variation.

#### 2.2 Improvement of Leonardo-DeStripe from a Bayesian perspective

When analyzing the aforementioned split Bregman framework for its suboptimal performance in stripe removal, we start by examining Eq. (1) from a Bayesian perspective.

Specifically, since restoring a stripe-less $\tilde{X}$ from the degraded $I_s$ is ill-posed, Eq. (1) originates from the following Maximum A Posteriori (MAP) estimation problem:

$$\tilde{X} = \underset{X}{\operatorname{argmax}}\left\{\log\big(p(I_s|X)\big) + \log(p(X))\right\}$$
$$\tilde{X} = \underset{X}{\operatorname{argmin}}\left\{\frac{1}{2\sigma^2}\|I_s - X\|^2 + \mathrm{R}(X)\right\} \quad (9)$$

where $\log\big(p(I_s|X)\big)$ is the log-likelihood of observing $I_s$ given $X$, and the log-prior $\log(p(X))$ delivers the prior of stripe-less $X$, independent of degraded $I_s$. Moreover, assuming pixel-independent Gaussian noise, $\log\big(p(I_s|X)\big)$ encourages data fidelity between the prediction $X$ and input $I_s$, whereas $\log(p(X))$ becomes the anisotropic TV prior $\mathrm{R}(X)$. In other words, the trade-off parameter $\lambda$ in Eq. (1) is equivalent to $2\sigma^2$, where $\sigma$ is the standard deviation of the Gaussian distribution assumed for the noise. However, the assumed Gaussian distribution does not hold for the stripe removal task, as $I_s - X$ represents the pixel-dependent, non-zero-mean stripe noise. To correct this, $\frac{1}{2\sigma^2}\|I_s - X\|^2$, or $\sigma$, should be pixel-variant, which is intractable in practice. Thus, Leonardo-DeStripe implicitly replaces the $\frac{1}{2\sigma^2}\|I_s - X\|^2$ with:

- Solving the $G_y$-related subproblem using a GNN, where the wedge-shaped mask draws the attention of the stripe resolver to a wedge-shaped region in the Fourier domain, perpendicular to the stripe orientation. Conceptually, the threshold $\frac{\lambda}{\alpha} = \frac{2\sigma^2}{\alpha}$ in the shrinkage operator is now spatial-variant with attention focused only on stripe-related pixels. This intuition was further supported by our ablation studies (Extended Data Fig. 1), where we observed a drastic drop in stripe removal performance when the GNN component was removed, while the removal or inclusion of the anisotropic TV unit had a relatively minor impact. This confirmed that the GNN—not the variational prior—was the key driver of effective Fourier inpainting and stripe elimination in our method.
- Composing the stripe-less $\tilde{X}$ based solely on $G_x$ and $G_y$. Hence, $\frac{1}{2\sigma^2}\|I_s - X\|^2$ is completely ignored in the $\tilde{X}$-related subproblem.

#### 2.3 Guided upsampling with various parameters

In Leonardo-DeStripe, guided filtering (GF) is used multiple times with different settings for the window size $w_k$ and the penalization parameter $\epsilon$. Here, on a murine heart specimen, we varied $w_k$ and $\epsilon$, using the striping image as both the filter input and the guidance image, to observe the effect. In **Figure SN 2.2**, from left to right, image details in the filter output were progressively removed as the penalization parameter $\epsilon$ increased, which was consistent with previous findings[11]. Moreover, due to the directionality of the stripes, $w_k$, either along column or row, empowered GF with different properties. Specifically, $w_k$ oriented against the stripes—termed column-wise GF (**Figure SN 2.2**)—could remove the stripes with a larger $\epsilon$, although fine details in the murine heart might also be oversmoothed. In comparison, row-wise GF, which operated along the stripes, did not remove the stripes even with $\epsilon = 10$. This, in turn, suggested that the learned $a_k$ and $b_k$ acted as window-based constants, thereby preserving most sample signals after row-wise GF. This explained why:

- The GF used by Leonardo-DeStripe to refine the stripe-less output $\tilde{X}$ from deep learning should be row-wise and use a small $\varepsilon$, in order to preserve sample information as much as possible.
- The GF used by Leonardo-DeStripe in the GF-based similarity loss term should be column-wise and use a large $\varepsilon$, in order to encourage the similarity between the learned stripe-less $\tilde{X}$ and the striping $I_s$ after the stripes and/or sample details have been removed.

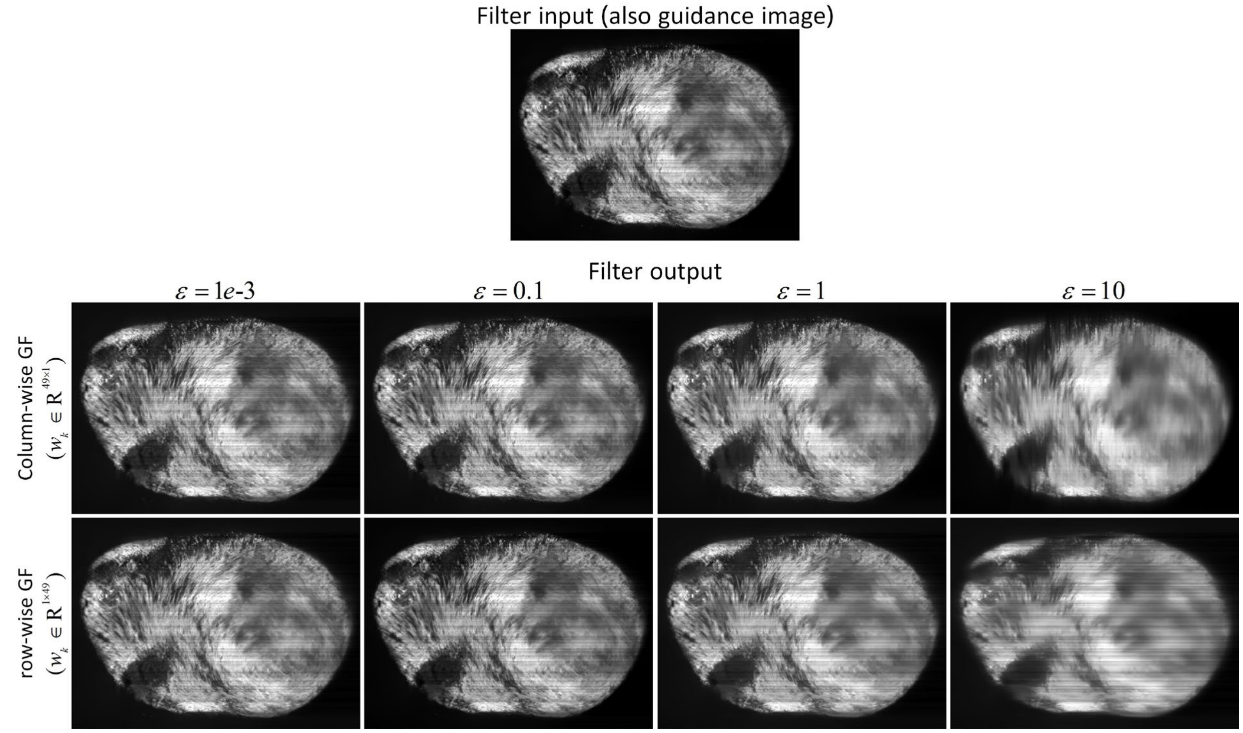

**Figure SN 2.2.** Column-wise and row-wise GFs with various $w_k$ and $\varepsilon$.

#### 2.4 Leonardo-DeStripe is rotatable for stripes in arbitrary directions

Beyond removing horizontal stripes, Leonardo-DeStripe is extensible to resolving stripes with arbitrary angular orientations, as all internal operations are rotatable. Specifically, there are four operators in Leonardo-DeStripe that need to be rotated when processing stripes oriented at an angle of $\theta$ (in radians):

- The wedge-shaped mask. The orientation of the mask is correspondingly rotated to $\theta + \pi/2$.
- First-order derivative operators including $\nabla_x$, $\nabla_y$, $\nabla_x^T$, and $\nabla_y^T$. In practice, when the orientation of the stripes is either horizontal or vertical, we use the total variation operator to extract the first-order derivative $\nabla_\perp$ and $\nabla_\parallel$ for the direction against or along the stripes, respectively:

$$\nabla_\perp = \nabla_y = \begin{bmatrix} 1 \\ -1 \end{bmatrix}, \quad \nabla_\parallel = \nabla_x = [-1,1] \tag{10}$$

  whose Fourier projection is given in **Figure SN 2.3A**. Since it is not rotatable, we instead use the following alternatives to extract the first-order derivative along horizontal and vertical directions:

$$\nabla_x = \begin{bmatrix} 0.3678 & 0 & -0.3678 \\ 0.6065 & 0 & -0.6065 \\ 0.3678 & 0 & -0.3678 \end{bmatrix}, \quad \nabla_y = \begin{bmatrix} 0.3678 & 0.6065 & 0.3678 \\ 0 & 0 & 0 \\ -0.3678 & -0.6065 & -0.3678 \end{bmatrix} \tag{11}$$

  whose Fourier projection is given in **Figure SN 2.3B**. Hence, the derivatives along and against the stripes can be calculated using the following operators, respectively:

$$\begin{aligned} \nabla_\perp &= \cos(-\theta\pi)\nabla_y + \sin(-\theta\pi)\nabla_x \\ \nabla_\parallel &= \cos\left(-\theta\pi + \frac{\pi}{2}\right)\nabla_y + \sin\left(-\theta\pi + \frac{\pi}{2}\right)\nabla_x \end{aligned} \tag{12}$$

  where $\nabla_\perp$ and $\nabla_\parallel$ denote the derivative operator for the direction against or along the stripes, respectively. The Fourier responses of $\nabla_\perp$ and $\nabla_\parallel$ when $\theta = \pi/4$ are given in **Figure SN 2.3B**. Hence, $\nabla_y$ and $\nabla_x$, which are used by the original Leonardo-DeStripe, should be replaced by $\nabla_\perp$ and $\nabla_\parallel$, respectively. $\nabla_y^T$, and $\nabla_x^T$ can be replaced by the inverse of $\nabla_\perp$ and $\nabla_\parallel$, correspondingly.
- Second-order derivative operators including $\nabla_{xx}$, $\nabla_{xy}$ and $\nabla_{yy}$. When dealing with stripes along vertical or horizontal directions, $\nabla_{xx}$, $\nabla_{xy}$ and $\nabla_{yy}$, which calculate the second-order derivatives of the input images, are defined using Gaussian Hessian kernels:

$$\begin{aligned} \nabla_{xx} &= \frac{x^2-\sigma^2}{2\pi\sigma^3}\exp\left(-\frac{x^2+y^2}{2\sigma^2}\right) \\ \nabla_{xy} &= \frac{xy}{2\pi\sigma^6}\exp\left(-\frac{x^2+y^2}{2\sigma^2}\right) \\ \nabla_{yy} &= \frac{y^2-\sigma^2}{2\pi\sigma^3}\exp\left(-\frac{x^2+y^2}{2\sigma^2}\right) \end{aligned} \tag{13}$$

where $\sigma$ is the standard deviation of the Gaussian distribution and can be manually set by the user (1 by default). When digitizing, the kernel size is $2 \times \mathrm{ceil}(6\sigma) + 1$ with $\mathrm{ceil}(\cdot)$ being the ceiling function. The Fourier responses of the Hessian operators when $\sigma = 1$ are given in **Figure SN 2.4A**. When stripes are oriented at degree $\theta$, the Hessian operators are rotated correspondingly:

$$\begin{aligned} \nabla_{\perp\perp} &= \mathrm{centerRotate}(\nabla_{yy}, \theta) \\ \nabla_{\perp\parallel} &= \mathrm{centerRotate}(\nabla_{xy}, \theta) \\ \nabla_{\parallel\parallel} &= \mathrm{centerRotate}(\nabla_{xx}, \theta) \end{aligned} \quad (14)$$

where $\mathrm{centerRotate}(X, \theta)$ rotates matrix $X$ by $\theta$ degrees, with the origin of rotation at $[\mathrm{ceil}(6\sigma), \mathrm{ceil}(6\sigma)]$. The Fourier responses of the Hessian operators $\nabla_{\perp\perp}$, $\nabla_{\perp\parallel}$, and $\nabla_{\parallel\parallel}$ when $\sigma = 1$ and $\theta = \pi/3$ are given in **Figure SN 2.4B**.

- Window $w_k$ in GFs. The original $w_k$ used by row-wise and column-wise GFs, respectively, is given in **Figure SN 2.5A**. When dealing with stripes oriented at $\theta$, $w_k$ should be rotated to along with the stripes, i.e., $\theta$, for the row-wise guided upsampling used in refining the neural network output. In comparison, the column-wise GF used in the GF-based similarity loss term should be rotated to oppose the stripes, that is $\theta + \pi/2$. The rotation of the window is implemented with `ndimage.rotate` function provided by `Scipy` package in Python. $w_k$ used to resolve stripes with $\theta = \pi/3$ is displayed in **Figure SN 2.5B**.

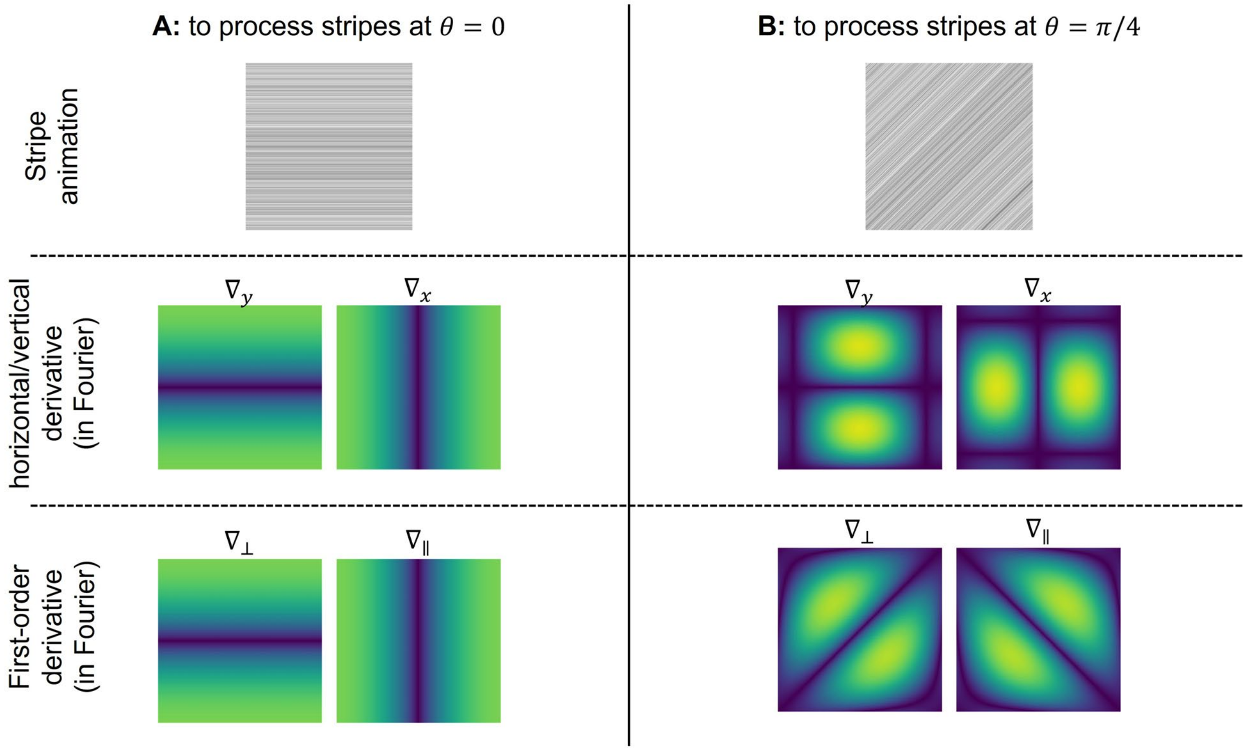

**Figure SN 2.3.** Fourier responses of first-order derivative operators $\nabla_{\perp}$ and $\nabla_{\parallel}$ used by Leonardo-DeStripe under (A) $\theta = 0$ (horizontal) stripe orientations and (B) $\theta = \pi/4$ (oblique) stripes.

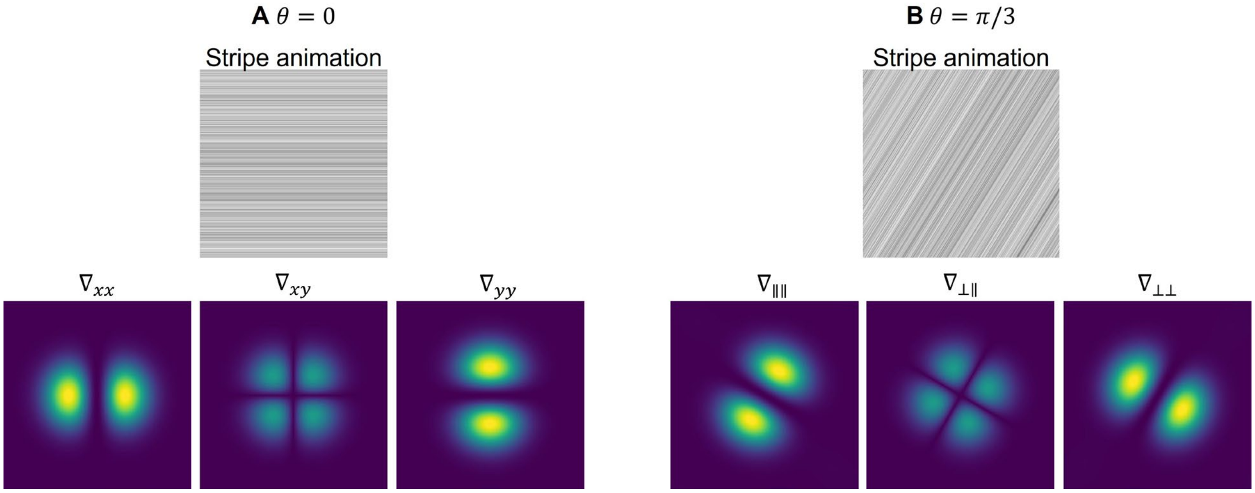

**Figure SN 2.4.** Fourier responses of Hessian operators used by Leonardo-DeStripe in different scenarios.

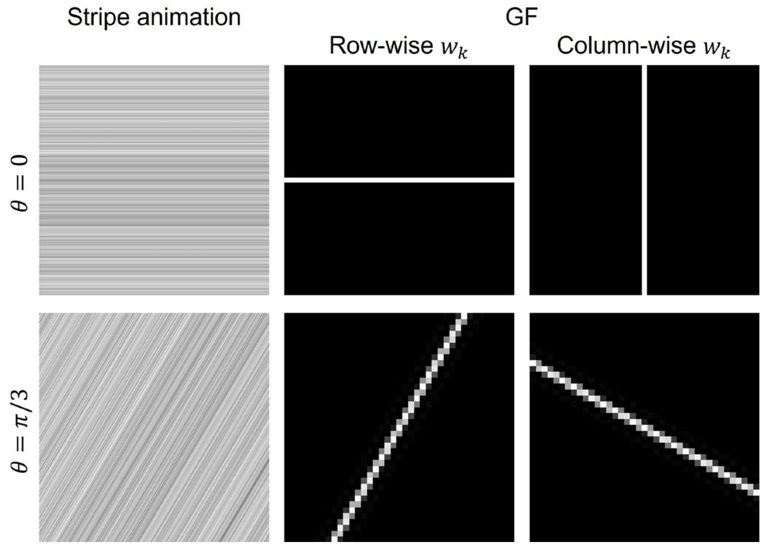

**Figure SN 2.5.** $w_k$ varies when resolving stripes with different orientations.

### Supplementary Note 3: Comparison with the previous DeStripe Method in MICCAI 2022

Before Leonardo-DeStripe, we introduced an earlier version of this approach at the International Conference on Medical Image Computing and Computer-Assisted Intervention (MICCAI) 2022[12], where we first explored the use of graph neural networks (GNNs) for stripe artifact removal in the Fourier domain. While this initial version—referred to as "DeStripe" in the following—demonstrated the feasibility of reconstructing corrupted Fourier coefficients using a GNN combined with an unfolded Hessian prior, the present work, i.e., Leonardo-DeStripe, substantially extends and improves upon that framework in terms of robustness, generalizability, and computational efficiency:

- **Definition of corrupted Fourier coefficients:** DeStripe identified corrupted coefficients by fitting a Rayleigh distribution to the Fourier magnitude histogram, which relied on a specific noise assumption and might yield inconsistent results. Leonardo-DeStripe instead uses a simple, robust wedge-based mask to define directional corruption in the Fourier domain. This model-agnostic and direction-aware approach enables more stable and controllable stripe removal across diverse samples.
- **Treatment of unmasked Fourier coefficients:** DeStripe updated all Fourier coefficients, including uncorrupted ones, which might have altered reliable frequency content and compromised structural fidelity. Instead, Leonardo-DeStripe restricts GNN updates to the wedge-masked region. While all coefficients are embedded into a shared feature space, only the corrupted region is modified by the network. This preserves uncorrupted components and reduces computational cost.
- **Structure-preserving regularization**: DeStripe used an unfolded Hessian prior via deep unfolding with Bregman iterations to preserve structure, but this increased inference time and complexity. Leonardo-DeStripe replaces it with a lightweight anisotropic total variation (TV) unit that constrains gradients along and across the stripe direction. The Hessian prior is retained as a loss term, avoiding unfolding and achieving comparable structure preservation with lower computational cost.
- **Fidelity loss (self-supervision):** DeStripe used a Self2Self denoising loss with a handcrafted Fourier-domain isotropy regularizer, relying on statistical priors. Leonardo-DeStripe instead uses a guided filtering-based loss with a downsampled, smoothed reference as pseudo-target.
- **Training resolution and scalability:** DeStripe lacked a guided upsampling mechanism and had to operate directly on high-resolution data, limiting scalability. Leonardo-DeStripe, instead, supports downsampled training via a guided upsampling module that restores corrections to the original resolution, enabling faster, memory-efficient training without sacrificing performance.
- **Support for multi-directional stripes:** DeStripe was primarily designed for single-directional stripe removal, whereas Leonardo-DeStripe extends the model to sup-

port multi-directional SPIM, by defining multiple wedge masks corresponding to different stripe orientations in Fourier space. This enables effective removal of complex stripe patterns from advanced imaging setups such as Ultramicroscope Blaze.

- **DC component modeling:** DeStripe did not explicitly model the DC (zero-frequency) component, which could cause biased reconstruction or brightness shifts under severe artifacts. Leonardo-DeStripe adds a dedicated MLP branch to estimate the DC term, resolving a mathematical gap and improving low-frequency fidelity and energy consistency.
- **Illumination-aware postprocessing:** DeStripe lacked postprocessing based on global priors. In Leonardo-DeStripe, illumination-direction priors, together with the global smoothness of the stripe artifacts, are used to preserve stripe-like sample structures along the light path, improving artifact removal in challenging cases while preserving fine details.

A comprehensive side-by-side comparison is provided in **Supplementary Table 4**.

#### Supplementary Note 4: Simulation of striping objects

To quantitatively evaluate the performance of Leonardo-DeStripe, we used a stripe-free stack of adipose tissue as ground truth and simulated stripe-shaped shadows on it (**Supplementary Fig. 6**). Specifically, attenuation bands were randomly generated with widths varying between 20–60 pixels and attenuation factors uniformly sampled from 0.3 to 0.6, mimicking the sparsity of absorbing obstacles. In **Supplementary Fig. 6**, we assumed illumination originated from the left-hand side of the image space. To ensure that simulated stripes originated from shadow-casting obstructions, the attenuation bands were masked. We performed a simple foreground segmentation using OTSU thresholding. Furthermore, to introduce spatial heterogeneity, a sinusoidally varying boundary was applied such that stripes commenced at the first foreground pixel behind the boundary and propagated across the rest of the image. The resulting stripe field was smoothed with a Gaussian filter ($\sigma = 5$) and multiplied with the input slice to yield the final corrupted image. It was noteworthy that this simulation remained a simplification compared to real experimental conditions, as phenomena such as stripe divergence due to scattering are not modeled.

#### Supplementary Note 5: Simulation of SPIM datasets with sequential dual-sided illumination

To quantify the performance of Leonardo-Fuse, specifically its along-illumination branch, in restoring a high-quality image stack by fusing two SPIM datasets illuminated via opposite lenses, we simulated two partially degraded stacks using a previously published PEGASOS-cleared mouse brain labeled with THY1-eGFP (158×512×512 voxels)[13]. Since the specimen had been well cleared in advance, it was treated as an almost optimal ground truth (GT), where the light sheet penetrated the entire brain with minimal scattering.

To simulate realistic degradation and preserve a well-defined fusion boundary for later quantification, we defined a spatially varying pseudo-fusion boundary $\omega_{z,x}^{GT}$ for each slice:

$$\omega_{z,x}^{GT} = 64\left(\cos\left(\frac{2\pi}{N_x}x - \pi\right)\cos\left(\frac{2\pi}{N_z}z - \pi\right) + 1\right) + 256 \quad (15)$$

where $x \in \{1,2,\cdots,N_x\}$, $z \in \{1,2,\cdots,N_z\}$, with $N_x = 512$, $N_z = 158$. This resulted in boundary positions within the range $\omega_{z,:}^{GT} \in [192,320]$, smoothly varying across slices.

Next, assuming illumination from the left-hand side, we modeled the output degradation by blending the original clean image $P$ with a globally blurred version $Q$ (Gaussian kernel, $\sigma = 50$), using a spatially varying weight. The weight was defined based on the horizontal distance from the simulated fusion boundary $\omega_{z,x}^{GT}$:

$$O_{z,x,y}^{left} = \alpha_{z,x,y}P_{z,x,y} + (1 - \alpha_{z,x,y})Q_{z,x,y} \quad (16)$$

where $O$ was the simulation result and the weight $\alpha_{z,x,y}$ was defined as:

$$\alpha_{z,x,y} = \frac{1}{1+\exp\left(-r(y-\omega_{z,x}^{GT})\right)} \quad (17)$$

with $r = 0.1$ controlling the steepness of the transition. This formulation ensured that regions left of the boundary (where $y < \omega_{z,x}^{GT}$) were largely preserved ($\alpha$ close to 1), while those further right were increasingly replaced by the degraded image ($\alpha$ approached 0). A symmetric strategy was used for simulating right-side illumination $O_{z,x,y}^{right}$ by flipping the sign in the sigmoid.

As a result, the obtained $O_{z,x,y}^{left}$ and $O_{z,x,y}^{right}$ mimicked the mouse brain tissue being sequentially illuminated from the left-hand and right-hand sides, respectively. During simulation experiments, results from various fusion algorithms were quantitatively compared to the ground truth $P$ (**Extended Data Fig. 8** in the main text). Additionally, the fusion boundary learned by Leonardo-Fuse (along illumination) was compared to the pseudo-fusion boundary $\omega^{GT}$. While this simulation setup might have appeared simplified—for instance, it assumed symmetrical degradation boundaries under opposite illuminations—it enabled a well-defined ground truth boundary $\omega^{GT}$, which was crucial for quantitatively evaluating both the fused image quality and Leonardo-Fuse's ability to accurately infer the fusion boundary. This trade-off allowed for more meaningful and reproducible benchmarking.

### Supplementary Note 6: Quantitative analysis

For the evaluation of either Leonardo-DeStripe or Leonardo-Fuse, we computed both the structural similarity index (SSIM) and peak signal-to-noise ratio (PSNR) slice by slice on normalized reconstruction outputs and ground truths with MATLAB (R2021a) and then reported the mean values and standard deviations in **Extended Data Figs. 4 and 8**.

The PSNR was calculated using the MATLAB function `psnr`, which computes the logarithmic ratio between the maximum possible pixel value and the mean squared error (MSE) between two images. Given a processed image $I$ (from either Leonardo-DeStripe or Leonardo-Fuse) and a reference image $K$ (either stripe-free or blur-free ground truth for Leonardo-DeStripe and Leonardo-Fuse, respectively), both of size $M \times N$, PSNR was computed as:

$$\mathrm{P}SNR = 10 \times \log_{10}\left(\frac{R^2}{\mathrm{MSE}(I,K)}\right) \tag{18}$$

where $\mathrm{MSE}(I,K) = \frac{\sum_{M,N} \| I(m,n) - K(m,n) \|^2}{M \times N}$ is the MSE between $I$ and $K$, $R$ is the maximum fluctuation in the input image data type. Since the inputs were normalized, $R = 1$.

Although PSNR is conventionally interpreted as a signal-to-noise ratio, it is in principle a rescaled representation of the mean squared error (MSE) between the reconstructed output and the reference. We reported PSNR rather than raw MSE because it provided a standardized and unit-consistent metric (in decibels), which enabled intuitive assessment of image fidelity across varying intensity distributions. In the context of stripe removal and dual-view fusion, PSNR served as a practical and interpretable indicator of reconstruction quality: higher PSNR values directly corresponded to lower pixel-wise differences from the reference image. For Leonardo-DeStripe, this reflected effective suppression of artificially simulated stripe artifacts; for Leonardo-Fuse, it reflected accurate structural integration from complementary views.

The SSIM was computed using MATLAB's `ssim` function. SSIM measures local luminance, contrast, and structure similarity between two images:

$$SSIM = \frac{(2\mu_I\mu_K + C_1)(2\sigma_{IK} + C_2)}{(\mu_I^2 + \mu_K^2 + C_1)(\sigma_I^2 + \sigma_K^2 + C_2)} \tag{19}$$

where $\mu_I$ and $\mu_K$ denote the local means of $I$ and $K$, $\sigma_I^2$ and $\sigma_K^2$ are their local variances, and $\sigma_{IK}$ is the local covariance between them. Constants $C_1$ and $C_2$ are defined as $C_1 = (P_1 L)^2$ and $C_2 = (P_2 L)^2$, where $L$ is the dynamic range of the pixel values, i.e., $1$ since our images were normalized, and $P_1$ and $P_2$ are small constants ($0.01$ and $0.03$, respectively in MATLAB).

Compared to pixel-wise metrics like PSNR, SSIM offers a perceptually motivated assessment by jointly evaluating luminance, contrast, and structural consistency between two images. This makes SSIM particularly valuable for image restoration tasks such as stripe removal and dual-view fusion, where preserving fine structural details is critical. In our evaluation, higher SSIM values indicated that the corrected or fused image more faithfully reproduced the structural content of the ground truth, even in cases where absolute intensity deviations existed. Therefore, SSIM complements PSNR by capturing perceptual and spatial coherence, providing a more holistic evaluation of the reconstruction quality achieved by Leonardo-DeStripe and Leonardo-Fuse.

### Supplementary Note 7: Information content assessment

Information content assessment was performed using a combination of the discrete cosine transform (DCT-II) and Shannon entropy to measure image quality[14]. Image patches $I \in \mathbb{R}^{M \times N}$ were first transformed into the cosine frequency domain:

$$F_{dct}(u,v) = \frac{2}{N} \sum_{i=0}^{M-1} \sum_{j=0}^{N-1} \delta(i)\delta(j) \cos\left[\frac{\pi u}{2M}(2i+1)\right] \cos\left[\frac{\pi v}{2N}(2j+1)\right] I(i,j)$$
$$\text{where } \delta(t) = \begin{cases} \frac{1}{\sqrt{2}} & \text{if } t = 0 \\ 1 & \text{otherwise} \end{cases} \qquad (20)$$

where $F_{dct} \in \mathbb{R}^{M \times N}$ is the discrete cosine transform of the patch $I$. Next, spectral entropy was used to calculate the information content of the patch $I$:

$$S_{shannon} = -\sum_{i=1}^{n} \sum_{j=1}^{n} p_{i,j} \ln\, p_{i,j} \qquad (21)$$

where $p_{i,j} = \frac{P_{i,j}}{\sum_{i,j} P_{i,j}}$, $P_{i,j} = \frac{F_{dct}(i,j)^2}{(M \times N)^2}$, $S_{shannon}$ is the Shannon entropy of the transformed image.

### Supplementary Note 8: SPIM registration

It is fundamentally crucial to fuse SPIM datasets using well-registered SPIM pairs[15]. In most commercial or scientific light sheet-based setups, for example, dual-sided illumination in SPIM and Blaze, and dual-sided detection in X-SPIM—the optical paths are typically pre-aligned. Hence, in Leonardo, algorithmic registration is only required when opposing detection views are simulated by physically rotating the specimen by 180°. However, because only complementary regions of the specimen are well imaged in different input stacks (**Fig. 1B** in the main text), registration of SPIM datasets is highly challenging. Therefore, we recommend that users of Leonardo-Fuse manually register the input stacks in advance, or consider using bead-based registration plugins[16], which require embedding fluorescent beads in the mounting medium around the specimen.

Nevertheless, Leonardo-Fuse includes an optimized 3D image-based registration workflow built upon ANTsPy[17], which can be useful when pre-registration is not available. Specifically, to maximize overlap of usable image content, registration is estimated based on the fusion result from Leonardo-Fuse (along illumination) and then applied to both image stacks captured via the back detection lens. In systems with simultaneous dual-sided illumination, such as Blaze, registration is performed directly on the raw input stacks. An illustration of the full Leonardo-Fuse workflow when registration is required is given in **Supplementary Fig. 14**.

Registration in Leonardo includes three stages. Given two fusion results, $X^{FD}$ by fusing stacks with dual-sided illumination before rotation, and $X^{BD}$ by fusing volumes with dual-sided illumination after rotation, Leonardo first aligns $X^{BD}$ into the coordinate space of $X^{FD}$ via 2D maximum intensity projection (MIP)-based translation. Specifically, the lateral shifts along the *x*-axis, namely $t_x$, and the *y*-axis, termed $t_y$, are first learned based on MIPs of $X^{FD}$ and $X^{BD}$ along the *z*-axis:

$$T_1 = \begin{bmatrix} 1 & 0 & 0 & 0 \\ 0 & 1 & 0 & t_x \\ 0 & 0 & 1 & t_y \\ 0 & 0 & 0 & 1 \end{bmatrix} \tag{22}$$

whereas the axial shift along the *z*-axis, termed $t_z$, is learned based on the MIPs of $X^{FD}$ and $X^{BD}_{T_1}$ mapped via $T_1$ along the *x*-axis:

$$T_2 = \begin{bmatrix} 1 & 0 & 0 & t_z \\ 0 & 1 & 0 & 0 \\ 0 & 0 & 1 & 0 \\ 0 & 0 & 0 & 1 \end{bmatrix} \tag{23}$$

As a result, $X^{BD}$ is mapped to $X^{BD}_{T_1,T_2}$, which is now coarsely aligned with $X^{FD}$.

Next, an affine registration is estimated to refine alignment, encompassing scaling, rotation, and translation in 3D. To this end, ANTsPy—a widely used image registration toolkit in biomedical imaging—is used. Nevertheless, one of the main drawbacks of ANTsPy is its high computational cost, when processing light sheet-based datasets at the terabyte scale. Thus, registration between $X^{FD}$ and $X^{BD}_{T_1,T_2}$ in 3D is estimated within a bounding box. The expansion of the bounding box in the *xy*-plane is defined by segmenting the MIPs of $X^{FD}$ and $X^{BD}_{T_1,T_2}$ along the *z*-axis. Meanwhile, expansion along the *z*-axis is determined such that the bounding box includes no more than $200 \times 1024 \times 1024$ pixels (empirically tested on a 50 GB random-access memory (RAM) system). Additionally, for systems with less RAM, users can define downsampling ratios—`axial_upsample` and `lateral_upsample`—for the *xy*-plane and *z*-axis, respectively.

To further accelerate the aforementioned 3D registration, masking is applied at all stages of ANTsPy's multi-resolution optimization. Specifically, in the first three resolution levels (shrink factors of 6, 4, and 2 (default setup in ANTsPy)), masks are derived from the known 2D shifts, which identify regions in each volume that lack physical correspondence due to partial sampling. Before the final full-resolution stage, foreground masks are independently generated from the fixed and warped moving volumes using OTSU method. Their intersection defines the overlapping high-intensity region, which is then used to restrict the final optimization step. This ensures that only information-rich regions are involved in the most computationally expensive step, thereby significantly improving overall efficiency while maintaining registration accuracy.

As a result, $X_{T_1,T_2}^{BD}$ is mapped to $X_{T_1,T_2,T_3}^{BD}$ after transformation using the learned affine matrix $T_3$ in the format of:

$$T_3 = \begin{bmatrix} 1 & m_z & n_z & t_z \\ m_x & 1 & n_x & t_x \\ m_y & n_y & 1 & t_y \\ 0 & 0 & 0 & 1 \end{bmatrix} \tag{24}$$

Finally, in the case of the aforementioned Affine transformation learned in a downsampled space, a descriptor-based registration can be optionally applied at full resolution. Specifically, the descriptor-based registration is estimated based on anchor points extracted using the 2D Difference of Gaussian (DoG) method, for the sake of memory efficiency. The extracted descriptors are then registered using the Iterative Closest Point (ICP) registration algorithm[18], implemented with the `Open3D` package in Python.

Overall, registration in Leonardo consists of three coarse-to-fine stages: (1) 2D MIP-based translational initialization, (2) 3D affine registration (potentially downsampled), and (3) optional full-resolution descriptor-based refinement. Knowledge of the lateral and axial resolutions of the input stacks is required throughout.

### Supplementary Note 9: Comparison between Leonardo registration and existing methods

Although Leonardo's coarse-to-fine registration is implemented using existing toolkits such as ANTsPy and Open3D, its overall performance is more reliable and robust. In particular, the initial 2D MIP-based initialization aligns volumes acquired before and after 180° rotation more effectively than standard strategies such as center-of-mass initialization.

For example, when registering zebrafish datasets in which only half of the specimen facing the detected camera is imaged before or after rotation by 180° (**Supplementary Fig. 13**), center-of-mass initialization failed due to significant differences in the imaged content. In comparison, Leonardo's MIP-based initialization successfully mapped the rotated dataset into the correct coordinate space, providing a reliable initialization for subsequent 3D registration. Without this initialization, ANTsPy failed to perform 3D registration, producing an error indicating that "the images do not sufficiently overlap."

Second, the masking strategy used in Leonardo significantly reduces memory usage and speeds up registration by limiting optimization to informative voxels. For example, on the zebrafish dataset shown in **Supplementary Fig. 13**, the use of masking led to a fourfold speed-up while requiring only half the memory compared to the baseline ANTsPy-only pipeline (**Supplementary Table 3**).

Finally, the descriptor-based registration step (optional) can leverage the affine matrix estimated from the previous stage as initialization, rather than starting from a random guess. This is particularly crucial given that ICP algorithms are notoriously sensitive to initialization. For example, when starting from a random transformation and without manual intervention (e.g., roughly aligning the datasets by hand), BigStitcher—one of the most widely used registration toolboxes in the SPIM community built upon ICP—failed and returned an identity transformation on the zebrafish shown in **Supplementary Fig. 13**.

In conclusion, compared to direct full-volume registration, Leonardo introduces a lightweight yet robust pipeline that combines 2D MIP-based initialization, bounding-box–constrained 3D affine registration, and multi-stage foreground masking. Together, this pipeline eliminates the need for manual intervention while enabling accurate and efficient alignment of partially imaged SPIM datasets.

### Supplementary Note 10: Optimized Leonardo-Fuse for large tissue

As a GPU-aided post-processing tool, Leonardo-Fuse requires a decent number of computational resources. Moreover, if registration is needed, sufficient system RAM is crucial (as previously mentioned, ANTsPy, the registration toolbox we use, is computationally demanding). Therefore, when processing extremely large specimens, a specialized version of Leonardo-Fuse is optimized, as illustrated in **Extended Data Fig. 10A** in the main text. In this version, the input volumes are first downsampled, with the sampling ratio defined by users based on the capabilities of their computing machine. Especially, this downsampling does not significantly affect the accuracy of the estimated fusion boundary, owing to the continuity of biological specimens. In comparison, the registration matrix learned in the downsampled space, if estimated, may be less accurate. Therefore, the learned registration matrix must be refined at full resolution. Considering the significant computational burden of image-based registration, this refinement is realized using only the final round of descriptor-based Affine registration which is described in the previous **Supplementary Note 9**. As a result, given the registered front-back input pair, Leonardo-Fuse can fuse them based on the upsampled fusion boundary. This follows the same strategy used to refine the fusion boundary using GF, as described in **Methods** in the main text.

#### Supplementary Note 11: Segmentation of periosteum and endosteum

To quantify the distances between periosteum (dashed colored lines in **Fig. 5H**, main text) and endosteum (solid colored lines in **Fig. 5H**, main text), we first applied OTSU thresholding to segment the foreground, followed by connected component labeling using `skimage.measure.label`. For the thin endosteum, the center point of each connected component was extracted at every *y* position, which was taken as the location of the endosteum. In contrast, the periosteum is relatively thick and solid; therefore, at each *y* position, the leftmost and rightmost boundary points of the connected component were identified, and their difference was used to quantify periosteal thickness, as shown in **Fig. 5I** (bottom panel, main text).

#### Supplementary Note 12: Leonardo-Fuse→DeStripe

Given the fusion boundary along illumination, which is pre-estimated on the stripe-corrupted data, and two input slices with opposing orientations, Leonardo-Fuse→DeStripe aims to simultaneously remove stripes in both inputs. In this process, Leonardo-DeStripe focuses exclusively on destriping regions that will be incorporated into the final fusion result. Specifically, within the deep learning framework (illustrated in **Extended Data Fig. 2C** in the main text), the two input slices (in the Fourier domain) are first mapped to a high-dimensional feature space using the same $\mathrm{MLP}^{\mathbb{C}}$ which has been discussed in **Methods** in the main text. In comparison, the subsequent anisotropic TV unit is not shared between the two opposing orientations. Instead, two separate groups of anisotropic TV units are deployed for each input in parallel. Each group may consist of multiple anisotropic TV units, with each unit dedicated to resolving stripes in one specific orientation. Thus, Leonardo-DeStripe-Fuse can be easily extended to multi-directional light sheet-based systems (e.g., UltraMicroscope Blaze). The learned feature maps from both inputs are then projected back to one-dimensional space using the same $\mathrm{MLP}^{\mathbb{C}}$. Meanwhile, two DC branches work in parallel to infer the baseline component for each input after stripe correction. The two outputs of the network, denoted as $\tilde{X}_1$ and $\tilde{X}_2$, are finally projected back to the image space using inverse FFT, and later fused into $\tilde{X}$ (following definitions used in **Methods** in the main text) using the same strategy as Leonardo-Fuse (along illumination). The remaining parts, together with the training process, of the network follow the same strategy as Leonardo-DeStripe. The only difference is that when composing full-resolution stripe-less result using guided upsampling, it is applied twice separately to $\tilde{X}_1$ and $\tilde{X}_2$, with the outputs fused again based on the fusion boundary.

### Supplementary Note 13: Parameter selection in Leonardo

In our GitHub repository, both Leonardo-DeStripe and Leonardo-Fuse modules are provided with well-tested default parameter settings, which are sufficient to produce high-quality results in most practical scenarios. Nevertheless, in certain challenging cases, fine-tuning specific parameters may further enhance performance or reduce computational cost. Here, we point out the most relevant parameters, their roles, and practical guidelines for adapting them to diverse imaging conditions. A detailed explanation of all parameters can be found in https://leonardo-toolset.readthedocs.io/en/latest/api.html.

#### 13.1 Leonardo-DeStripe: parameter discussion

Some of the key parameters that influence destriping performance—in terms of stripe removal quality and structural preservation—have already been systematically analyzed in **Supplementary Figs. 20 and 22**. Corresponding to the code space, these include the angular coverage of the wedge-shaped mask in Fourier `wedge_degree`, and the impact of the window size of the guided filtering in the guided upsampling `guided_upsample_kernel`. We next summarize additional parameters that may affect the runtime, stability, or fine control of the destriping pipeline:

- `resample_ratio` **(int, default = 3): Downsampling factor along the stripe direction used when training the graph neural network (GNN).** Since the GNN is trained separately for each slice, high-resolution training can be computationally expensive. Increasing the downsampling factor can significantly reduce training time. However, we do not recommend values greater than 5, as this may degrade the network's ability to model fine-scale stripe patterns.
- `illu_orient` **(str, optional):** Illumination orientation in the image space, which activates the optional post-processing module for sample structure preservation, i.e., ill. prior. Valid options are "top", "bottom", "left", "right", and dual-side orientations "top-bottom" or "left-right" (e.g., Ultramicroscope Blaze). Enabling this parameter may slightly increase runtime, but in some cases improve structural fidelity.
- `allow_stripe_deviation` **(bool, default = False):** A flag that controls an additional penalty term in the post-processing module for sample structure preservation. When set to True, challenging stripes (e.g., slightly tilted, wavy stripes, or illumination drift) will be more strongly suppressed. This comes at the potential cost of slightly sacrificing fine sample details. For datasets where stripes are mostly stable, keeping this parameter False to better preserve structural fidelity. Note that this option only takes effect when post-processing model is enabled via giving illu_orient.

#### 13.2 Leonardo-Fuse: parameter discussion

We are proud to say that Leonardo-Fuse is a highly robust and stable module, capable of producing reliable fusion results across a wide range of biological samples and imaging conditions without the need for extensive parameter tuning. That said, there are a few parameters that can be tuned to improve runtime efficiency, especially when processing large volumetric datasets or running on resource-limited systems:

- Common parameters that apply to all fusion tasks:
  - `resample_ratio` **(int, default = 2): Downsampling factor for estimating the fusion boundary.** A higher value (e.g., 4) reduces memory usage and speeds up computation and works well when the sample exhibits smooth or slowly varying structures.
  - `require_segmentation` **(bool, default = True): Whether segmentation is required as part of the fusion pipeline.** Segmentation is primarily used to exclude background regions and suppress ghost artifacts near sample boundaries. Thus, when ghost artifacts are not severe, segmentation can be safely disabled to reduce runtime. Also, if the data is highly sparse, no segmentation is advised.
- Dual-sided detection specific parameters, specifically if registration is required:
  - `registration_params` **(dict, optional): Dictionary controlling the behavior of the internal registration step during fusion.** If not provided, default values are used. Valid keys include:
    - `use_exist_reg` **(bool, default = False): Whether to use existing registration results instead of re-computing them.** If registration is available, for instance, in multi-channel scenario where registration is estimated on one channel and applied to the others, Leonardo-Fuse will look for `reg.npy` file under `save_folder` and apply it to the current data.
    - `require_reg_finetune` **(bool, default = True): Whether to perform additional fine-tuning registration using Open3D.** Disabling this can speed up fusion.
    - `axial_downsample` **(int, default = 1): Downsampling factor along the axial (z) direction for ANTsPy registration.** Higher values reduce memory usage but may degrade registration accuracy.
    - `lateral_downsample` **(int, default = 2): Downsampling factor along the lateral (x–y) directions for ANTsPy registration.** Setting to 1 keeps full lateral resolution during registration.
    - `skip_refine_registration` **(bool, default = False): Whether to skip**

**the refinement stage entirely under big data mode.** Primarily used when big data mode is on and registration needs to be refined at the original resolution. See https://leonardo-toolset.readthedocs.io/en/latest/fuse_det.html to check out big data mode.
